# Dentate gyrus amplifies rate-based coding while sparing global remapping across the CA3 transverse axis

**DOI:** 10.64898/2025.12.04.692471

**Authors:** Shu-Yi Hu, Ya-Li Duan, Jia-Li Long, Xing Cai, Li Lu

## Abstract

The dentate gyrus (DG) transforms overlapping experiences into discrete memory traces, yet how DG output shapes population coding within the hippocampus remains unclear. We combined selective DG lesions with large-scale recordings across the CA3 proximo-distal axis as rats explored environments varying in contextual similarity. Global remapping in CA3 remained robust following DG lesion, whereas rate-based contextual coding was substantially attenuated, most strongly in proximal CA3 and attenuating distally, paralleling mossy-fiber topography. Granule cells and mossy cells additionally coded complementary aspects of local objects and mediated CA3 responses to object manipulation. These data causally link the enhanced rate coding of proximal CA3 to its preferential DG innervation and support a dual organization of contextual discrimination: categorical changes in environmental identity are signaled independently of the DG, whereas fine-grained discrimination is facilitated by DG-mediated rate modulation. By selectively amplifying rate-coded discrimination of subtle contextual differences, the DG sharpens the hippocampal network’s capacity to resolve overlapping experiences into distinct representations.

## Introduction

The hippocampus assigns distinct representations to overlapping experiences, enabling contextual discrimination (1). In the classical view, CA3 operates as a homogeneous attractor network whose recurrent auto-associative connections support pattern completion, while the dentate gyrus (DG) decorrelates overlapping inputs via pattern separation (2). This framework has been challenged by evidence that DG and proximal CA3 (pCA3) may instead form a dedicated module for fine contextual discrimination (3): DG granule cells (GCs) fire sparsely and remap discretely across contexts, mossy cells (MCs) provide complementary, condition-specific modulation within the DG circuit (4–9), and CA3 place cells exhibit a transverse gradient of remapping, strong in pCA3 and weak in distal CA3 (dCA3) (10, 11).

Whether DG output drives this transverse organization remains untested. Lesion studies with behavioral readouts establish that the DG is necessary for contextual discrimination (12, 13), and recording in DG-lesioned animals has shown that CA3 coding of ambiguous visual scenes depends on DG input (14). However, CA3 was treated as a single structure in that study, leaving open whether DG input drives the heightened contextual sensitivity of pCA3 specifically, or whether CA3’s two coding strategies, global remapping, in which distinct cell ensembles are recruited to signal categorical changes in environmental identity, and rate-based coding, in which the population map is preserved but firing rates change systematically (15), depend on DG input differentially.

To address these gaps, we combined bilateral DG lesions with large-scale recording of CA3 place cells across the transverse axis as rats foraged in environments that varied in contextual similarity, including manipulations of local objects. Eliminating DG input substantially attenuated rate-based contextual coding in pCA3 while sparing global remapping, revealing a dual organization of contextual discrimination: categorical changes in environmental identity are signaled independently of the DG, whereas fine-grained discrimination of both non-spatial context and local objects is facilitated by DG-mediated rate modulation, with its strongest impact in pCA3. DG inputs proved most critical for local object discrimination in CA3, with GCs and MCs contributing to distinct dimensions of object features. These findings delineate how the hippocampal circuit deploys separable coding strategies under varying contextual demands.

## Results

### Recording sites and lesion profile

To determine how DG output influences contextual coding in CA3, we recorded CA3 neurons across the transverse axis in rats with selective DG lesions (LES, 7 rats) and sham-operated controls (CON, 6 rats) as they foraged in arenas where spatial, non-spatial, and local object cues were manipulated (Fig. 1A, B). In controls, tetrodes terminated in the supra- and infrapyramidal blades of the DG (23 sites) and across the entire proximodistal axis of CA3 (39 sites) (Figs. 1C; S1). In lesioned rats, all 46 tetrode tips localized to CA3 and were spatially indistinguishable from control placements (Fig. 1C, D). Recording-site distributions did not differ between groups along either the transverse or longitudinal axis (Fig. 1E; Fig. S1N; Table S1), ensuring that group differences in coding reflect loss of DG input rather than sampling bias. For analysis, CA3 sites were assigned normalized positions along the transverse axis and partitioned into proximal (pCA3), middle (mCA3), and distal (dCA3) bands (11).

**Figure 1.**
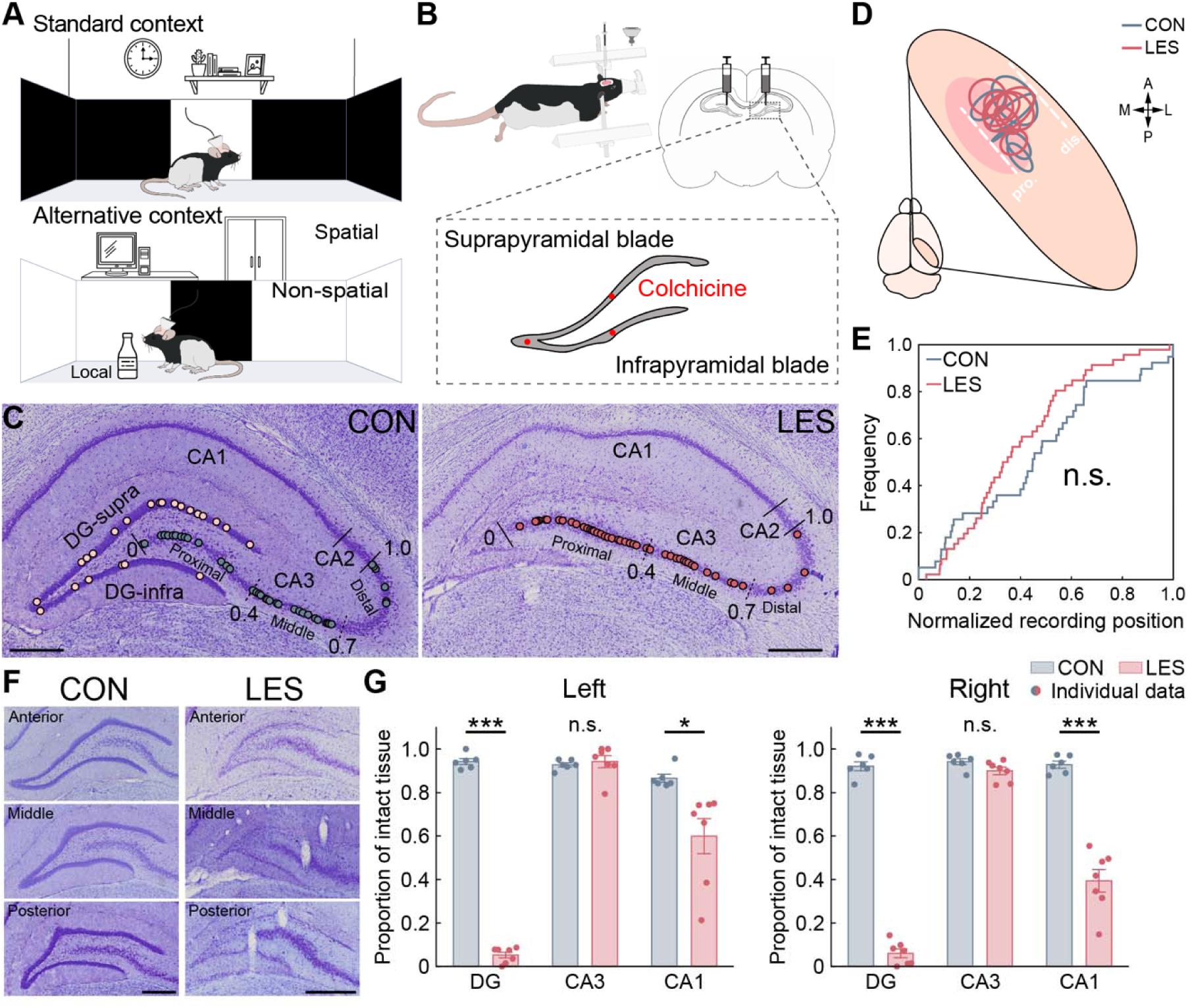
Recording sites and lesion profile. (A) Schematic of contextual manipulations. Top and bottom panels show standard and altered contexts, respectively. (B) Colchicine injection sites (red dots) for bilateral DG lesions. (C) Nissl-stained coronal sections showing hippocampal recording sites (colored dots) in control (left) and lesioned (right) animals. Solid lines, subregions; dashed lines, CA3 subdivisions. (D) Estimated CA3 recording regions for each animal. Gray circles, control (CON); red circles, lesioned (LES). Dashed lines, CA3 boundaries; pink shading, lesioned area. (E) Cumulative distribution of tetrode tip positions along the CA3 transverse axis. (F) Representative anterior-to-posterior hippocampal sections from control (left) and lesioned (right) rats, showing intact and lesioned DG. (G) Spared tissue proportions (mean ± SEM) in the left and right hippocampi; dots, individual animals. Scale bars: 100 µm (J, M); 0.5 mm (all others). n.s., not significant; *, *P* < 0.05; ***, *P* < 0.001.

Colchicine treatment produced near-total GC depletion at injection sites while sparing CA3 pyramidal layers (Fig. 1F, G). MCs and interneurons associated with the GC circuit (16) were also substantially reduced (Fig. S2A–F), and residual DG neurons in recorded rats exhibited spatially diffuse firing with low spatial information (Fig. S2G), confirming effective disruption of the DG microcircuit (Fig. S2H). CA1 overlying the injection sites was partially damaged, consistent with prior reports (12, 14, 17).

### Global remapping persists without DG input

We first examined how CA3 represents categorical environmental changes using a two-room paradigm, in which identically configured arenas were placed in two spatially distinct rooms (15). In both groups, a subset of animals (2 of 6 controls and 2 of 7 lesioned rats) exhibited only rate remapping between rooms (mean across-cell SC > 0.5, as defined by (18); Fig. S3A). We excluded these animals from global remapping analyses and report their data separately. The lesion did not alter exploratory behavior (Fig. S3B).

In control rats, room switching induced robust global remapping in both DG and CA3 (Fig. 2A). Decorrelation of spatial maps was strongest in DG and pCA3, declining progressively toward dCA3 (Fig. S3C), consistent with the established transverse organization (11). Dentate neurons were classified into GCs and MCs based on distinct activity patterns (19) (Fig. 2B, C). Between-room spatial correlations (SCs) remained low in both control and lesion groups (Fig. 2D) with comparable fractions of globally remapped neurons (76.2–87.4, all *P* > 0.55; Table S1). Equivalent decorrelation was confirmed by overlapping SC–position regressions at the tetrode level (Fig. 2E) and by matching SCs at the animal level (Fig. S3D). However, the proximo-distal gradient in the fraction of active neurons was attenuated (Fig. 2B, C), indicating disrupted dentate drive to CA3 activity. Room-switch sensitivity at both cellular (group × band interaction; Fig. 2D) and tetrode (permutation analysis of regression slopes; Fig. 2E) levels was also reduced. Rate differences (RDs) between trial pairs were unchanged (Fig. S3E–G). Population vector correlations (PVCs), computed across all neurons (Table S2), confirmed preserved room discrimination (Fig. 2F, G; S3H; Table S1). Categorical contextual changes are therefore signaled by CA3 independently of DG input, which may help to maintain the subtle proximo-distal gradient in discrimination.

**Figure 2.**
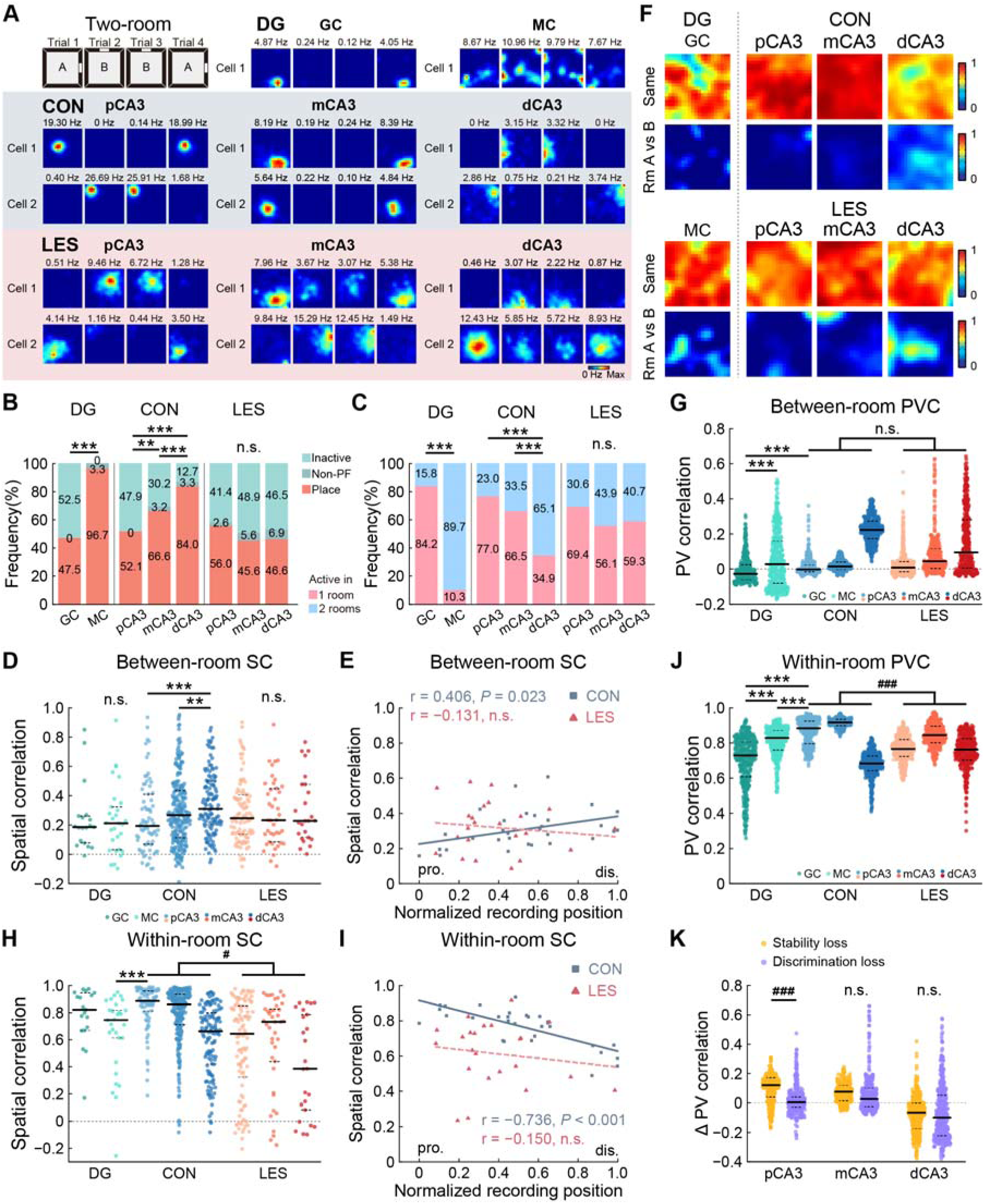
Global remapping persists without DG input. (A) Representative rate maps of DG and CA3 neurons in control (CON, top) and DG-lesioned (LES, bottom) rats in the two-room task. (B) Cell-type composition. Inactive: mean firing rate < 0.1 Hz; non-place: active without a field. Percentages denote category proportions. (C) Room-specific activity for both groups. (D) Between-room SCs in DG and across the CA3 transverse axis. Dots, individual neurons. (E) Correlation between CA3 recording position and mean between-room SC. Solid lines, significant; dashed, non-significant. (F) PVC heatmaps for DG (left) and CA3 (CON, top; LES, bottom). (G) Between-room PVCs. (H, I) Within-room SCs at cellular and tetrode levels, as in (D, E). (J) Within-room PVCs. (K) Stability loss (Δ within-room PVC) and discrimination loss (Δ between-room PVC) across CA3 bands. n.s., not significant; within-group: **, *P* < 0.01; ***, *P* ≤ 0.001; between-group: ^#^, *P* < 0.05; ^###^, *P* < 0.001.

By contrast, the lesion modestly compromised representational stability (20): within-room SCs were significantly reduced in lesioned animals (Fig. 2H), with clearly separated between-group SC at both the tetrode and animal levels (Fig. 2I; S3I). PVC analyses yielded similar results with graded loss of within-room stability (pCA3, 14%; mCA3, 8%, and dCA3, slightly negative; Fig. 2J). Decomposing population-level effects into stability loss (SL) and discrimination loss (DL) revealed that DL was negligible across the proximo-distal axis, whereas SL dominated the deficit in pCA3 (97%) and tapered progressively toward dCA3 (75% in mCA3, 40% in dCA3) (Fig. 2K). Notably, this effect was independent of CA1 lesion size (Table S3), and was not predicted by the intrinsic stability of individual DG neurons (Fig. 2H, J), suggesting that DG inputs maintain CA3 trial-over-trial stability through a circuit-level mechanism.

Alongside these stability deficits, theta power and spike-theta coupling increased in pCA3 following lesion, yielding a flattened proximo-distal gradient. The effect may reflect disrupted phase precession (21) (Fig. S4 A–E) driven by DG inputs (22). Measures of position selectivity were also reduced, with all deficits most pronounced in pCA3 (Fig. S4 F–M; group × band, all *P* < 0.049), consistent with earlier findings (17, 20, 23). These relatively mild impairments (< 29% in pCA3) suggest that graded mossy-fiber inputs (24, 25) contribute to CA3 transverse gradients in temporal and spatial coding, likely through coordinated excitatory drive and recruitment of inhibitory networks (26), supported by decreased interneuron firing rates after lesion (Fig. S4N).

### DG input gates rate-remapping along the transverse axis

We next assessed CA3 coding of non-spatial contextual change using a color-reversal paradigm, in which wall color was reversed while spatial geometry and other non-spatial cues remained constant. In controls, this manipulation elicited modest but significant rate remapping (Fig. 3A), defined as systematic firing-rate changes without shifts in place-field location (15), across all rats (Table S4). Along the CA3 proximo-distal axis, the proportion of place cells increased (Fig. 3B), whereas the magnitude of rate modulation progressively declined with minimal place-field displacement, at both cellular and tetrode levels (Fig. 3C–E; S5A–C). This graded rate remapping reflected small, regionally varying fractions of rate-remapping neurons (pCA3, 17.73%; mCA3, 4.76%; dCA3, 4.36%; Fig. 3F). Similar observations were made in both GCs and MCs in the DG, classified by a random-forest classifier (5) (Fig. S6), with rate coding comparable to that in pCA3 (Fig. 3C, D).

**Figure 3.**
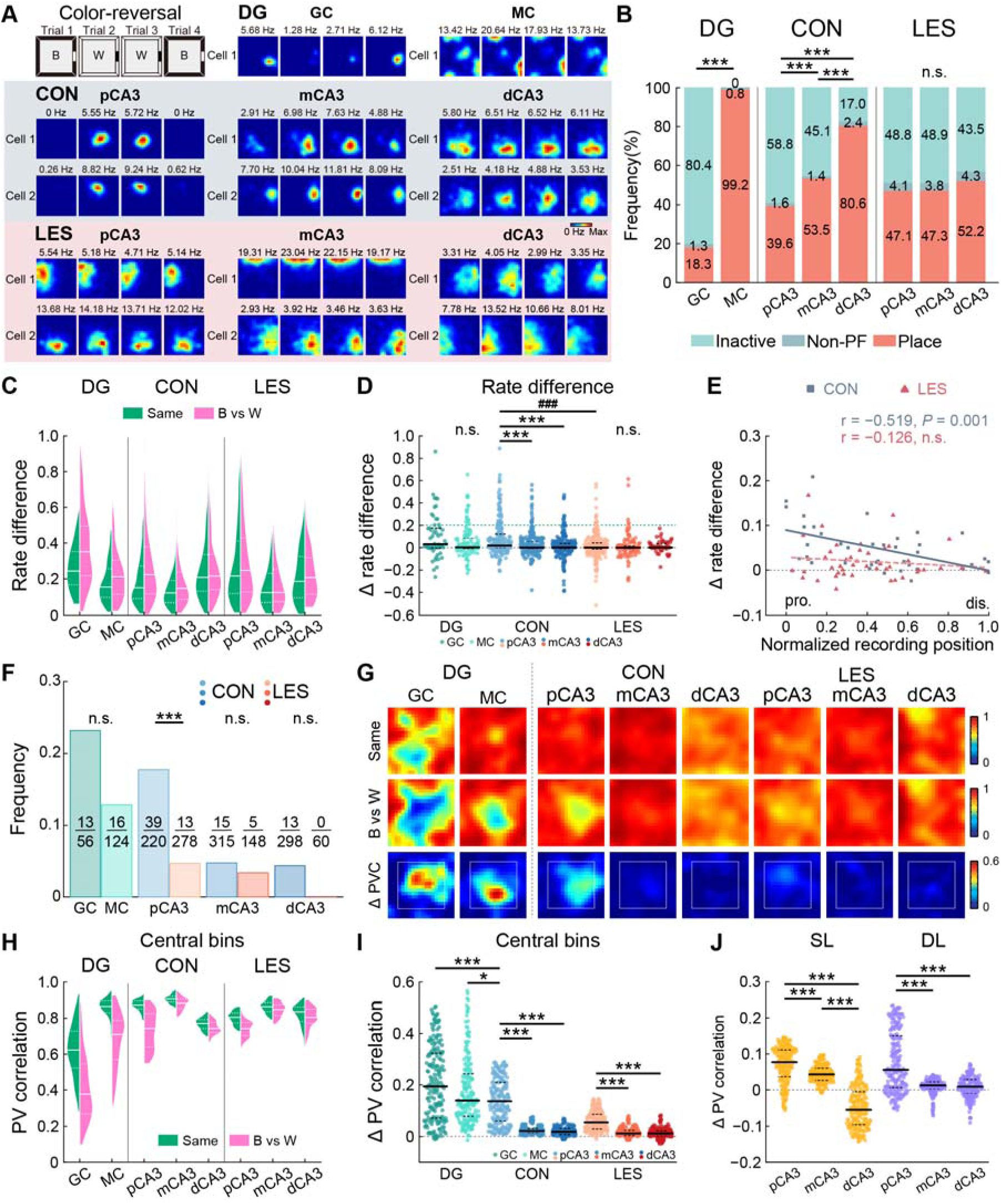
DG input gates rate-remapping along the transverse axis. (A) Representative rate maps in CON (top) and LES (bottom) rats in the color-reversal task. (B) Cell-type composition. (C) RD distributions across DG and CA3 subregions. (D) Color-reversal ΔRD (reversed-color RD − same-color RD). Green dashed line, threshold (0.2) for rate-modulated cells. (E) Correlation between CA3 position and mean ΔRD. (F) Fraction of rate-remapped cells (ΔRD > 0.2) by group. (G) PVC heatmaps for DG (left) and CA3 (CON, middle; LES, right). (H) PVC distributions for central bins across CA3 subregions. (I) Color-reversal ΔPVC for central bins. (J) Stability loss (SL, Δ same-color PVC) and discrimination loss (DL, Δ color-reversal PVC + SL) for central bins across CA3 bands. DL exceeded zero in all subregions. n.s., not significant; within-group: *, *P* < 0.05; ***, *P* < 0.001; between-group: ^###^, *P* < 0.001.

PVCs between color conditions remained high across regions (> 0.567; Fig. 3G), yet GCs exhibited the strongest color discrimination, quantified as ΔPVC (reversed vs. identical color decorrelation), decreasing progressively toward dCA3 (Fig. S5D–F), consistent with previous findings (11). Notably, rate modulation was expressed predominantly in the central arena (Fig. 3H, I; S5G, H), similar to a prior report (18). Furthermore, GCs in the suprapyramidal blade exhibited slightly stronger population-level responses than their infrapyramidal counterparts (Fig. S5I, J), consistent with preferential lateral entorhinal cortex (LEC) innervation of this blade (27), suggesting that LEC input density may modestly influence the magnitude of GC rate remapping (28, 29).

DG lesions modestly but significantly attenuated rate coding (Fig. 3A; Table S4) without altering navigational behavior (all *P* > 0.058; Table S1). The graded proportions of active place cells flattened after lesion (Fig. 3B) and the pCA3-selective population of rate-coding neurons was reduced by 73% (*P* < 0.001; Fig. 3F), with no significant changes in mCA3 or dCA3 (both *P* > 0.13). Accordingly, the pCA3-to-dCA3 gradient of changes in RD was abolished (group × band, *P* = 0.001; Fig. 3C, D), with convergence at the tetrode level (decrease, *P* = 0.002; correlation loss, *P* = 0.049; Fig. 3E) and animal level (Fig. S5K). The overall discrimination deficit was most pronounced in pCA3 (Fig. 3G; S5D–F). In the central arena, relative DL was highest in pCA3 (59.4%) and declined distally (mCA3, 45.5%; dCA3, 36.8%; Fig. 3J), whereas both color discrimination and DL following lesion were negligible for peripheral bins (Fig. S5G, H, L). These deficits were independent of spatial coding metrics (Table S5), and CA1 lesion size (Table S3). Given the modest baseline of color-induced rate remapping, we validated the lesion effect by downsampling to exclude sample-size artifacts (Fig. S7), and by identifying four animals with strong room-switch-induced rate remapping that was nearly abolished after lesion (Fig. S8). These convergent findings indicate that DG input contributes to CA3 rate remapping and preserves the proximo-distal gradient of cue discrimination.

### DG lesion abolishes gradient CA3 rate coding to object placement

Local objects provide event markers fundamental to episodic memory (30). We examined CA3 coding of object presence by introducing two identical objects into a fixed arena. In controls, object placement altered firing rates near object locations across a substantial fraction of neurons (Fig. 4A), consistent with earlier observations (19, 31). This object-induced rate modulation parallels classical rate remapping in that it alters firing rates without global ensemble reorganization (15), yet it differs in being spatially localized to object sites; we therefore refer to it as rate-based object coding (Fig. 4B). GCs generated the largest shift in population activity, exceeding that of MCs, with a clear pCA3-to-dCA3 gradient evident both globally and within a 20-cm object-centered region (Fig. 4C). Notably, elevated GC sensitivity in the suprapyramidal blade (Fig. S9A, B) likely reflects stronger LEC innervation (27), suggesting the LEC contribution to DG-mediated CA3 rate coding.

**Figure 4.**
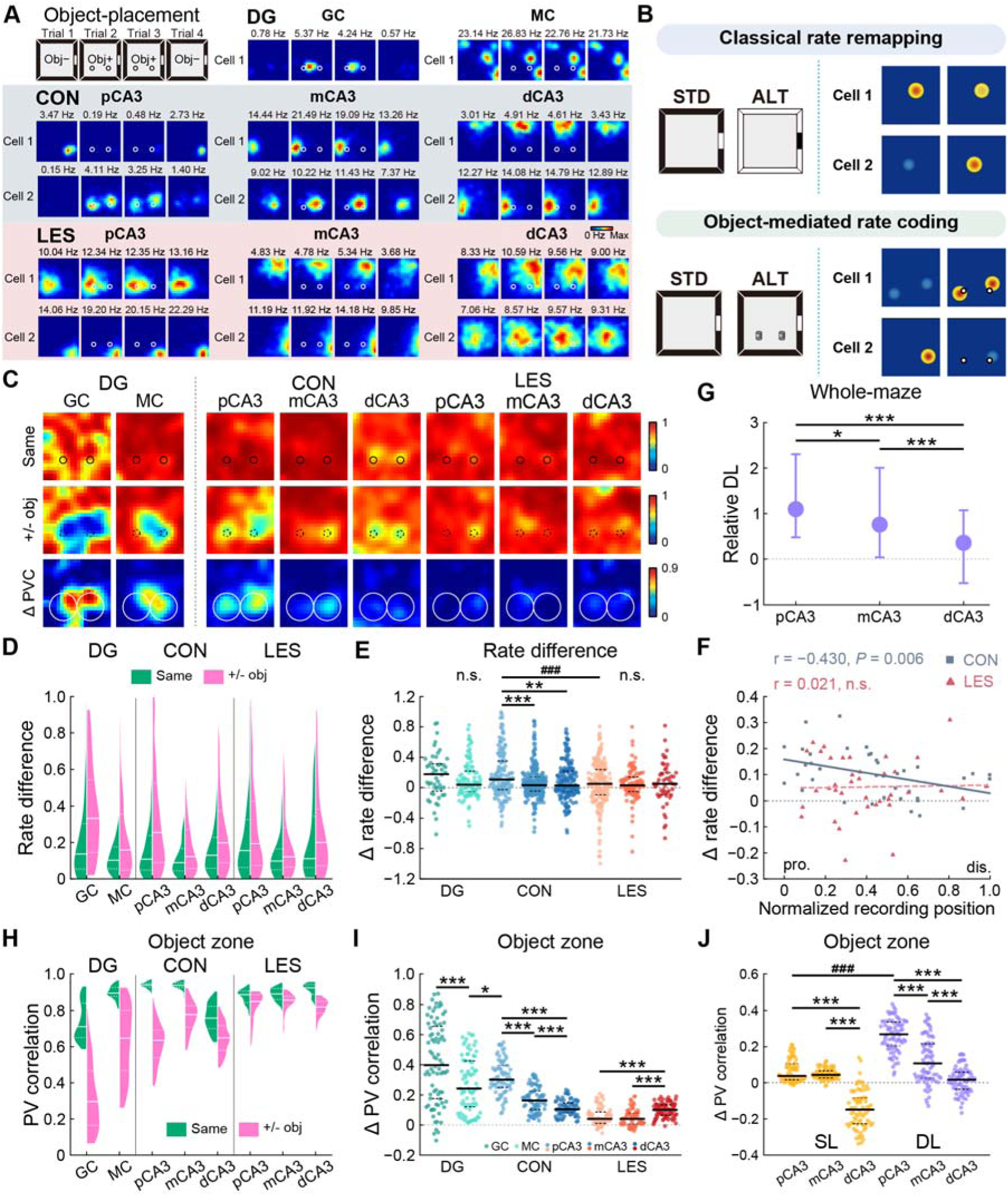
GC-driven rate coding of object placement is abolished by DG lesion. (A) Representative rate maps in CON (top) and LES (bottom) rats during object introduction. (B) Schematic: object-triggered rate modulation parallels classical rate remapping but is restricted to object-proximal regions. ALT, altered environment; STD, standard environment. (C) PVC heatmaps for DG (left) and CA3 (CON, middle; LES, right). (D) RD distributions across DG and CA3 subregions. (E) Object-induced ΔRD (object-induced RD – repeated-trial RD). (F) Correlation between CA3 position and mean ΔRD. (G) Relative DL (absolute DL / control discrimination). (H) PVC distributions in the object zone across CA3 subregions. (I) Object-induced ΔPVC in the object zone. (J) SL and DL in the object zone across CA3 bands. n.s., not significant; within-group: *, *P* < 0.05; **, *P* < 0.01; ***, *P* < 0.001; between-group: ###, *P* < 0.001.

DG lesions abolished this response without altering exploratory behavior (Fig. S9C). Whole-maze analyses revealed rate modulation deficits at the cellular (group, *P* = 0.025; group × band, *P* = 0.031; Fig. 4D, E; Table S1) and tetrode levels (ΔRD, *P* = 0.047; slope difference, *P* = 0.044; Fig. 4F). These deficits were replicated at the individual-animal level (Fig. S9D) and were partially present in SC (Fig. S9E–G), yet absent in other spatial measures (Table S5). PVC maps revealed focal responses at object sites in controls that were near-absent after lesion (Fig. S9H–J). The typical pCA3 > mCA3/dCA3 gradient in object-related discrimination reversed following DG loss, with nearly complete loss in pCA3 (∼100% relative DL) and progressively smaller deficits in mCA3 (76%) and dCA3 (36%) (Fig. 4G), parallel to mossy-fiber inputs (24, 25). This graded reduction was even more pronounced within object-centered regions (Fig. 4H–J). Thus, dentate input is essential for CA3 representation of local objects and modulates the expression of graded sensitivity of this representation along the transverse axis.

### Object relocation dissociates GC-driven presence signals from MC-driven removal signals

To dissociate coding of object position from object identity, we relocated one object between quadrants while the other remained stationary. In controls, CA3 neurons with fields near the current (CL) or original (OL) object location exhibited pronounced rate changes, with no modulation around the stationary object (Fig. 5A, B). Notably, the two loci were signaled by different dentate subpopulations: GCs preferentially signaled object presence at CL, whereas MCs signaled object absence at OL, producing equivalent modulation in pCA3 (Fig. 5C, D). The GC bias at CL persisted across individual rats (Fig. S10A) and aligned with LEC-dependent influences (Fig. S10B, C), supporting the view that the DG updates local object position (12, 32). MCs at OL exhibited net firing increases (Fig. S10D, E), signifying retrieval of an object-associated memory trace rather than simple loss of sensory drive (33).

**Figure 5.**
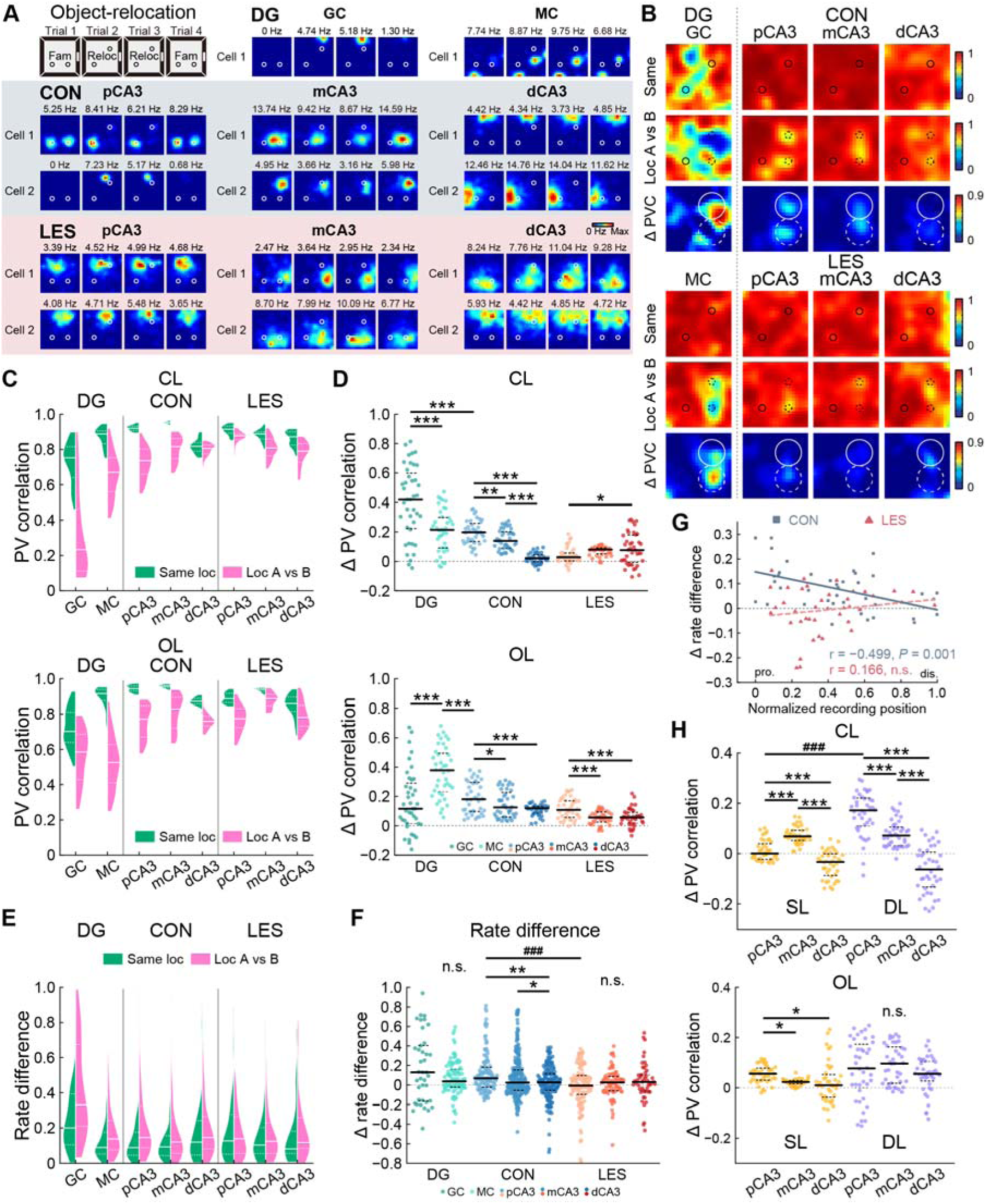
Object relocation dissociates GC-driven presence signals from MC-driven removal signals. (A) Representative rate maps in CON (top) and LES (bottom) rats during object relocation. (B) PVC heatmaps for DG (left) and CA3 (CON, middle; LES, right). (C) PVC distributions at current (CL, top) and original (OL, bottom) locations across CA3 subregions. (D) ΔPVC in CL (top) and OL (bottom) zones. (E) RD distributions across DG and CA3 subregions. (F) Relocation-induced ΔRD (relocation-induced RD – repeated-trial RD). (G) Correlation between CA3 position and mean ΔRD. (H) SL and DL in CL (top) and OL (bottom) zones across CA3 bands. n.s., not significant; within-group: *, *P* < 0.05; **, *P* < 0.01; ***, *P* ≤ 0.001; between-group: ^###^, *P* ≤ 0.001.

DG lesions eliminated CA3 rate responses to object relocation (Fig. 5A, B) without altering exploratory behavior (all *P* > 0.237; Table S1) or place field locations (Fig. S10F–H). Whole-maze rate modulation was reduced with a graded pattern across the transverse axis, with elevated ΔRD in pCA3 response abolished at both cellular (group × band, *P* = 0.003; Fig. 5E, F) and tetrode levels (ΔRD, *P* < 0.001; slope difference, *P* = 0.003; Fig. 5G). A similar trend was evident across animals (Fig. S10I). At the population level, focal responses at CL and OL were significantly diminished (Fig. 5B–D), paralleling whole-maze responses (Fig. S10J, K). These deficits were not detected by other spatial measures (Table S5) or correlated with CA1 lesion size (Table S3).

Decomposing by locus revealed the source of this loss: at CL, DG depletion reversed the typical proximodistal gradient in ΔPVC, with severe loss localized to pCA3 (Fig. 5H, top); at OL, responses preserved the normal gradient but showed uniform reductions in magnitude (Fig. 5H, bottom). These regionally distinct effects mirror the object-placement results (Fig. 4H, I) and align with complementary GC and MC coding (Fig. 5D). Together, these findings indicate that GCs and MCs signal local object information through parallel pathways. GCs drive CA3 coding of object presence via mossy fibers, whereas MCs sustain removal-related signals via associational connections. Both converge to shape CA3 representations of object position.

## Discussion

Hippocampal contextual discrimination operates through global remapping and rate remapping (15), yet their differential dependence on DG input has remained unclear. By combining focal DG lesions with spatially resolved CA3 recordings, we found that intact DG output enhances rate remapping and rate-based object coding, sharpens the proximo-distal gradient in temporal, spatial, and discriminative coding, while exerting limited influence on global remapping. The resulting deficits were sharpest in pCA3 and attenuated distally, paralleling mossy-fiber topography, indicating that DG circuit integrity amplifies the proximo-distal gradient of rate-based discrimination without being required for categorical global remapping. Furthermore, GCs and MCs complemented one another by signaling object presence and removal, respectively. These findings establish that the DG network preferentially amplifies fine-grained contextual discrimination through rate modulation, consistent with a major contribution from topographically organized GC-to-CA3 projections.

### Robust GC response to contextual manipulations

Prior studies have reported inconsistent remapping strength in GCs (2, 3, 5, 6), likely reflecting sampling bias: reports of weak remapping have relied on the minority of GCs active across contexts, overlooking the selectively engaged majority (8, 34). In the present data, room discrimination reached its maximum when the entire GC population was analyzed, yielding constant decorrelation across GC, MC and pCA3 pyramidal neurons, highlighting how sampling bias can distort estimates of remapping magnitude. During rate-based manipulations, GCs exhibited stronger rate modulation than MCs and surpassed pCA3 response amplitude under most conditions, indicating that the DG primarily amplifies downstream discrimination through enhanced rate coding.

### DG influences CA3 computation via direct and indirect pathways

Anatomical and physiological evidence indicates that the DG projects predominantly to pCA3 via mossy fibers (24), with excitatory postsynaptic currents tapering distally (25). The observed gradient in rate remapping loss (> 59% in pCA3) is consistent with a major contribution from the topographically organized GC-to-CA3 pathway. Meanwhile, the mild reduction in theta precession and spatial selectivity (< 29% in pCA3) suggests an indirect pathway involving GC-driven interneuron networks that regulate theta oscillations and place field formation (35–37). Furthermore, object-removal signals at the original location were sustained by MC associational input, consistent with retrieval of an object-associated memory trace (16, 33, 38) rather than simple loss of sensory drive, whereas graded responses at the current location align with GC-driven mossy-fiber input. These findings suggest parallel mechanisms through which GCs and MCs coordinate CA3 coding of local object position, although their precise synaptic architecture and cellular effectors remain to be determined.

### Entorhinal inputs support dual-mode contextual discrimination in CA3

Global remapping is elicited by substantial afferent divergence (15) and is often accompanied by MEC realignments (39, 40) that engage new hippocampal ensembles (41) (Fig. 6). Although MEC realignment may not be strictly necessary for hippocampal global remapping (42, 43), shifted MEC input can nevertheless contribute to hippocampal discrimination via heavy perforant projections, independently of DG input. Consistent with this, lesions left between-room discrimination largely intact while attenuating the proximo-distal gradient in active neurons. In contrast, rate-based contextual coding is evoked by finer sensory changes without shifts in neuron identity (15, 31). Our finding of enhanced rate coding in the dentate blade receiving stronger LEC innervations is consistent with the view that LEC provides sensory input for rate remapping (28, 44). Because LEC neurons themselves exhibit minimal rate modulation, we speculate that sparse GC activation transforms sensory divergence into a hippocampal rate code, selectively enhancing the resolution of ambiguous inputs before their relay to CA3. Therefore, by integrating our causal findings in the DG-CA3 network with established models of entorhinal function, we propose a working model in which entorhinal inputs upstream can support global remapping through multiple pathways, whereas DG output downstream amplifies the full expression of rate-based discrimination (Fig. 6).

**Figure 6.**
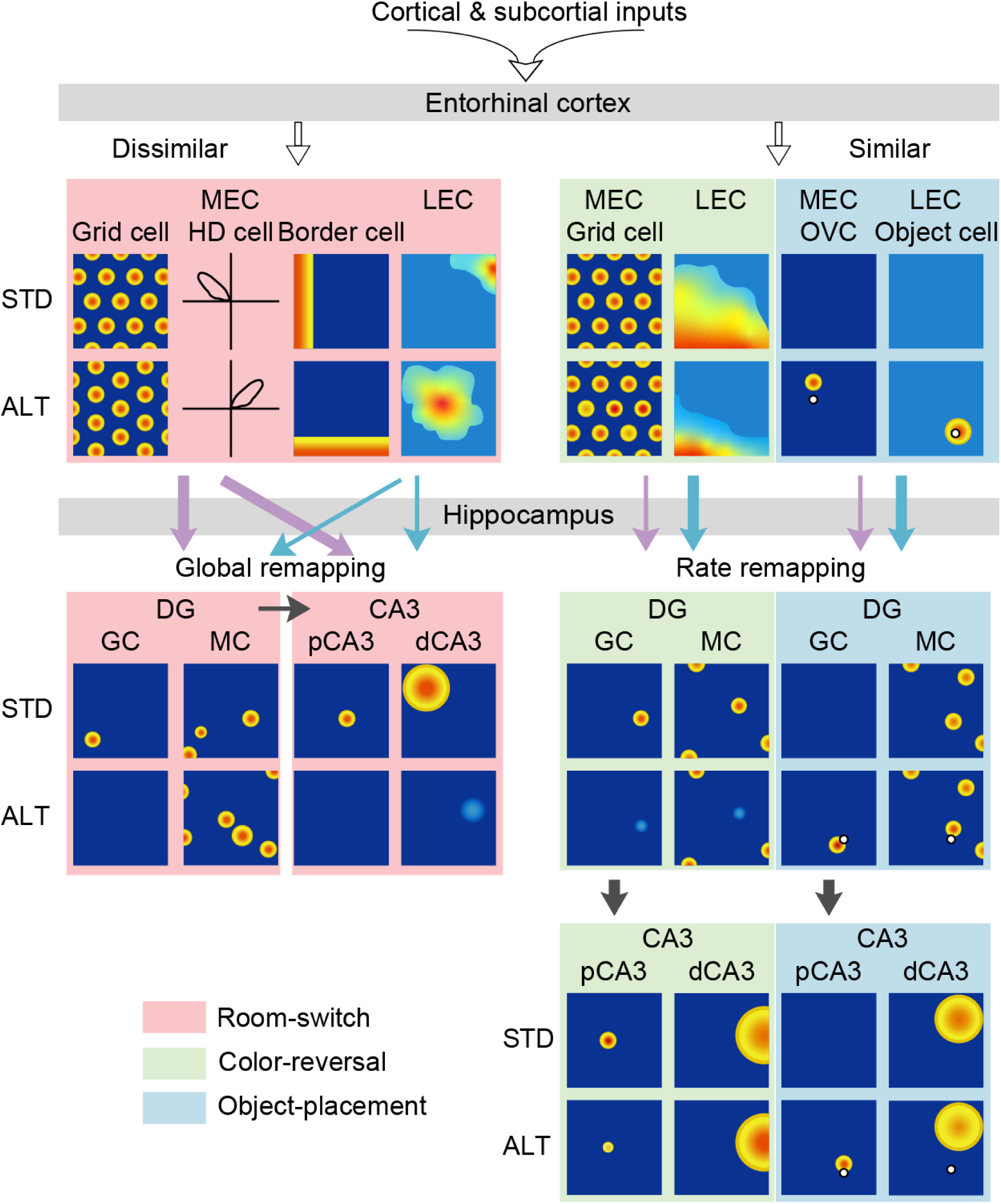
Working model of the entorhinal-hippocampal contextual discrimination. Top: Cortical and subcortical inputs converge on the entorhinal cortex. Left: Categorical changes can engage MEC realignment (prior work) to support CA3 global remapping independently of DG input (present findings); DG may nonetheless modulate the proximo-distal gradient in active neurons. Right: During subtle changes, stable MEC grids and LEC sensory input (prior work) reach the DG. We propose that sparse GC firing transforms these inputs into rate-modulated outputs that amplify rate-based discrimination in pCA3 (present findings). Abbreviations: ALT, altered environment; dCA3, distal CA3; DG, dentate gyrus; GC, granule cell; HD, head direction; MC, mossy cell; MEC, medial entorhinal cortex; LEC, lateral entorhinal cortex; OVC, object-vector cell; STD, standard environment. Activity patterns adapted from (39, 40, 44).

### Limitations of the study

First, the absence of parametrically varied sensory input precludes direct conclusions regarding pattern separation, restricting our findings to remapping phenomena. Second, the possible influence of partial CA1 damage through the CA1→EC→DG feedback loop (45) was not directly examined, yet the lack of correlation between CA1 lesion size and CA3 discrimination loss argues against this confound. Third, colchicine lesions affect GCs, MCs, and associated interneurons indiscriminately, precluding attribution to individual cell types and raising the possibility of compensatory plasticity. Finally, our classifier has not been validated for adult-born GCs (3, 5), which are important for pattern separation (46). Future studies combining pathway-specific manipulations such as optogenetic or chemogenetic silencing (47, 48) with transgenic labeling of DG subpopulations will be needed to clarify how distinct cell types and circuits shape CA3 rate coding across diverse contextual demands.

## Materials and Methods

### Subjects and surgery

Sixteen male Long-Evans rats (3–4 months) were used. Seven received bilateral DG lesions via colchicine micro-infusion and tetrode implantation targeting the right CA3; six sham-operated controls received PBS infusion and identical implants. Anesthesia was maintained with isoflurane and post-operative analgesia was administered for 5–7 days (see Supplementary Methods for details). All procedures were approved by the Institutional Animal Care and Use Committee of the Kunming Institute of Zoology (KIZ-IACUC-RE-2021-06-019).

### Electrophysiology

An 18-tetrode hyperdrive (49) was implanted for each animal. Tetrodes were advanced into either the DG cell layer or the CA3 pyramidal layer (44). Electrophysiological recording procedures followed previously described methods (18). Neural signals were amplified, band-pass filtered (300 Hz–7.5 kHz), and digitized at 30 kHz. Spike waveforms and local field potentials (0.3–300 Hz) were recorded simultaneously. Position was tracked at 50 Hz via head-mounted LEDs.

### Behavioral paradigms

Rats foraged for scattered food in a 100 × 100 cm square arena surrounded by curtains. All tasks followed an A-B-B-A design (10 min per trial, 5-min inter-trial rests). The four tasks were: (1) color-reversal (black vs. white walls); (2) two-room (identical arenas in distinct rooms); (3) object-placement (two identical objects introduced); and (4) object-relocation (one object moved between quadrants). Rats were pre-trained on the color-reversal task until consistent exploration was achieved.

### Histology and lesion quantification

After recording, rats were perfused and brains were sectioned coronally at 40 µm. Nissl staining and immunofluorescence (calbindin, PCP4, parvalbumin) were used to verify recording sites, quantify spared tissue, and assess MC and interneuron counts. Tetrode positions were normalized along the CA3 transverse axis and classified as proximal (pCA3), middle (mCA3), or distal (dCA3).

### Behavioral metrics

Navigational behavior was quantified by speed, tortuosity, arena coverage, and spatial evenness (50). Object/wall exploration was measured by occupation time within 20 cm of objects/along the walls and discrimination index.

### Spike sorting and cell classification

Single units were isolated offline with MClust. Putative excitatory neurons were identified by spike width (> 200 µs), mean firing rate (< 5 Hz), and burst discharge patterns (11). An additional filter of spatial information content > 0.5 bits/spike was applied to exclude putative hilar interneurons (5). Putative CA3 interneurons were classified based on their spike width, firing rate and waveform. Units failing criteria or recorded repeatedly in the same task were excluded (18). DG principal neurons were classified into granule cells (GCs) and mossy cells (MCs) using a random-forest classifier (5) based on firing rate, burstiness, rest-to-navigation ratio, and peak in-box rate. Classification confidence was quantified by vote proportion across 300 decision trees.

### Spatial and rate analyses

Rate maps were constructed in 5 × 5 cm bins, smoothed with a Gaussian kernel, and place fields were defined by contiguous regions exceeding 20% of the cell’s peak rate with a minimum peak of 1 Hz (18). Spatial modulation was quantified by spatial information rate and content (51), spatial coherence (first-order autocorrelation between adjacent bins), and spatial stability (correlation between first- and second-half rate maps). Spatial correlation (SC) was computed as the Pearson correlation between rate maps from different trials. Animals with mean SC values across all active CA3 pyramidal neurons higher than 0.5 were classified as exhibiting rate remapping (18). Population vector correlation (PVC) was computed from composite population firing arrays across spatial bins (15). Rate difference (RD) and rate index (RI) quantified normalized firing changes between conditions at the whole-arena and object-zone levels, respectively.

### Stability and discrimination loss

Population-level effects were decomposed into stability loss (SL; reduced within-condition PVC after lesion) and discrimination loss (DL; reduced between-condition PVC after lesion). Relative contributions were expressed as proportions of the total deficit.

### LFP and temporal analyses

Time-frequency decomposition used continuous wavelet transforms. Theta power and spike-theta phase coupling were quantified after removing aperiodic components including dentate spikes and other non-oscillatory DG activity (52, 53). Phase precession was estimated from the difference between intrinsic theta frequency (spike train autocorrelogram) and local theta frequency (LFP power spectrum) (54).

### Statistics

Linear mixed-effects models examined fixed effects of group, CA3 band, and their interaction, with random intercepts for neurons nested within rats. Models with random slopes were evaluated but not used (Table S6), due to unbalanced cell sampling across animals and CA3 bands (55). PVCs were analyzed with repeated-measures two-way ANOVA. Significant interactions were followed by Kruskal–Wallis or Friedman tests. Pairwise comparisons used paired or unpaired *t*-tests or Wilcoxon rank-sum/signed-rank tests with Holm–Bonferroni correction. Permutation tests (10,000 iterations) assessed differences in regression slopes. Frequency data were compared using Pearson’s chi-square or Fisher’s exact tests. Linear relationships were assessed with Pearson correlation coefficients. Multiple comparisons between frequency bands were corrected using false discovery rate (FDR). The cell (or spatial bin for PVC, tetrode for LFP) was the unit of analysis. All tests were two-tailed; significance was set at *P* < 0.05.

## Data availability

Data and code are available from the Lead Contact upon request. A preprint of this work is available on bioRxiv (doi: 10.64898/2025.12.04.692471).

## Acknowledgments

This research was supported by grants from the Science and Technology Innovation (STI) 2030-Major Projects (2022ZD0205000 to L.L.), National Natural Science Foundation of China (31970963 to L.L.), CAS “Light of West China” Program (xbzg-zdsys-202404 to L.L.), Yunnan Revitalization Talent Support Program (Yunling Scholar Project to L.L.), Yunnan Province (202305AH340006 to L.L.), and Yunnan Fundamental Research Projects (202401AT070206 to X.C.).

The authors express gratitude to Cheng-Ji Li, Rong Zhang, Xiao-Fan Ge, Chu Deng, Yi-Qing Hui, and Yi-Fan Ye, as well as the staff of the National Research Facility for Phenotypic & Genetic Analysis of Model Animals (Primate Facility) (https://cstr.cn/31137.02.NPRC), for providing technical support and assistance in data collection and analysis.

## Supplementary Material

Supplementary material is accessible in the online version of this paper.

## Notes

### Competing Interest Statement

The authors have declared no competing interest.

### Summary of Updates

Version 3 has been substantially revised to sharpen the central conclusion that DG input differentially contributes to distinct modes of CA3 contextual coding. The conceptual framework was refined to more clearly distinguish categorical environmental changes represented by global remapping from finer contextual differences represented through rate-based coding. The Results were reorganized to emphasize that DG lesion leaves between-room global remapping largely intact while reducing within-room representational stability, with the strongest effects in proximal CA3 and progressively weaker effects toward distal CA3. The color-reversal analyses were also revised to more clearly demonstrate the preferential attenuation of rate-based contextual coding in proximal CA3 following DG lesion and the disruption of the normal transverse gradient along the CA3 axis. The object-related results were reorganized to highlight the complementary contributions of dentate granule cells and mossy cells to local object information and the preferential dependence of proximal CA3 responses on DG input. The Abstract, Introduction, Discussion, figures, Methods, and Supporting Information were revised accordingly to improve consistency with this central framework and to present the findings in a clearer and more focused manner. Overall, these revisions strengthen the organization and interpretation of the manuscript and better align the conclusions with the experimental evidence.

## References

1. L. L. Colgin, E. I. Moser, M. B. Moser, Understanding memory through hippocampal remapping. Trends Neurosci 31, 469–477 (2008).

2. J. P. Neunuebel, J. J. Knierim, CA3 Retrieves Coherent Representations from Degraded Input: Direct Evidence for CA3 Pattern Completion and Dentate Gyrus Pattern Separation. Neuron 81, 416–427 (2014).

3. D. GoodSmith, H. Lee, J. P. Neunuebel, H. Song, J. J. Knierim, Dentate gyrus mossy cells share a role in pattern separation with dentate granule cells and proximal CA3 pyramidal cells. J Neurosci 39, 9570–9584 (2019).

4. N. B. Danielson et al., In vivo imaging of dentate gyrus mossy cells in behaving mice. Neuron 93, 552–559 e554 (2017).

5. D. GoodSmith et al., Spatial representations of granule cells and mossy cells of the dentate gyrus. Neuron 93, 677–690 e675 (2017).

6. Y. Senzai, G. Buzsaki, Physiological properties and behavioral correlates of hippocampal granule cells and mossy cells. Neuron 93, 691–704 e695 (2017).

7. J. P. Neunuebel, J. J. Knierim, Spatial firing correlates of physiologically distinct cell types of the rat dentate gyrus. J Neurosci 32, 3848–3858 (2012).

8. S. H. Kim et al., Global remapping in granule cells and mossy cells of the mouse dentate gyrus. Cell Rep 42, 112334 (2023).

9. L. W. Huang, F. Torelli, H. L. Chen, M. Bartos, Context and space coding in mossy cell population activity. Cell Rep 43, 114386 (2024).

10. H. Lee, C. Wang, S. S. Deshmukh, J. J. Knierim, Neural population evidence of functional heterogeneity along the CA3 transverse axis: pattern completion versus pattern separation. Neuron 87, 1093–1105 (2015).

11. L. Lu, K. M. Igarashi, M. P. Witter, E. I. Moser, M. B. Moser, Topography of place maps along the CA3-to-CA2 axis of the hippocampus. Neuron 87, 1078–1092 (2015).

12. I. Lee, F. Solivan, Dentate gyrus is necessary for disambiguating similar object-place representations. Learn Mem 17, 252–258 (2010).

13. A. M. Morris, C. S. Weeden, J. C. Churchwell, R. P. Kesner, The role of the dentate gyrus in the formation of contextual representations. Hippocampus 23, 162–168 (2013).

14. C. H. Lee, I. Lee, Impairment of pattern separation of ambiguous scenes by single units in the CA3 in the absence of the dentate gyrus. J Neurosci 40, 3576–3590 (2020).

15. S. Leutgeb et al., Independent codes for spatial and episodic memory in hippocampal neuronal ensembles. Science 309, 619–623 (2005).

16. D. G. Amaral, H. E. Scharfman, P. Lavenex, The dentate gyrus: fundamental neuroanatomical organization (dentate gyrus for dummies). Prog Brain Res 163, 3–22 (2007).

17. T. Sasaki et al., Dentate network activity is necessary for spatial working memory by supporting CA3 sharp-wave ripple generation and prospective firing of CA3 neurons. Nat Neurosci 21, 258–269 (2018).

18. Y. L. Duan et al., Object-translocation induces event coding in the rat hippocampus. Commun Biol 8, 797 (2025).

19. D. GoodSmith et al., Flexible encoding of objects and space in single cells of the dentate gyrus. Curr Biol 32, 1088–1101.e1085 (2022).

20. X. Chen, N. Cheng, C. Wang, J. J. Knierim, Impaired spatial coding of the hippocampus in a dentate gyrus hypoplasia mouse model. Proc Natl Acad Sci U S A 122, e2416214122 (2025).

21. M. W. Jones, M. A. Wilson, Phase precession of medial prefrontal cortical activity relative to the hippocampal theta rhythm. Hippocampus 15, 867–873 (2005).

22. S. Ahmadi et al., Distinct roles of dentate gyrus and medial entorhinal cortex inputs for phase precession and temporal correlations in the hippocampal CA3 area. Nat Commun 16, 13 (2025).

23. B. L. McNaughton, C. A. Barnes, J. Meltzer, R. J. Sutherland, Hippocampal granule cells are necessary for normal spatial learning but not for spatially-selective pyramidal cell discharge. Exp Brain Res 76, 485–496 (1989).

24. M. P. Witter, Intrinsic and extrinsic wiring of CA3: indications for connectional heterogeneity. Learn Mem 14, 705–713 (2007).

25. Q. Sun et al., Proximodistal heterogeneity of hippocampal CA3 pyramidal neuron intrinsic properties, connectivity, and reactivation during memory recall. Neuron 95, 656–672 e653 (2017).

26. L. Acsády, A. Kamondi, A. Sík, T. Freund, G. Buzsáki, GABAergic cells are the major postsynaptic targets of mossy fibers in the rat hippocampus. J Neurosci 18, 3386–3403 (1998).

27. N. Tamamaki, Organization of the entorhinal projection to the rat dentate gyrus revealed by Dil anterograde labeling. Exp Brain Res 116, 250–258 (1997).

28. C. Rennó-Costa, J. E. Lisman, P. F. Verschure, The mechanism of rate remapping in the dentate gyrus. Neuron 68, 1051–1058 (2010).

29. T. Cholvin, M. Bartos, The dentate gyrus efficiently converges LEC and MEC inputs into multimodal, highly specific and reliable environmental representations. Nat Neurosci 10.1038/s41593-026-02240-0 (2026).

30. S. S. Deshmukh, J. J. Knierim, Influence of local objects on hippocampal representations: Landmark vectors and memory. Hippocampus 23, 253–267 (2013).

31. A. Nagelhus, S. O. Andersson, S. G. Cogno, E. I. Moser, M. B. Moser, Object-centered population coding in CA1 of the hippocampus. Neuron 111, 2091–2104 e2014 (2023).

32. M. R. Hunsaker, J. S. Rosenberg, R. P. Kesner, The role of the dentate gyrus, CA3a,b, and CA3c for detecting spatial and environmental novelty. Hippocampus 18, 1064–1073 (2008).

33. X. Li et al., A circuit of mossy cells controls the efficacy of memory retrieval by Gria2I inhibition of Gria2. Cell Rep 34, 108741 (2021).

34. T. Hainmueller, M. Bartos, Parallel emergence of stable and dynamic memory engrams in the hippocampus. Nature 558, 292–296 (2018).

35. A. Chadwick, M. C. van Rossum, M. F. Nolan, Flexible theta sequence compression mediated via phase precessing interneurons. Elife 5 (2016).

36. C. Y. Huh et al., Excitatory Inputs Determine Phase-Locking Strength and Spike-Timing of CA1 Stratum Oriens/Alveus Parvalbumin and Somatostatin Interneurons during Intrinsically Generated Hippocampal Theta Rhythm. J Neurosci 36, 6605–6622 (2016).

37. M. Valero, A. Navas-Olive, L. M. de la Prida, G. Buzsáki, Inhibitory conductance controls place field dynamics in the hippocampus. Cell Rep 40, 111232 (2022).

38. S. Jinde et al., Hilar mossy cell degeneration causes transient dentate granule cell hyperexcitability and impaired pattern separation. Neuron 76, 1189–1200 (2012).

39. T. Solstad, C. N. Boccara, E. Kropff, M. B. Moser, E. I. Moser, Representation of geometric borders in the entorhinal cortex. Science 322, 1865–1868 (2008).

40. M. Fyhn, T. Hafting, A. Treves, M. B. Moser, E. I. Moser, Hippocampal remapping and grid realignment in entorhinal cortex. Nature 446, 190–194 (2007).

41. T. Solstad, H. N. Yousif, T. J. Sejnowski, Place cell rate remapping by CA3 recurrent collaterals. PLoS Comput Biol 10, e1003648 (2014).

42. M. P. Brandon, J. Koenig, J. K. Leutgeb, S. Leutgeb, New and distinct hippocampal place codes are generated in a new environment during septal inactivation. Neuron 82, 789–796 (2014).

43. M. I. Schlesiger, B. L. Boublil, J. B. Hales, J. K. Leutgeb, S. Leutgeb, Hippocampal Global Remapping Can Occur without Input from the Medial Entorhinal Cortex. Cell Rep 22, 3152–3159 (2018).

44. L. Lu et al., Impaired hippocampal rate coding after lesions of the lateral entorhinal cortex. Nat Neurosci 16, 1085–1093 (2013).

45. 45. N. L. Cappaert, N. M. van Strien, M. P. Witter, “Hippocampal Formation” in The Rat Nervous System, G. Paxinos, Ed. (Academic Press, 2015), 10.1016/B978-0-12-374245-2.00020-6 chap. 20, pp. 511–573.

46. C. D. Clelland et al., A functional role for adult hippocampal neurogenesis in spatial pattern separation. Science 325, 210–213 (2009).

47. L. Madisen et al., A toolbox of Cre-dependent optogenetic transgenic mice for light-induced activation and silencing. Nat Neurosci 15, 793–802 (2012).

48. B. L. Roth, DREADDs for Neuroscientists. Neuron 89, 683–694 (2016).

49. L. Lu, B. Popeney, J. D. Dickman, D. E. Angelaki, Construction of an Improved Multi-Tetrode Hyperdrive for Large-Scale Neural Recording in Behaving Rats. J Vis Exp 10.3791/57388 (2018).

50. P. A. P. Moran, The Interpretation of Statistical Maps. Journal of the Royal Statistical Society: Series B (Methodological) 10, 243–251 (1948).

51. W. E. Skaggs, B. L. McNaughton, M. A. Wilson, C. A. Barnes, Theta phase precession in hippocampal neuronal populations and the compression of temporal sequences. Hippocampus 6, 149–172 (1996).

52. S. B. McHugh et al., Offline hippocampal reactivation during dentate spikes supports flexible memory. Neuron 112, 3768–3781.e3768 (2024).

53. T. Donoghue et al., Parameterizing neural power spectra into periodic and aperiodic components. Nat Neurosci 23, 1655–1665 (2020).

54. T. Eliav et al., Nonoscillatory Phase Coding and Synchronization in the Bat Hippocampal Formation. Cell 175, 1119–1130 e1115 (2018).

55. X. A. Harrison et al., A brief introduction to mixed effects modelling and multi-model inference in ecology. PeerJ 6, e4794 (2018).

