## Supplementary materials for "Dentate gyrus amplifies rate-based coding while sparing global remapping across the CA3 transverse axis"

Li Lu

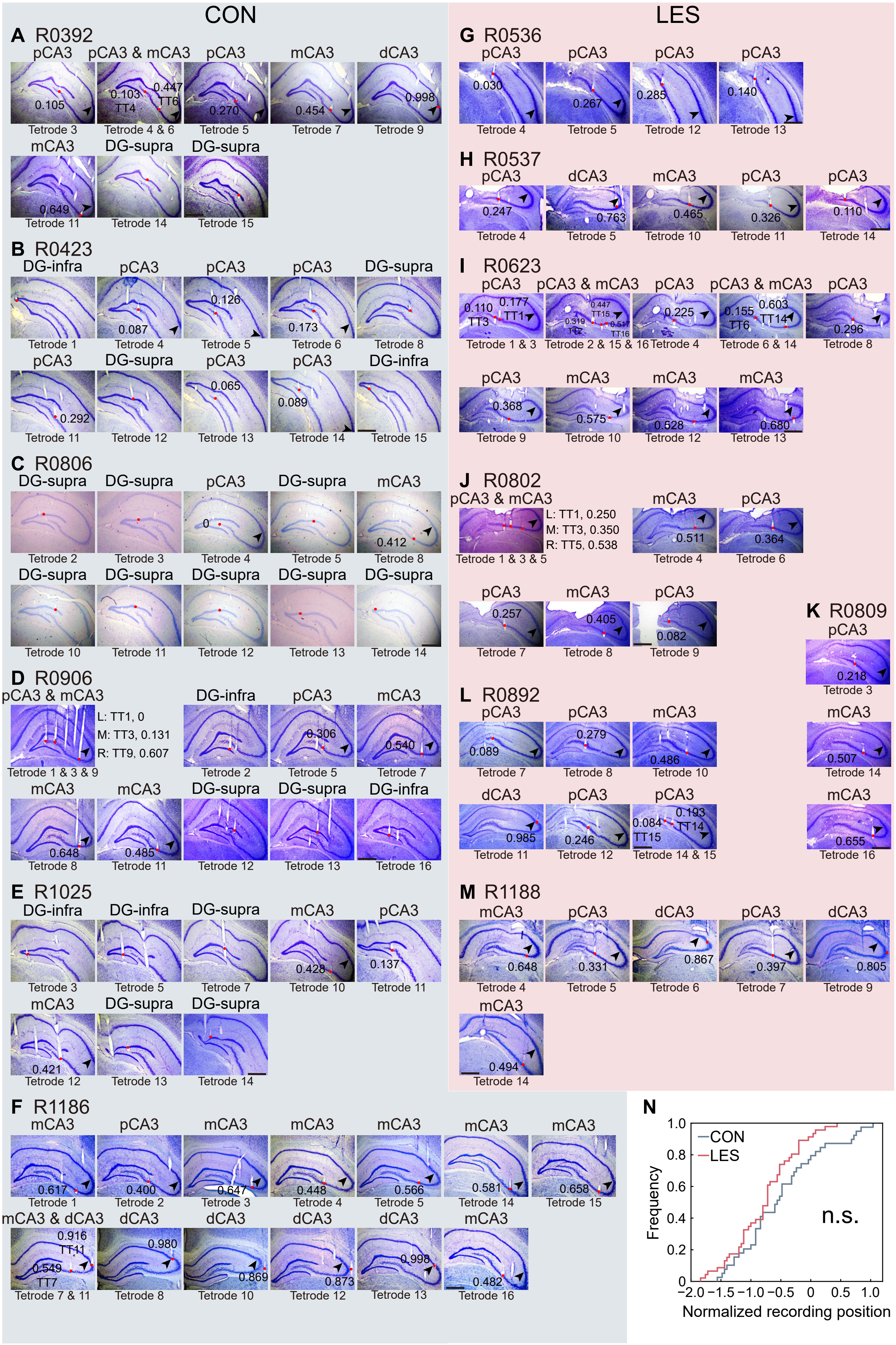
CON
LES
A R0392
pCA3 pCA3 & mCA3 pCA3 mCA3 dCA3
0.105 0.103 TT4 0.447 TT6 0.270 0.454 0.998
Tetrode 3 Tetrode 4 & 6 Tetrode 5 Tetrode 7 Tetrode 9
mCA3 DG-supra DG-supra
0.649
Tetrode 11 Tetrode 14 Tetrode 15
B R0423
DG-infra pCA3 pCA3 pCA3 DG-supra
0.087 0.126 0.173
Tetrode 1 Tetrode 4 Tetrode 5 Tetrode 6 Tetrode 8
pCA3 DG-supra pCA3 pCA3 DG-infra
0.292 0.065 0.089
Tetrode 11 Tetrode 12 Tetrode 13 Tetrode 14 Tetrode 15
C R0806
DG-supra DG-supra pCA3 DG-supra mCA3
0 0.412
Tetrode 2 Tetrode 3 Tetrode 4 Tetrode 5 Tetrode 8
DG-supra DG-supra DG-supra DG-supra DG-supra
Tetrode 10 Tetrode 11 Tetrode 12 Tetrode 13 Tetrode 14
D R0906
pCA3 & mCA3 DG-infra pCA3 mCA3
L: TT1, 0
M: TT3, 0.131
R: TT9, 0.607
0.306 0.540
Tetrode 1 & 3 & 9 Tetrode 2 Tetrode 5 Tetrode 7
mCA3 mCA3 DG-supra DG-supra DG-infra
0.648 0.485
Tetrode 8 Tetrode 11 Tetrode 12 Tetrode 13 Tetrode 16
E R1025
DG-infra DG-infra DG-supra mCA3 pCA3
0.428 0.137
Tetrode 3 Tetrode 5 Tetrode 7 Tetrode 10 Tetrode 11
mCA3 DG-supra DG-supra
0.421
Tetrode 12 Tetrode 13 Tetrode 14
F R1186
mCA3 pCA3 mCA3 mCA3 mCA3 mCA3 mCA3
0.617 0.400 0.647 0.448 0.566 0.581 0.658
Tetrode 1 Tetrode 2 Tetrode 3 Tetrode 4 Tetrode 5 Tetrode 14 Tetrode 15
mCA3 & dCA3 dCA3 dCA3 dCA3 dCA3 mCA3
0.916 TT11 0.549 TT7 0.980 0.869 0.873 0.998 0.482
Tetrode 7 & 11 Tetrode 8 Tetrode 10 Tetrode 12 Tetrode 13 Tetrode 16
G R0536
pCA3 pCA3 pCA3 pCA3
0.030 0.267 0.285 0.140
Tetrode 4 Tetrode 5 Tetrode 12 Tetrode 13
H R0537
pCA3 dCA3 mCA3 pCA3 pCA3
0.247 0.763 0.465 0.326 0.110
Tetrode 4 Tetrode 5 Tetrode 10 Tetrode 11 Tetrode 14
I R0623
pCA3 pCA3 & mCA3 pCA3 pCA3 & mCA3 pCA3
0.110 TT3 0.177 TT1 0.447 TT15 0.319 TT2 0.517 TT16 0.225 0.155 TT6 0.603 TT14 0.296
Tetrode 1 & 3 Tetrode 2 & 15 & 16 Tetrode 4 Tetrode 6 & 14 Tetrode 8
pCA3 mCA3 mCA3 mCA3
0.368 0.575 0.528 0.680
Tetrode 9 Tetrode 10 Tetrode 12 Tetrode 13
J R0802
pCA3 & mCA3 mCA3 pCA3
L: TT1, 0.250
M: TT3, 0.350
R: TT5, 0.538
0.511 0.364
Tetrode 1 & 3 & 5 Tetrode 4 Tetrode 6
pCA3 mCA3 pCA3
0.257 0.405 0.082
Tetrode 7 Tetrode 8 Tetrode 9
K R0809
pCA3
0.218
Tetrode 3
L R0892
pCA3 pCA3 mCA3 mCA3
0.089 0.279 0.486 0.507
Tetrode 7 Tetrode 8 Tetrode 10 Tetrode 14
dCA3 pCA3 pCA3 mCA3
0.985 0.246 0.084 TT15 0.193 TT14 0.655
Tetrode 11 Tetrode 12 Tetrode 14 & 15 Tetrode 16
M R1188
mCA3 pCA3 dCA3 pCA3 dCA3
0.648 0.331 0.867 0.397 0.805
Tetrode 4 Tetrode 5 Tetrode 6 Tetrode 7 Tetrode 9
mCA3
0.494
Tetrode 14
N
CON
LES
n.s.
Frequency
0 0.2 0.4 0.6 0.8 1.0
−2.0 −1.5 −1.0 −0.5 0 0.5 1.0
Normalized recording position

**Figure S1. Recording-site anatomy in control and DG-lesioned rats.**

(A–M) Nissl-stained coronal sections with tetrode tracks in DG and CA3 for each animal (CON, 6 control rats, A–F; LES, 7 DG-lesioned rats, G–M). Tetrode IDs and their positions normalized along the proximodistal axis of CA3 are indicated. Red dots, recording sites. Black arrowhead, CA3/CA2 border. Note that tetrodes located near the distal end may capture neuronal activity from CA2. For rat R1025, TT7 data were acquired ~320 μm dorsal to its final location (pCA3), and likely reflect DG activity; these data were therefore excluded from CA3 analyses. DG, dentate gyrus; pCA3, proximal CA3; mCA3, middle CA3; dCA3, distal CA3; supra, suprapyramidal blade; infra, infrapyramidal blade. Scale bars: 1 mm.

(N) Cumulative distribution of tetrode tip positions along the dorsoventral axis. Distances were calculated relative to the X-shaped intersection (zero point) of dorsal and ventral CA3 pyramidal layers in coronal sections, corresponding to the transition between dorsal and ventral hippocampus. Kolmogorov–Smirnov test comparing control and lesioned distributions: 85 sites, $Z = 0.922$, $P = 0.363$.

n.s., not significant.

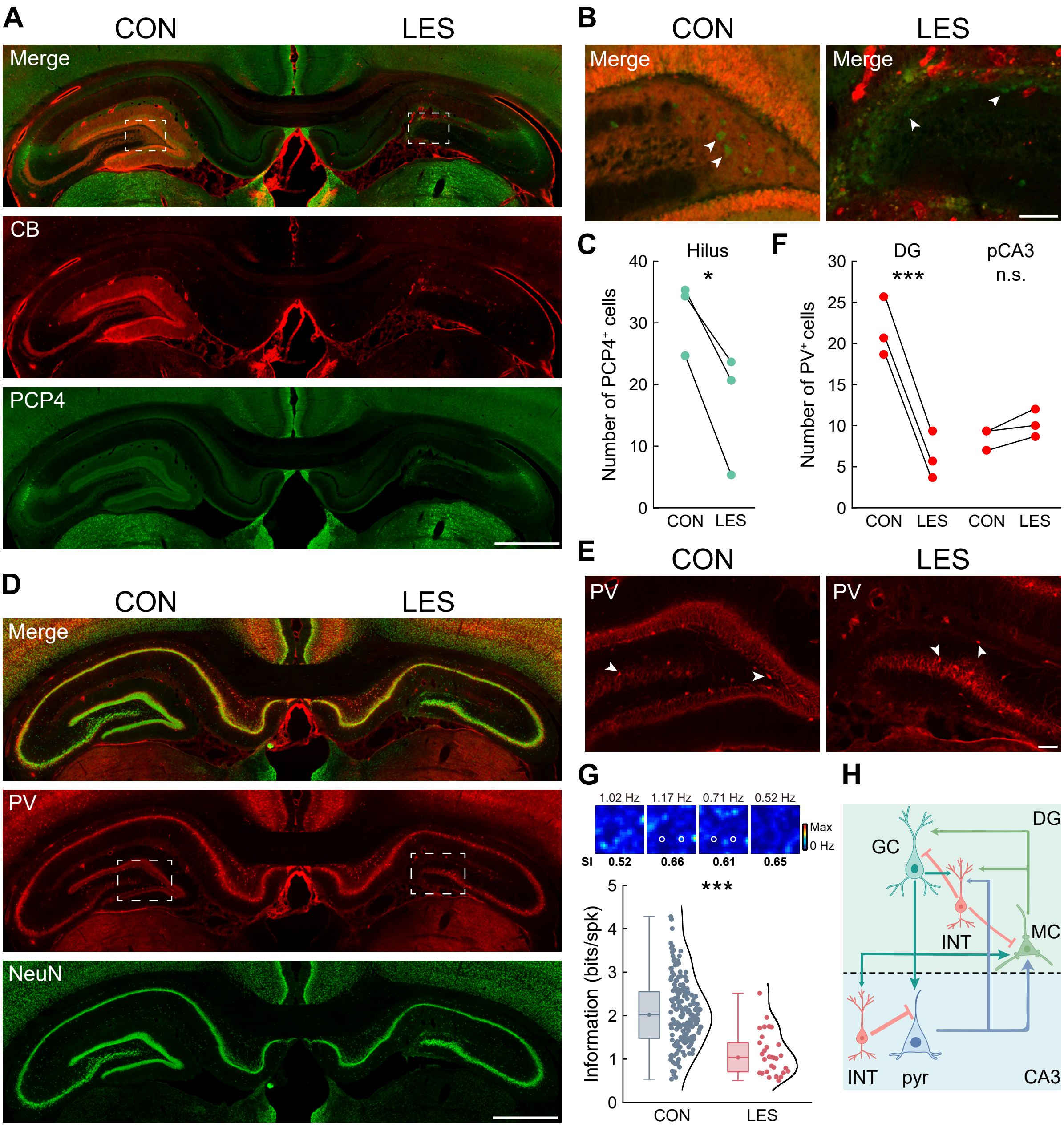
A
CON
LES
Merge
CB
PCP4
B
CON
LES
Merge
Merge
C
Hilus
*
Number of PCP4+ cells
CON LES
F
DG
***
pCA3
n.s.
Number of PV+ cells
CON LES CON LES
D
CON
LES
Merge
PV
NeuN
E
CON
LES
PV
PV
G
1.02 Hz 1.17 Hz 0.71 Hz 0.52 Hz
Max
0 Hz
SI 0.52 0.66 0.61 0.65
***
Information (bits/spk)
CON LES
H
DG
GC
INT
MC
INT
pyr
CA3

**Figure S2. Colchicine depletes mossy cells and PV-interneurons.**

(A) Low-magnification image showing both hippocampi, with the right DG lesioned. CB, calbindin; PCP4, Purkinje cell protein 4.

(B) Magnified views of boxed hilar regions in the left panel. Arrowheads indicate mossy cells (MCs).

(C) MC counts (PCP4+/CB−) per hilus. Three rats, three sections each; dots, individual animals. CON, 31.444 ± 3.401; LES, 16.556 ± 5.678. Paired *t*-test, t = 5.945, *P* = 0.027; Cohen's d = 3.433.

(D–F) Same layout as (A–C), showing parvalbumin (PV)+ interneurons in the hilus and CA3. DG: CON, 21.667 ± 2.082; LES, 6.222 ± 1.659, t = 34.750, *P* = 0.001, Cohen's d = 20.063; CA3: CON, 8.556 ± 0.778; LES, 10.222 ± 0.969, t = −2.887, *P* = 0.102.

(G) Top: example rate map of a residual excitatory neuron in the lesioned DG, with peak rate and spatial information (SI, bits/spike) indicated. Bottom: SI distribution in the DG of both groups. CON: 2.021 (1.479–2.551); LES: 1.036 (0.708–1.374), Wilcoxon rank-sum test, 209 cells, Z = 6.315, *P* < 0.001, $\eta^2$ = 0.191.

(H) Schematic of the local DG-CA3 network (GC, granule cell; MC, mossy cell; INT, interneuron; pyr, pyramidal neuron).

n.s., not significant; *, *P* < 0.05; ***, *P* ≤ 0.001.

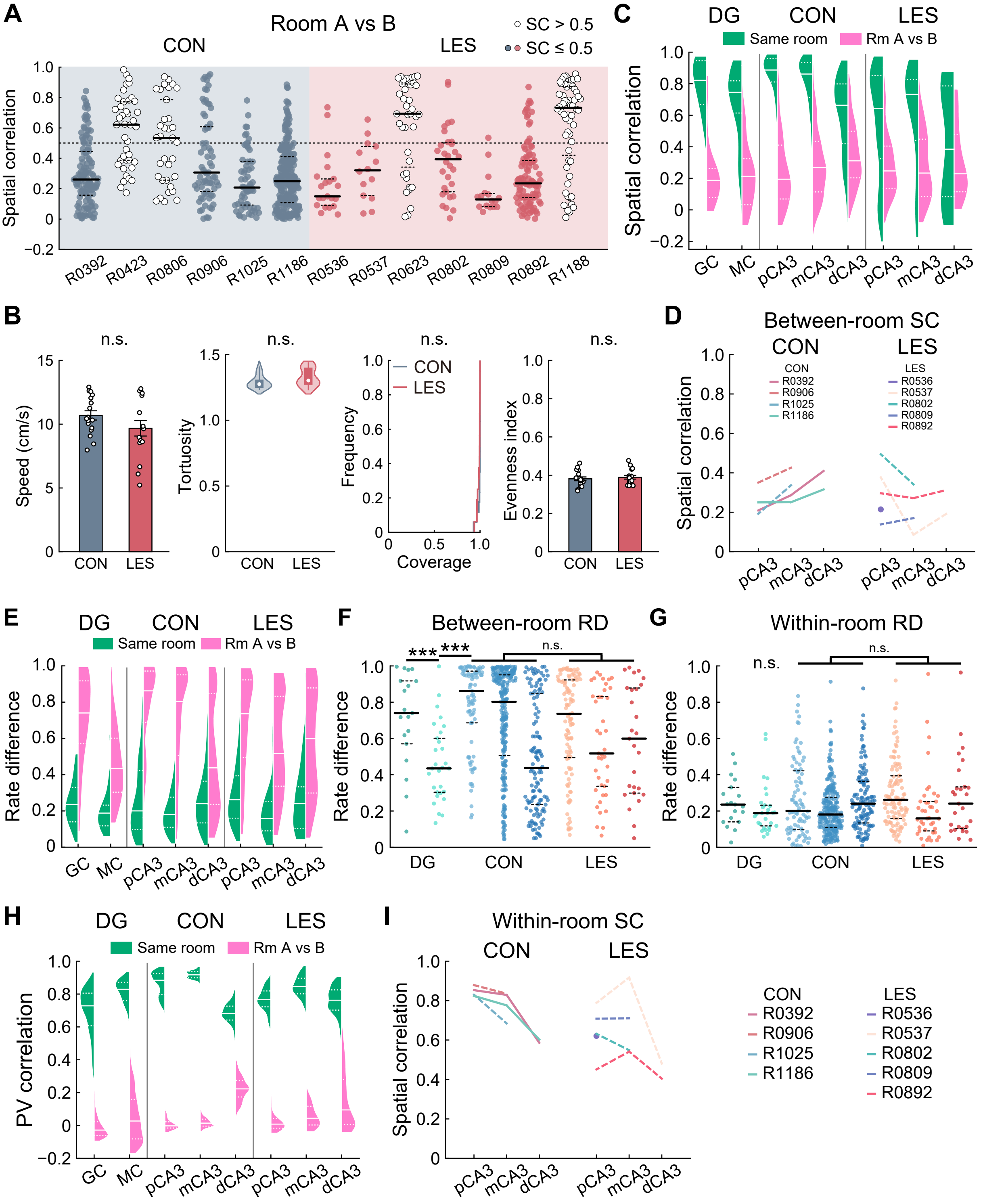

A
Room A vs B
CON
LES
SC > 0.5
SC ≤ 0.5
Spatial correlation
R0392
R0423
R0806
R0906
R1025
R1186
R0536
R0537
R0623
R0802
R0809
R0892
R1188
B
n.s.
Speed (cm/s)
Tortuosity
Frequency
Coverage
Evenness index
CON
LES
C
DG
CON
LES
Same room
Rm A vs B
Spatial correlation
GC
MC
pCA3
mCA3
dCA3
D
Between-room SC
Spatial correlation
E
Rate difference
F
Between-room RD
***
G
Within-room RD
H
PV correlation
I
Within-room SC

**Figure S3. Extended analyses of the two-room task.**

(A) Between-room spatial correlation (SC) of all place cells for each animal. Rats R0423, R0806, R0623, and R1188 showed median correlations > 0.5, indicative of rate remapping, thus were excluded from subsequent analyses. Dots, individual neurons; solid line, median; dashed lines, quartiles.

(B) Behavioral control variables (mean ± SEM or median (IQR)): Speed: CON, 10.690 ± 0.364 cm/s, LES, 9.678 ± 0.608 cm/s, *t*-test, 33 experiments, t = 1.448, $P = 0.158$. Tortuosity: CON, 1.275 (1.250–1.305), LES, 1.303 (1.274–1.397), Wilcoxon rank sum test, Z = −1.657, $P = 0.102$. Coverage: Kolmogorov-Smirnov test, Z = 1.319, $P = 0.062$. Evenness: CON, 0.381 ± 0.010, LES, 0.389 ± 0.010, t = −0.576, $P = 0.569$.

(C) Violin plots showing distributions of SC across DG and CA3 subregions. Solid line, median; dashed lines, quartiles.

(D) Between-room SC for each rat. Colored lines and dots, animal means; dashed lines, < 20 place cells. Trends mirror cell-level comparisons.

(E) Rate difference (RD) distributions across DG and CA3 subregions.

(F) Between-room RD. GC, 0.740 (0.570–0.917); MC, 0.435 (0.303–0.601); CON: pCA3, 0.862 (0.686–0.972); mCA3, 0.802 (0.507–0.951); dCA3, 0.438 (0.236–0.847); LES: pCA3, 0.736 (0.495–0.923); mCA3, 0.517 (0.336–0.832); dCA3, 0.599 (0.299–0.878). GC vs. MC vs. pCA3, Linear mixed-effects model with animal as a random intercept (LMM): 506 cells; group, $F\,(2, 132) = 20.196$, $P < 0.001$; negligible between-animal variance. CON vs. LES, LMM, 634 cells, group, $F\,(1, 3.946) = 0.287$, $P = 0.621$; band, $F\,(2, 385.383) = 12.771$, $P < 0.001$; group × band, $F\,(2, 385.383) = 4.108$, $P = 0.017$; random effect of animal, $P = 0.385$.

(G) Within-room RD. GC, 0.236 (0.140–0.331); MC, 0.188 (0.118–0.232); CON: pCA3, 0.200 (0.097–0.422); mCA3, 0.181 (0.110–0.276); dCA3, 0.240 (0.134–0.364). LES: pCA3, 0.262 (0.160–0.394); mCA3, 0.159 (0.090–0.252); dCA3, 0.241 (0.103–0.333). GC vs. MC vs. pCA3, LMM: 506 cells; group, $F\,(2, 48.357) = 0.583$, $P = 0.562$; random effect of animal, $P = 0.670$. CON vs. LES, LMM, 634 cells, Group, $F\,(1, 9.808) = 1.160$, $P = 0.307$; band, $F\,(2, 418.456) = 7.531$, $P = 0.001$; group × band, $F\,(2, 418.456) = 0.107$, $P = 0.898$; random effect of animal, $P = 0.353$.

(H) Population vector correlation (PVC) distributions across DG and CA3 subregions.

(I) Within-room SC for each rat. Colored lines and dots, animal means; dashed lines, < 20 place cells. Trends mirror cell-level comparisons. n.s., not significant; ***, $P \leq 0.001$.

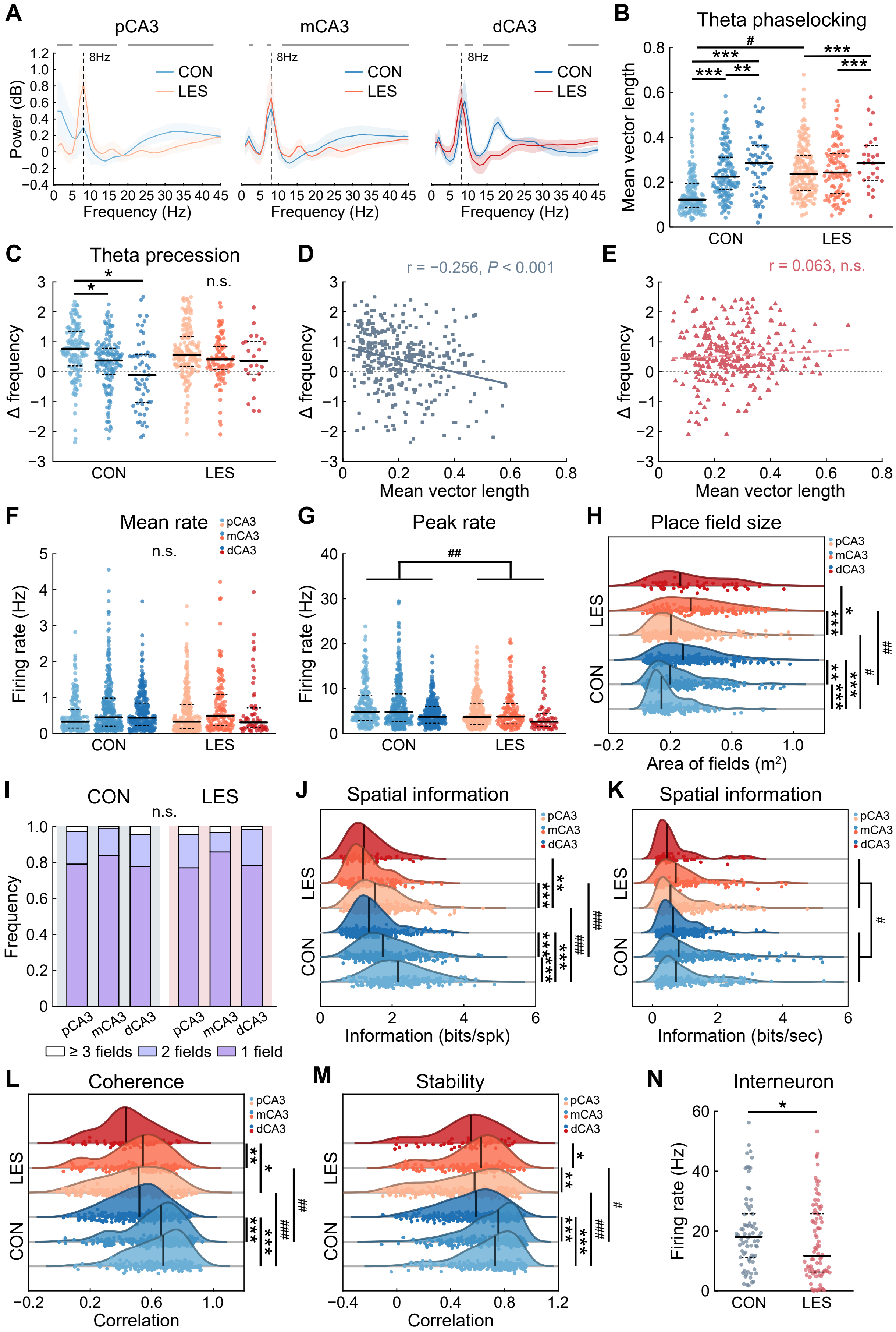
A
pCA3
mCA3
dCA3
8Hz
CON
LES
Power (dB)
Frequency (Hz)
B
Theta phaselocking
Mean vector length
CON
LES
C
Theta precession
Δ frequency
n.s.
D
r = −0.256, P < 0.001
Mean vector length
E
r = 0.063, n.s.
F
Mean rate
n.s.
Firing rate (Hz)
pCA3
mCA3
dCA3
G
Peak rate
##
H
Place field size
Area of fields (m2)
I
Frequency
≥ 3 fields
2 fields
1 field
J
Spatial information
Information (bits/spk)
K
Spatial information
Information (bits/sec)
L
Coherence
Correlation
M
Stability
Correlation
N
Interneuron
Firing rate (Hz)

**Figure S4. DG lesion attenuates CA3 temporal and spatial coding.**

Temporal and spatial activity related measures for each CA3 subregion. n.s., not significant; within-group: *, $P \leq 0.05$; **, $P < 0.01$; ***, $P < 0.001$; between-group: $^{\#}$, $P < 0.05$; $^{\#\#}$, $P < 0.01$; $^{\#\#\#}$, $P \leq 0.001$.

(A) CA3 LFP power spectra without aperiodic components. Solid lines, mean power; shaded areas, standard deviation. Multiple comparisons, FDR corrected. Gray horizontal bars, significant between-group differences. Comparison of peak theta power (8 Hz, dashed vertical lines): two-way ANOVA, 436 experiments, group, $F$ (1, 430) = 26.883, $P < 0.001$, $\eta^2 = 0.059$; region, $F$ (2, 430) = 0.518, $P = 0.596$; group × region, $F$ (2, 430) = 20.881, $P < 0.001$, $\eta^2 = 0.089$. Holm-Bonferroni post hoc tests: CON group, pCA3 < mCA3 ($P < 0.001$), LES group, pCA3 > mCA3 ($P = 0.001$), mCA3 vs. dCA3, n.s. in both groups.

(B) Theta phase-locking strength. Dots, individual neurons; solid line, median; dashed lines, quartiles. CON: pCA3, 0.122 (0.088–0.194); mCA3, 0.225 (0.166–0.311); dCA3, 0.284 (0.175–0.362). LES: pCA3, 0.236 (0.163–0.318); mCA3, 0.243 (0.149–0.327); dCA3, 0.284 (0.209–0.362). LMM: 726 cells, group, $F$ (1, 8.835) = 2.476, $P = 0.151$; band, $F$ (2, 711.781) = 20.110, $P < 0.001$; group × band, $F$ (2, 711.781) = 6.423, $P = 0.002$; random effect of animal, $P = 0.066$.

(C) Strength of theta phase precession, expressed as the increase in intrinsic theta frequency. CON: pCA3, 0.770 (0.194–1.351); mCA3, 0.375 (−0.102–0.785); dCA3, −0.114 (−1.021–0.569). LES: pCA3, 0.556 (0.182–1.182); mCA3, 0.409 (0.077–0.838); dCA3, 0.363 (−0.077–1.004). LMM: 654 cells, group, $F$ (1, 10.489) = 0.102, $P = 0.756$; band, $F$ (2, 514.006) = 6.485, $P = 0.002$; group × band, $F$ (2, 514.006) = 0.382, $P = 0.683$; random effect of animal, $P = 0.134$.

(D–E) Correlation between theta phase-locking strength and phase precession strength in control (D) and lesioned (E) groups. Squares and triangles represent individual neurons. Solid regression lines indicate significant correlations; dashed lines indicate non-significant correlations. A significant negative correlation was observed in control rats, which was degraded in lesioned rats, indicating an abolished gradient in the lesioned group (permutation test, $P < 0.001$).

(F) Mean firing rates. Dots indicate individual neurons; solid line, median; dashed lines, quartiles. CON: pCA3, 0.328 (0.153–0.672) Hz; mCA3, 0.449 (0.206–0.987) Hz; dCA3, 0.443 (0.223–0.847) Hz. LES: pCA3, 0.329 (0.144–0.813) Hz; mCA3, 0.497 (0.229–1.092) Hz; dCA3, 0.309 (0.160–0.715) Hz. LMM: 1 319 cells, group, $F$

(1, 11.232) = 0.186, $P$ = 0.674; band, $F$ (2, 915.716) = 17.615, $P$ < 0.001; group × band, $F$ (2, 915.716) = 0.424, $P$ = 0.655; random effect of animal, $P$ = 0.096.

(G) Peak firing rates. CON: pCA3, 4.834 (2.970–8.395) Hz; mCA3, 4.795 (2.657–8.830) Hz; dCA3, 3.761 (2.309–6.033) Hz. LES: pCA3, 3.660 (2.067–6.773) Hz; mCA3, 3.796 (2.151–6.651) Hz; and dCA3, 2.641 (1.592–4.389) Hz. LMM: 1 319 cells, group, $F$ (1, 8.379) = 11.587, $P$ = 0.009; band, $F$ (2, 366.825) = 12.893, $P$ < 0.001; group × band, $F$ (2, 366.825) = 0.173, $P$ = 0.841; random effect of animal, $P$ = 0.368.

(H) Place-field sizes (ridgeline plots). CON: pCA3, 0.143 (0.095–0.238) $m^2$; mCA3, 0.198 (0.118–0.340) $m^2$; dCA3, 0.281 (0.170–0.428) $m^2$. LES: pCA3, 0.204 (0.120–0.335) $m^2$; mCA3, 0.333 (0.175–0.499) $m^2$; dCA3, 0.265 (0.150–0.513) $m^2$. LMM: 1 319 cells, group, $F$ (1, 10.663) = 4.249, $P$ = 0.065; band, $F$ (2, 1 086.393) = 36.403, $P$ < 0.001; group × band, $F$ (2, 1 086.393) = 4.862, $P$ = 0.008; random effect of animal, $P$ = 0.074.

(I) Number of place fields per cell. CON: proportion of one-field place cells, pCA3, 0.791; mCA3, 0.838; dCA3, 0.779. LES: pCA3, 0.770; mCA3, 0.858; dCA3, 0.783. LMM: 1 319 cells, group, $F$ (1, 7.724) = 0.008, $P$ = 0.933; band, $F$ (2, 204.392) = 3.486, $P$ = 0.032; group × band, $F$ (2, 204.392) = 0.539, $P$ = 0.584; random effect of animal, $P$ = 0.760.

(J) Spatial information content. CON: pCA3, 2.165 (1.640–2.714) bits/spike; mCA3, 1.740 (1.303–2.320) bits/spike; dCA3, 1.355 (1.075–1.742) bits/spike. LES: pCA3, 1.527 (1.164–2.057) bits/spike; mCA3, 1.186 (0.906–1.626) bits/spike; dCA3, 1.220 (0.918–1.587) bits/spike. LMM: 1 319 cells, group, $F$ (1, 11.341) = 18.202, $P$ = 0.001; band, $F$ (2, 1 038.782) = 52.954, $P$ < 0.001; group × band, $F$ (2, 1 038.782) = 7.028, $P$ = 0.001; animal, $P$ = 0.074.

(K) Spatial information rate. CON: pCA3, 0.717 (0.437–1.372) bits/second; mCA3, 0.802 (0.422–1.652) bits/second; dCA3, 0.628 (0.368–1.020) bits/second. LES: pCA3, 0.554 (0.296–1.082) bits/second; mCA3, 0.713 (0.342–1.179) bits/second; dCA3, 0.449 (0.243–0.705) bits/second. LMM: 1 319 cells, group, $F$ (1, 9.514) = 5.622, $P$ = 0.040; band, $F$ (2, 531.534) = 12.195, $P$ < 0.001; group × band, $F$ (2, 531.534) = 0.105, $P$ = 0.900; random effect of animal, $P$ = 0.234.

(L) Coherence. CON: pCA3, 0.675 (0.526–0.782); mCA3, 0.660 (0.526–0.749); dCA3, 0.520 (0.386–0.628). LES: pCA3, 0.514 (0.336–0.656); mCA3, 0.542 (0.392–0.626); dCA3, 0.431 (0.342–0.569). LMM: 1 319 cells; group, $F$ (1, 11.596) = 16.636, $P$ = 0.002; band, $F$ (2, 991.414) = 25.300, $P$ < 0.001; group × band, $F$ (2, 991.414) = 3.016, $P$ = 0.049; animal, $P$ = 0.079.

(M) Stability. CON: pCA3, 0.728 (0.573–0.846); mCA3, 0.754 (0.610–0.855); dCA3, 0.588 (0.406–0.717). LES: pCA3, 0.577 (0.342–0.749); mCA3, 0.626 (0.461–0.724); dCA3, 0.552 (0.400–0.692). LMM: 1 317 cells, group, $F$ (1, 12.095) = 7.068, $P$ = 0.021; band, $F$ (2, 1 203.049) = 18.407, $P$ < 0.001; group × band, $F$ (2, 1 203.049) = 5.276, $P$ = 0.005; random effect of animal, $P$ = 0.043.

(N) Mean firing rate of CA3 interneurons (Hz). CON, 18.017 (11.053–25.704); LES, 11.742 (6.294–25.727). Wilcoxon rank-sum test, 162 cells, Z = 2.071, $P$ = 0.038, $\eta^2$ = 0.026.

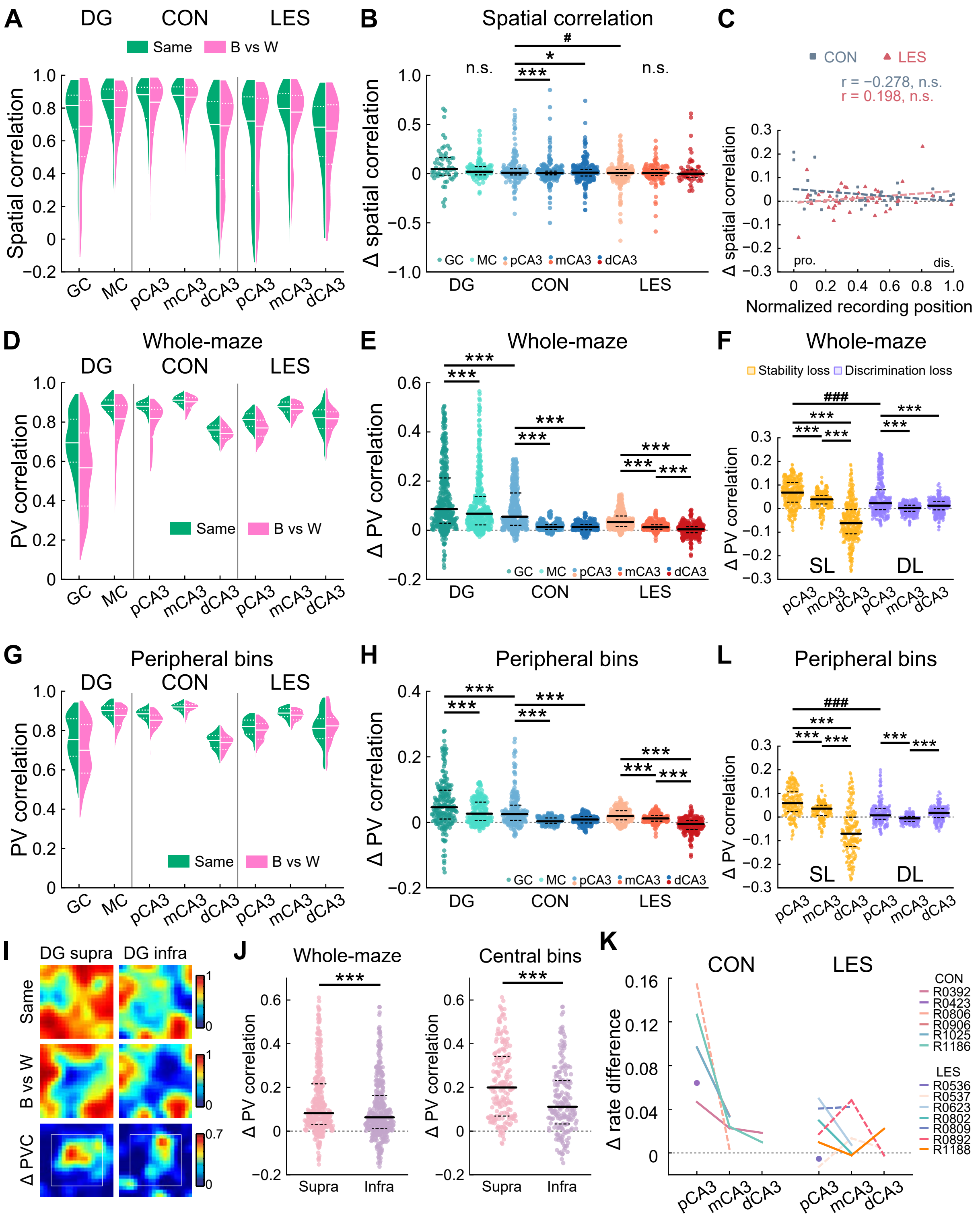
A
DG
CON
LES
Same
B vs W
Spatial correlation
GC
MC
pCA3
mCA3
dCA3
B
Spatial correlation
Δ spatial correlation
n.s.
#
*
***
GC
MC
pCA3
mCA3
dCA3
DG
CON
LES
C
CON
LES
r = −0.278, n.s.
r = 0.198, n.s.
Δ spatial correlation
pro.
dis.
Normalized recording position
D
Whole-maze
PV correlation
E
Whole-maze
Δ PV correlation
F
Whole-maze
Stability loss
Discrimination loss
###
SL
DL
G
Peripheral bins
H
Peripheral bins
L
Peripheral bins
I
DG supra
DG infra
Same
B vs W
Δ PVC
J
Whole-maze
Central bins
Supra
Infra
K
Δ rate difference
CON
R0392
R0423
R0806
R0906
R1025
R1186
LES
R0536
R0537
R0623
R0802
R0809
R0892
R1188

**Figure S5. Extended analyses of the color-reversal task.**

(A) SC distributions across DG and CA3 subregions.

(B) Color-reversal ΔSC (reversed-color SC − same-color SC). DG: GC, 0.047 (−0.015–0.164); MC, 0.020 (−0.012–0.071). CON: pCA3, 0.009 (−0.013–0.050); mCA3, 0.006 (−0.012–0.026); dCA3, 0.010 (−0.024–0.043). LES: pCA3, 0.008 (−0.022–0.041); mCA3, 0.006 (−0.017–0.043); dCA3, 0 (−0.037–0.038). GC vs. MC vs. pCA3, LMM: 398 cells, group, $F$ (2, 310.811) = 2.068, $P$ = 0.128; random effect of animal, $P$ = 0.227. CON vs. LES, LMM: 1 310 cells, group, $F$ (1, 8.488) = 1.174, $P$ = 0.308; band, $F$ (2, 905.465) = 4.155, $P$ = 0.016; group × band, $F$ (2, 905.465) = 3.161, $P$ = 0.043; random effect of animal, $P$ = 0.133.

(C) Correlation between CA3 recording position and mean color-reversal ΔSC (CON, 39 tetrodes; LES, 45 tetrodes). The control and lesion groups showed similar ΔSC between same-color and color-reversed pairs (Wilcoxon rank-sum test, Z = 1.108, $P$ = 0.268). No significant transverse gradients were observed in either group, nor was there a significant difference in gradient slope between groups (permutation test, $P$ = 0.061).

(D) Violin plots showing PVC distributions for GCs, MCs, and CA3 neurons in control and DG lesioned rats. Solid line, median; dashed lines, quartiles.

(E) Color-reversal ΔPVC (same-color PVC − reversed-color PVC). Dots, spatial bins. DG: GC, 0.086 (0.029–0.213); MC, 0.067 (0.022–0.137). CON: pCA3, 0.055 (0.020–0.152); mCA3, 0.013 (0.003–0.022); dCA3, 0.014 (0.002–0.024). LES: pCA3, 0.033 (0.015–0.058); mCA3, 0.012 (0.004–0.022); dCA3, 0.004 (−0.010–0.014). GC vs. MC vs. pCA3, Friedman test, 400 bins, $\chi^2$ (2) = 64.655, $P$ < 0.001, Kendall's W = 0.081. CON vs. LES, Repeated-measure two-way ANOVA (rmANOVA), 400 bins, group, $F$ (1, 1 197) = 207.596, $P$ < 0.001, $\eta^2$ = 0.148; band, $F$ (2, 1 197) = 331.902, $P$ < 0.001, $\eta^2$ = 0.357; group × band, $F$ (2, 1 197) = 102.536, $P$ < 0.001, $\eta^2$ = 0.146.

(F) Stability loss (SL, Δ same-color PVC between control and lesioned groups, left) and discrimination loss (DL, Δ same-color − Δ color-reversal PVC between groups, right) across CA3 bands. SL: pCA3, 0.068 (0.027–0.111); mCA3, 0.039 (0.020–0.056); dCA3, −0.062 (−0.107–−0.004). DL: pCA3, 0.023 (−0.004–0.080); mCA3, 0.002 (−0.011–0.014); dCA3, 0.012 (−0.005–0.031). rmANOVA, 400bins, group, $F$ (1, 1 197) = 4.548, $P$ = 0.033, $\eta^2$ = 0.004; band, $F$ (2, 1 197) = 348.236, $P$ < 0.001, $\eta^2$ = 0.368; group × band, $F$ (2, 1 197) = 275.907, $P$ < 0.001, $\eta^2$ = 0.316.

(G) Distributions of PVC for peripheral bins across DG and CA3 subregions.

(H) Color-reversal ΔPVC (same-color PVC − reversed-color PVC) for peripheral bins. Dots, spatial bins. DG: GC, 0.046 (0.010–0.098); MC, 0.027 (0.005–0.062). CON: pCA3, 0.025 (0.006–0.053); mCA3, 0.003 (−0.003–0.013); dCA3, 0.009 (−0.002–0.018). LES: pCA3, 0.019 (0.008–0.035); mCA3, 0.012 (0.004–0.021); dCA3, −0.005 (−0.021–0.005). GC vs. MC vs. pCA3, Friedman test, 204 bins, $\chi^2$ (2) = 49.324, $P < 0.001$, Kendall's W = 0.121. CON vs. LES, rmANOVA, 204 bins, group, $F$ (1, 609) = 38.401, $P < 0.001$, $\eta^2 = 0.059$; band, $F$ (2, 609) = 117.687, $P < 0.001$, $\eta^2 = 0.279$; group × band, $F$ (2, 609) = 34.380, $P < 0.001$, $\eta^2 = 0.101$.

(I) PVC heatmaps for GCs in supra- vs infrapyramidal blades. Boxed region: maze center.

(J) ΔPVC (same-color PVC − reversed-color PVC) for supra- and infrapyramidal GCs. Left: whole-maze, supra, 187 cells, 0.082 (0.029–0.216); Infra, 119 cells, 0.063 (0.011–0.162); Wilcoxon signed-rank test, 400 bins, Z = 4.110, $P < 0.001$, $\eta^2 = 0.042$. Right: central bins, supra, 0.199 (0.068–0.341); Infra, 0.111 (0.032–0.230); 196 bins, Z = 5.375, $P < 0.001$, $\eta^2 = 0.147$.

(K) Color-reversal ΔRD (reversed-color RD − same-color RD) per rat. Colored lines and dots, animal means; dashed lines, < 20 place cells. Trends match pooled-cell analyses.

(L) SL and DL for peripheral bins across CA3 bands. SL: pCA3, 0.059 (0.023–0.107); mCA3, 0.035 (0.007–0.050); dCA3, −0.071 (−0.124–−0.001). DL: pCA3, 0.007 (−0.010–0.035); mCA3, −0.006 (−0.019–0.002); dCA3, 0.017 (−0.002–0.035). rmANOVA, 204 bins, group, $F$ (1, 609) = 0.181, $P = 0.671$; band, $F$ (2, 609) = 117.305, $P < 0.001$, $\eta^2 = 0.278$; group × band, $F$ (2, 609) = 207.918, $P < 0.001$, $\eta^2 = 0.406$.

n.s., not significant; within-group: *, $P < 0.05$; ***, $P \leq 0.001$; between-group: $^{\#}$, $P < 0.05$; $^{\#\#\#}$, $P < 0.001$.

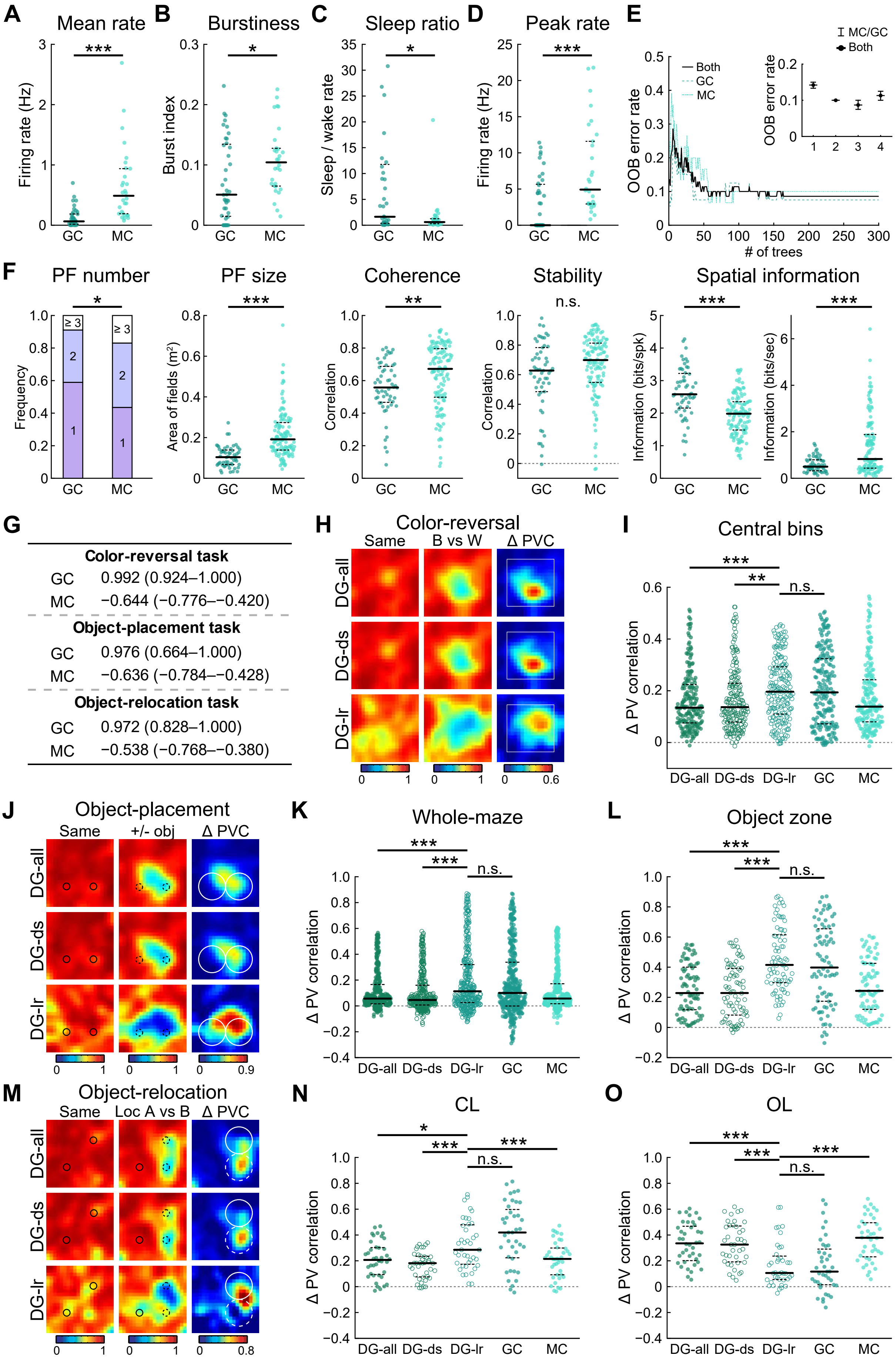
A
Mean rate
Firing rate (Hz)
***
GC
MC
B
Burstiness
Burst index
*
C
Sleep ratio
Sleep / wake rate
*
D
Peak rate
Firing rate (Hz)
***
E
OOB error rate
### of trees
Both
GC
MC
MC/GC
Both
OOB error rate
F
PF number
Frequency
≥ 3
PF size
Area of fields (m²)
Coherence
Correlation
**
Stability
n.s.
Spatial information
Information (bits/spk)
Information (bits/sec)
G
Color-reversal task
GC 0.992 (0.924–1.000)
MC −0.644 (−0.776–−0.420)
Object-placement task
GC 0.976 (0.664–1.000)
MC −0.636 (−0.784–−0.428)
Object-relocation task
GC 0.972 (0.828–1.000)
MC −0.538 (−0.768–−0.380)
H
Color-reversal
Same
B vs W
Δ PVC
DG-all
DG-ds
DG-lr
I
Central bins
Δ PV correlation
J
Object-placement
Same
+/- obj
Δ PVC
K
Whole-maze
L
Object zone
M
Object-relocation
Same
Loc A vs B
Δ PVC
N
CL
O
OL

**Figure S6. Training and performance of the random-forest classifier for GCs vs MCs, and validates sparse GCs as primary non-spatial rate coders.**

(A–D) Electrophysiological features used to train the random-forest classifier. Dots, spatial bins; solid line, median; dashed lines, quartiles. Wilcoxon rank-sum test, 70 cells: mean rate (A), GC, 0.064 (0.015–0.187); MC, 0.488 (0.185–0.940); Z = −5.459, $P < 0.001$, $\eta^2 = 0.426$; burstiness (B), GC, 0.051 (0.014–0.135); MC, 0.104 (0.065–0.131); Z = −2.447, $P = 0.014$, $\eta^2 = 0.086$; sleep ratio (C), GC, 1.631 (0.352–11.980); MC, 0.601 (0.277–1.337); Z = 2.350, $P = 0.019$, $\eta^2 = 0.079$; peak rate (D), GC, 0 (0–5.712); MC, 4.923 (2.930–11.829); Z = −3.609, $P < 0.001$, $\eta^2 = 0.186$.

(E) Out-of-bag (OOB) error as a function of decision tree numbers. Inset: OOB error vs. number of features tried at each split; solid circles mark the optimum (300 trees, 3 random features).

(F) Spatial firing metrics between GCs and MCs (180 cells). Place fields per cell (1: 1 field; 2: 2 fields; ≥ 3: ≥ 3 fields): GC, 1.00 (1.00–2.00); MC, 2.00 (1.00–2.00); Wilcoxon rank sum test, Z = −2.006, $P = 0.045$, $\eta^2 = 0.022$. Place field size: GC, 0.104 (0.068–0.141) $m^2$, MC, 0.191 (0.138–0.276) $m^2$, Z = −7.200, $P < 0.001$, $\eta^2 = 0.288$. Coherence: GC, 0.558 (0.465–0.691), MC, 0.672 (0.498–0.797), Z = −3.019, $P = 0.003$, $\eta^2 = 0.051$. Stability: GC, 0.628 (0.485–0.783), MC, 0.699 (0.547–0.814), Z = −1.788, $P = 0.074$. Spatial information content: GC, 2.579 (2.143–3.229) bits/spike, MC, 1.984 (1.482–2.358) bits/spike, Z = 5.599, $P < 0.001$, $\eta^2 = 0.174$. Spatial information rate: GC, 0.500 (0.319–0.797) bits/second, MC, 0.829 (0.436–1.892) bits/second, Z = −3.955, $P < 0.001$, $\eta^2 = 0.087$.

(G) Random-forest classifier confidence for dentate cells across three tasks. Note: values approaching ±1 reflect maximal certainty; values near 0 indicate ambiguity.

(H) PVC heatmaps for all DG neurons (DG-all), downsampled DG (DG-ds), low-rate subset (mean rate < 0.5 Hz; DG-lr) in the color-reversal task. Left columns show neuron populations across repeated and color-reversed trials; right column shows ΔPVC. Boxed region: maze center.

(I) Color-reversal ΔPVC (same-color PVC − reversed-color PVC) for central bins. DG-all, 0.135 (0.076–0.224); DG-ds, 0.137 (0.079–0.229); DG-lr, 0.196 (0.110–0.293); GC, 0.194 (0.072–0.324); MC, 0.139 (0.080–0.242). rmANOVA, region × color: DG-all vs. DG-ds vs. DG-lr, 196 bins, $F(1, 585) = 6.859$, $P = 0.001$, $\eta^2 = 0.023$; DG-lr vs. GC, $F(1, 390) = 0.036$, $P = 0.849$. Similar results were obtained for the whole-maze: DG-all, 0.060 (0.020–0.133); DG-ds, 0.060 (0.019–0.131); DG-lr, 0.085 (0.027–0.201); GC, 0.086 (0.029–0.213); MC, 0.067 (0.022–

0.137). Region × color: DG-all vs. DG-ds vs. DG-lr, 400 bins, $F$ (1, 1 197) = 6.770, $P$ = 0.001, $\eta^2$ = 0.011; DG-lr vs. GC, $F$ (1, 798) = 0.671, $P$ = 0.413. Excluding high-rate, MC-like neurons yielded equivalent discrimination in the remaining GC-dominated population, confirming classifier robustness.

(J) PVC heatmaps for all DG neurons and subsets in the object-placement task: left columns compare repeated trials and trials with/without objects, right column shows ΔPVC. Small circle: object; large circle: object zone.

(K) Object-induced ΔPVC (whole-maze). DG-all, 0.057 (0.016–0.166); DG-ds, 0.046 (0.007–0.160); DG-lr, 0.113 (0.026–0.320); GC, 0.100 (−0.001–0.338); MC, 0.057 (0.017–0.172). rmANOVA, region × object: DG-all vs. DG-ds vs. DG-lr, 400 bins, $F$ (1, 1 197) = 30.504, $P$ < 0.001, $\eta^2$ = 0.048; DG-lr vs GC: $F$ (1, 798) = 1.318, $P$ = 0.251.

(L) Object-zone ΔPVC. DG-all, 0.228 (0.118–0.400); DG-ds, 0.228 (0.082–0.392); DG-lr, 0.415 (0.297–0.615); GC, 0.398 (0.174–0.655); MC, 0.243 (0.120–0.425). Region × object: DG-all vs. DG-ds vs. DG-lr, 80 bins, $F$ (1, 237) = 32.449, $P$ < 0.001, $\eta^2$ = 0.215; DG-lr vs GC: $F$ (1, 158) = 0.905, $P$ = 0.343.

(M) PVC heatmaps for all DG neurons and subsets in the object-relocation task. Left columns: repeated trials vs pre-/post-relocation; right column: ΔPVC. Small circle: object; large solid: current location (CL) zone; large dashed: original location (OL) zone.

(N) CL-zone ΔPVC. DG-all, 0.207 (0.093–0.302); DG-ds, 0.182 (0.076–0.235); DG-lr, 0.285 (0.174–0.479); GC, 0.419 (0.224–0.598); MC, 0.215 (0.092–0.299). rmANOVA, region × relocation: DG-all vs. DG-ds vs. DG-lr, 40 bins, $F$ (1, 117) = 11.981, $P$ < 0.001, $\eta^2$ = 0.170; DG-lr vs. GC, $F$ (1, 78) = 2.662, $P$ = 0.107; DG-lr vs. MC, 40 bins, $F$ (1, 78) = 9.301, $P$ = 0.003, $\eta^2$ = 0.107.

(O) OL-zone ΔPVC. DG-all, 0.335 (0.202–0.468); DG-ds, 0.326 (0.192–0.470); DG-lr, 0.106 (0.055–0.237); GC, 0.116 (0.015–0.291); MC, 0.378 (0.231–0.495). Region × relocation: DG-all vs. DG-ds vs. DG-lr, 40 bins, $F$ (1, 117) = 12.926, $P$ < 0.001, $\eta^2$ = 0.181; DG-lr vs. GC, $F$ (1, 78) = 0.046, $P$ = 0.831; DG-lr vs. MC, 40 bins, $F$ (1, 78) = 26.248, $P$ < 0.001, $\eta^2$ = 0.252.

n.s., not significant; *, $P$ < 0.05; **, $P$ < 0.01; ***, $P$ < 0.001.

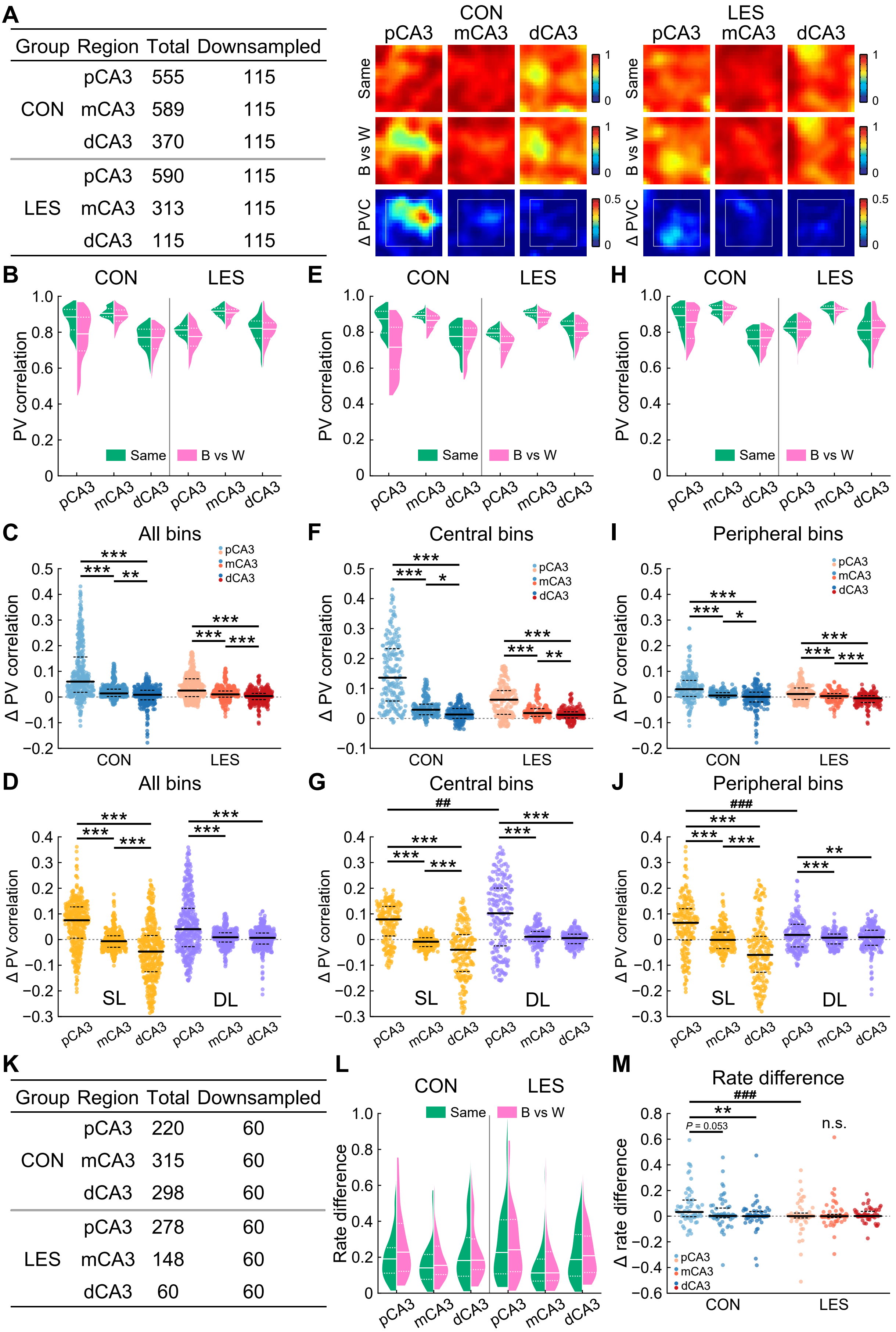

A
Group Region Total Downsampled
CON pCA3 555 115
mCA3 589 115
dCA3 370 115
LES pCA3 590 115
mCA3 313 115
dCA3 115 115
CON
LES
pCA3 mCA3 dCA3
Same
B vs W
Δ PVC
B
E
H
CON
LES
PV correlation
Same
B vs W
C
All bins
F
Central bins
I
Peripheral bins
Δ PV correlation
pCA3
mCA3
dCA3
D
G
J
SL
DL
##
###
K
Group Region Total Downsampled
CON pCA3 220 60
mCA3 315 60
dCA3 298 60
LES pCA3 278 60
mCA3 148 60
dCA3 60 60
L
Rate difference
M
Rate difference
P = 0.053
n.s.
Δ rate difference

**Figure S7. Color-reversal discrimination remains robust following cell downsampling.**

(A) Cell counts before and after downsampling (left), and PVC heatmaps for control (CON) and lesioned (LES) groups after downsampling (right).

(B) Distributions of PVC for all spatial bins across CA3 subregions. Solid line, median; dashed lines, quartiles.

(C) Color-reversal ΔPVC (same-color PVC − reversed-color PVC) computed across all spatial bins. CON: pCA3, 0.060 (0.018–0.156); mCA3, 0.014 (0.002–0.031); dCA3, 0.009 (−0.011–0.026). LES: pCA3, 0.025 (0.002–0.071); mCA3, 0.010 (0–0.024); dCA3, 0.004 (−0.010–0.014). rmANOVA, 400 bins, group, $F$ (1, 1 197) = 123.622, $P < 0.001$, $\eta^2 = 0.094$; band, $F$ (2, 1 197) = 278.462, $P < 0.001$, $\eta^2 = 0.318$; group × band, $F$ (2, 1 197) = 68.259, $P < 0.001$, $\eta^2 = 0.102$. Rate-remapping induced discrimination was enhanced in the maze center, resulting in a relatively low overall discrimination when all spatial bins were included.

(D) Stability loss (SL, difference in same-color PVC between control and lesioned groups, left) and discrimination loss (DL, difference in color-reversal ΔPVC between groups, right) across CA3 bands for all bins. SL: pCA3, 0.075 (0.005–0.127); mCA3, −0.007 (−0.030–0.015); dCA3, −0.047 (−0.125–−0.016). DL: pCA3, 0.040 (−0.028–0.121); mCA3, 0.009 (−0.010–0.026); dCA3, 0.007 (−0.017–0.025). rmANOVA, 400 bins, group, $F$ (1, 1 197) = 36.151, $P < 0.001$, $\eta^2 = 0.029$; band, $F$ (2, 1 197) = 241.234, $P < 0.001$, $\eta^2 = 0.287$; group × band, $F$ (2, 1 197) = 27.348, $P < 0.001$, $\eta^2 = 0.044$.

(E–G) Same as (B–D), but for central bins. Color-reversal ΔPVC: CON, pCA3, 0.136 (0.058–0.233); mCA3, 0.029 (0.012–0.048); dCA3, 0.013 (0–0.033). LES, pCA3, 0.062 (0.013–0.093); mCA3, 0.018 (0.007–0.033); dCA3, 0.012 (0.001–0.022). rmANOVA, 196 bins, group, $F$ (1, 585) = 108.039, $P < 0.001$, $\eta^2 = 0.156$; band, $F$ (2, 585) = 409.944, $P < 0.001$, $\eta^2 = 0.584$; group × band, $F$ (2, 585) = 68.291, $P < 0.001$, $\eta^2 = 0.189$. SL vs. DL: SL, pCA3, 0.078(0.014–0.129); mCA3, −0.008 (−0.027–0.007); dCA3, −0.040 (−0.125–0.019). DL, pCA3, 0.102 (−0.025–0.201); mCA3, 0.011 (−0.007–0.032); dCA3, 0.006 (−0.016–0.021). 196 bins, group, $F$ (1, 585) = 48.375, $P < 0.001$, $\eta^2 = 0.076$; band, $F$ (2, 585) = 210.094, $P < 0.001$, $\eta^2 = 0.418$; group × band, $F$ (2, 585) = 3.887, $P = 0.021$, $\eta^2 = 0.013$. Central bins contributed the majority of DL and exceeded SL in pCA3.

(H–J) As (B–D), but for peripheral bins. Color-reversal ΔPVC: CON, pCA3, 0.030 (0.002–0.065); mCA3, 0.006 (−0.002–0.017); dCA3, 0.001 (−0.019–0.019). LES, pCA3, 0.012 (−0.009–0.035); mCA3, 0.004 (−0.005–0.013); dCA3, −0.005 (−0.021–0.005). rmANOVA, 204 bins, group, $F$ (1, 609) = 27.229, $P < 0.001$, $\eta^2 = 0.043$; band, $F$ (2, 609) = 93.202, $P < 0.001$, $\eta^2 = 0.234$; group × band, $F$ (2, 609) = 8.969, $P < 0.001$, $\eta^2 = 0.029$. SL vs. DL:

SL, pCA3, 0.065 (−0.002–0.120); mCA3, −0.001 (−0.035–0.029); dCA3, −0.060 (−0.127–0.012). DL, pCA3, 0.018 (−0.029–0.059); mCA3, 0.007 (−0.017–0.021); dCA3, 0.009 (−0.022–0.036). 204 bins, group, $F$ (1, 609) = 2.189, $P$ = 0.140; band, $F$ (2, 609) = 67.822, $P < 0.001$, $\eta^2 = 0.182$; group × band, $F$ (2, 609) = 34.658, $P < 0.001$, $\eta^2 = 0.102$. Peripheral bins showed negligible changes in discrimination following DG lesion.

(K) Cell counts before and after downsampling in the RD analysis.

(L) RD distributions for CA3 subregions in both control and lesion groups.

(M) Color-reversal ΔRD (reversed-color RD − same-color RD). Dots, individual neurons. CON, pCA3, 0.033 (−0.004–0.125); mCA3, 0.002 (−0.012–0.063); dCA3, 0 (−0.005–0.036). LES, pCA3, 0 (−0.014–0.024); mCA3, 0 (−0.011–0.015); dCA3, 0.001 (−0.002–0.037). LMM, group, $F$ (1, 354) = 4.681, $P$ = 0.031; band, $F$ (2, 354) = 2.511, $P$ = 0.083; group × band, $F$ (2, 354) = 3.876, $P$ = 0.022; between-animal variance was negligible.

n.s., not significant; within-group: *, $P < 0.05$; **, $P < 0.01$; ***, $P < 0.001$; between-group: $^{##}$, $P < 0.01$; $^{###}$, $P \leq 0.001$.

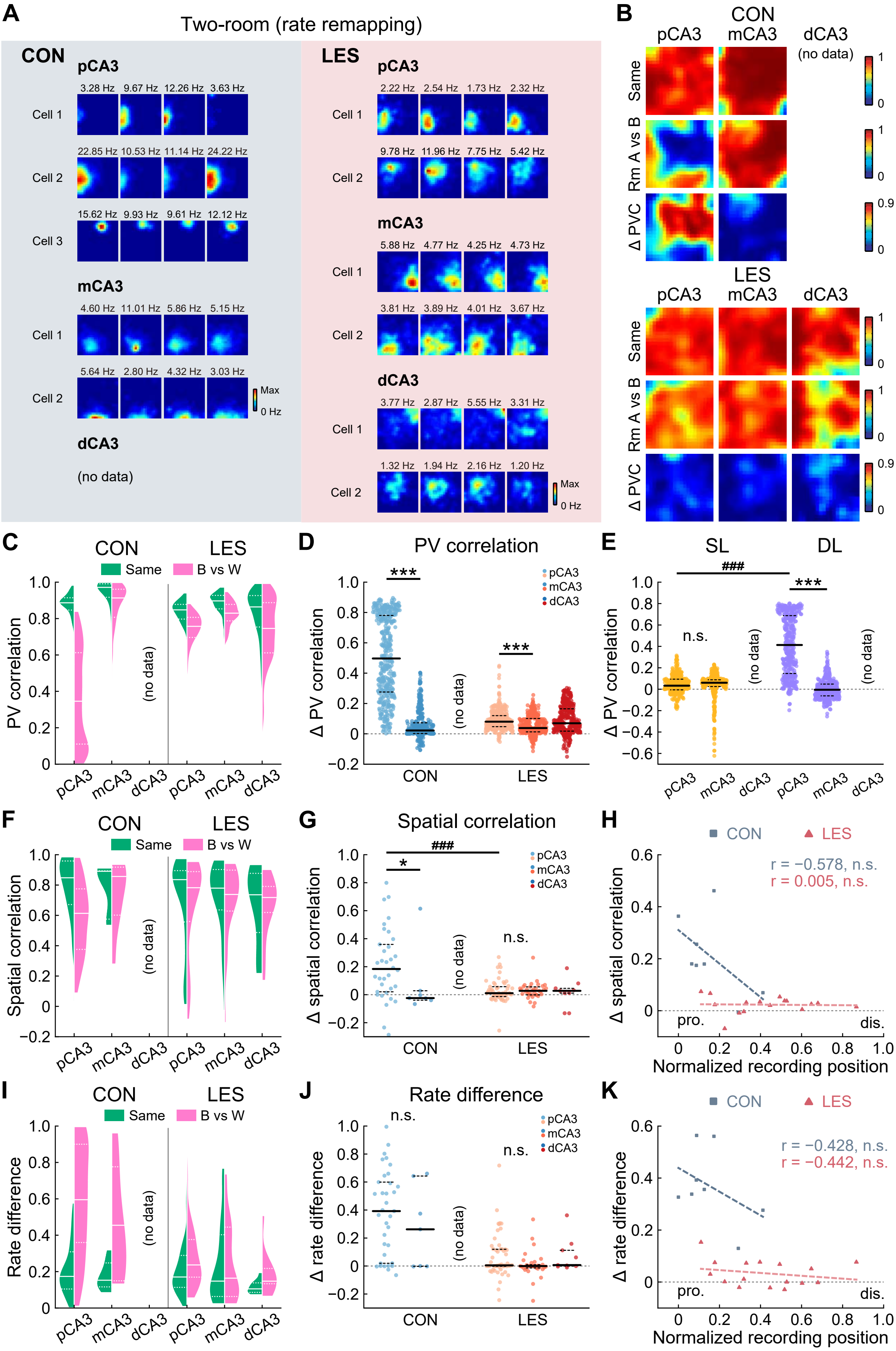
A
Two-room (rate remapping)
CON
pCA3
Cell 1
Cell 2
Cell 3
mCA3
dCA3
(no data)
LES
B
CON
pCA3
mCA3
dCA3
Same
Rm A vs B
Δ PVC
C
Same
B vs W
PV correlation
D
PV correlation
Δ PV correlation
E
SL
DL
F
Spatial correlation
G
Δ spatial correlation
H
Normalized recording position
r = −0.578, n.s.
r = 0.005, n.s.
I
Rate difference
J
Δ rate difference
K
r = −0.428, n.s.
r = −0.442, n.s.
pro.
dis.

**Figure S8. Analyses of the four rate-remapped rats in the two-room task.**

(A) Representative rate maps in control and DG-lesioned rats showing rate remapping across rooms.

(B) PVC heatmaps for CON (top) and LES (bottom) groups. Rate remapping was strong in pCA3 of control rats but nearly abolished in lesioned rats.

(C) Distributions of PVC across CA3 subregions in the task. Solid line, median; dashed lines, quartiles.

(D) Rate remapping induced ΔPVC (within-room PVC − between-room PVC). Dots, spatial bins. CON: pCA3, 0.497 (0.276–0.780); mCA3, 0.022 (0.002–0.073). LES: pCA3, 0.080 (0.047–0.120); mCA3, 0.038 (0.013–0.101); dCA3, 0.070 (0.018–0.165). rmANOVA, 400 bins, group, $F$ (1, 798) = 830.614, $P$ < 0.001, $\eta^2$ = 0.510; band, $F$ (2, 798) = 1 012.102, $P$ < 0.001, $\eta^2$ = 0.559; group × band, $F$ (2, 798) = 831.464, $P$ < 0.001, $\eta^2$ = 0.510.

(E) Stability loss (SL, difference in repeated-trial PVC between control and lesioned groups, left) and discrimination loss (DL, difference in repeated-trial − Δ room A vs. B PVC between groups, right) across CA3 bands. SL: pCA3, 0.033 (−0.006–0.094); mCA3, 0.060 (0.025–0.088). DL: pCA3, 0.413 (0.147–0.687); mCA3, −0.005 (−0.062–0.048). rmANOVA, 400 bins, group, $F$ (1, 798) = 392.620, $P$ < 0.001, $\eta^2$ = 0.330; band, $F$ (2, 798) = 760.360, $P$ < 0.001, $\eta^2$ = 0.488; group × band, $F$ (2, 798) = 567.835, $P$ < 0.001, $\eta^2$ = 0.416.

(F) SC distributions for each CA3 band.

(G) Rate remapping induced ΔSC (within-room SC – between-room SC). CON: pCA3, 0.184 (0.021–0.359); mCA3, −0.024 (−0.039–0.028). LES: pCA3, 0.011 (−0.013–0.057); mCA3, 0.028 (−0.001–0.056); dCA3, 0.028 (0.011–0.045). LMM: 138 cells, group, $F$ (1, 126) = 9.733, $P$ = 0.002; band, $F$ (2, 126) = 4.005, $P$ = 0.048; group × band, $F$ (2, 126) = 3.914, $P$ = 0.050; negligible between-animal variance.

(H) Correlation between CA3 recording position and mean two-room ΔSC (CON, 8 tetrodes; LES, 15 tetrodes). CON ΔSC vs. LES ΔSC, *t*-test, t = 3.449, $P$ = 0.010, Cohen's d = −2.038.

(I and J) RD analyses (same format as in F and G). CON: pCA3, 0.392 (0.020–0.599); mCA3, 0.262 (−0.002–0.643). LES: pCA3, 0.004 (−0.002–0.120); mCA3, 0 (−0.014–0.009); dCA3, 0.007 (−0.001–0.112). 139 cells, group, $F$ (1, 126) = 36.741, $P$ < 0.001; band, $F$ (2, 126) = 2.786, $P$ = 0.098; group × band, $F$ (2, 126) = 0.183, $P$ = 0.670; negligible between-animal variance.

(K) Correlation between CA3 recording position and mean two-room ΔRD (CON, 8 tetrodes; LES, 15 tetrodes). CON ΔRD vs. LES ΔRD, t = 6.434, $P$ < 0.001, Cohen's d = −3.639.

n.s., not significant; within-group: *, $P$ < 0.05; ***, $P$ < 0.001; between-group: $^{\#\#\#}$, $P$ < 0.001.

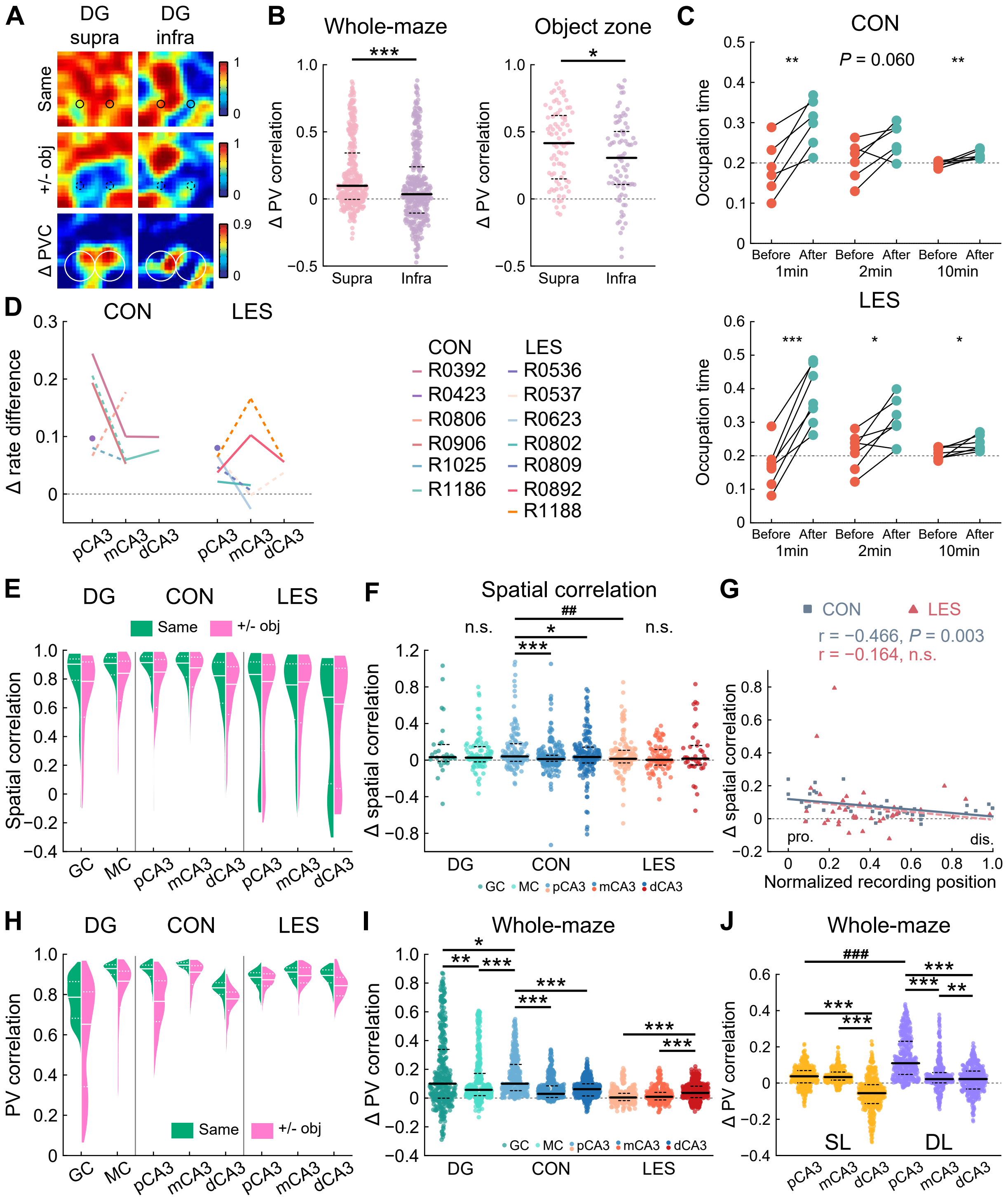

A
DG supra
DG infra
Same
+/- obj
Δ PVC
B
Whole-maze
Object zone
Δ PV correlation
Supra
Infra
C
CON
LES
Occupation time
P = 0.060
Before After
1min
2min
10min
D
CON
LES
Δ rate difference
pCA3 mCA3 dCA3
CON
R0392
R0423
R0806
R0906
R1025
R1186
LES
R0536
R0537
R0623
R0802
R0809
R0892
R1188
E
DG
CON
LES
Same
+/- obj
Spatial correlation
GC MC pCA3 mCA3 dCA3 pCA3 mCA3 dCA3
F
Spatial correlation
Δ spatial correlation
n.s.
##
GC MC pCA3 mCA3 dCA3
G
CON
LES
r = −0.466, P = 0.003
r = −0.164, n.s.
pro.
dis.
Normalized recording position
H
PV correlation
I
Whole-maze
Δ PV correlation
J
Whole-maze
###
SL
DL

**Figure S9. Extended analyses of the object-placement task.**

(A) PVC heatmaps for GCs located in supra- vs infrapyramidal blades.

(B) Left: whole-maze ΔPVC by blade. Supra, 139 cells, 0.098 (−0.003–0.343); Infra, 55 cells, 0.035 (−0.105–0.239); Wilcoxon signed-rank test, 400 bins, Z = 5.760, $P$ < 0.001, $\eta^2$ = 0.083. Right: object-zone ΔPVC by blade. Supra, 0.416 (0.150–0.622); Infra, 0.306 (0.108–0.503); 80 bins, Z = 2.034, $P$ = 0.042, $\eta^2$ = 0.052.

(C) Object-zone occupation over time. Both control and lesioned rats increased exploration after object placement. CON: 1 min, before, 0.187 ± 0.027, after, 0.300 ± 0.024, paired *t*-test, 6 rats, t = −4.915, $P$ = 0.004, Cohen's d = −2.007; 2 min, before, 0.204 ± 0.020, after, 0.257 ± 0.017, t = −2.426, $P$ = 0.060; 10 min, before, 0.196 ± 0.003, after, 0.221 ± 0.005, t = −4.091, $P$ = 0.009, Cohen's d = 1.670; LES: 1 min, before, 0.169 ± 0.025, after, 0.380 ± 0.033, 7 rats, t = −7.998, $P$ < 0.001, Cohen's d = 3.023; 2 min, before, 0.212 ± 0.021, after, 0.301 ± 0.026, t = −3.507, $P$ = 0.013, Cohen's d = 1.325; 10 min, before, 0.207 ± 0.007, after, 0.236 ± 0.009, t = −2.619, $P$ = 0.040, Cohen's d = 0.990.

(D) Object-induced ΔRD (repeated-trial RD – RD between object-present and object-absent trials) per rat. Colored lines and dots, animal means; dashed lines, < 20 place cells. Trends mirror pooled-cell analyses.

(E) SC distributions across DG and CA3 subregions.

(F) Object-induced ΔSC (repeated-trial SC – SC between object-present and object-absent trials). DG: GC, 0.032 (−0.015–0.170); MC, 0.027 (−0.019–0.147). CON: pCA3, 0.040 (−0.013–0.179); mCA3, 0.012 (−0.013–0.052); dCA3, 0.034 (−0.032–0.142). LES: pCA3, 0.014 (−0.031–0.108); mCA3, 0.003 (−0.054–0.116); dCA3, 0.016 (−0.053–0.160). GC vs. MC vs. pCA3, LMM: 257 cells; group, $F$ (2, 139.889) = 1.323, $P$ = 0.270; random effect of animal, $P$ = 0.430. CON vs. LES, LMM, 855 cells, $F$ (1, 4.157) = 0.766, $P$ = 0.429; band, $F$ (2, 287.232) = 6.973, $P$ = 0.001; group × band, $F$ (2, 287.232) = 3.671, $P$ = 0.027; random effect of animal, $P$ = 0.768.

(G) Correlation between CA3 recording position and mean object-induced ΔSC (CON, 39 tetrodes; LES, 39 tetrodes). The control and lesion groups showed similar ΔSC between repeated trials and object-present vs. object-absent trials (Wilcoxon rank-sum test, Z = 1.844, $P$ = 0.065). No significant difference in gradient slope was observed between groups (permutation test, $P$ = 0.456).

(H) PVC distributions across CA3 subregions.

(I) Object-induced whole-maze ΔPVC (repeated-trial PVC – PVC between object-present and object-absent trials). Dots, spatial bins. DG: GC, 0.100 (−0.001–0.338); MC, 0.057 (0.017–0.172). CON: pCA3, 0.100 (0.052–

0.234); mCA3, 0.029 (0.004–0.085); dCA3, 0.062 (0.015–0.100). LES: pCA3, 0.004 (−0.017–0.033); mCA3, 0.009 (−0.013–0.040); dCA3, 0.036 (0.003–0.083). GC vs. MC vs. pCA3, Friedman test, 400 bins, $\chi^2$ (2) = 37.980, $P < 0.001$, Kendall's W = 0.047. CON vs. LES, rmANOVA, 400 bins, group, $F$ (1, 1 197) = 645.692, $P < 0.001$, $\eta^2 = 0.350$; band, $F$ (2, 1 197) = 57.623, $P < 0.001$, $\eta^2 = 0.088$; group × band, $F$ (2, 1 197) = 205.121, $P < 0.001$, $\eta^2 = 0.255$.

(J) Stability loss (SL, difference in repeated-trial PVC between control and lesioned groups, left) and discrimination loss (DL, difference in repeated-trial − Δ with/without objects PVC between groups, right) across CA3 bands. SL, pCA3, 0.038 (0.001–0.069); mCA3, 0.033 (0.016–0.061); dCA3, −0.056 (−0.113–−0.009). DL, pCA3, 0.110 (0.048–0.230); mCA3, 0.022 (0.001–0.058); dCA3, 0.022 (−0.033–0.066). rmANOVA, 400 bins, Group, $F$ (1, 1 197) = 459.981, $P < 0.001$, $\eta^2 = 0.278$; band, $F$ (2, 1 197) = 303.146, $P < 0.001$, $\eta^2 = 0.336$; group × band, $F$ (2, 1 197) = 122.540, $P < 0.001$, $\eta^2 = 0.170$.

n.s., not significant; within-group: *, $P < 0.05$; **, $P < 0.01$; ***, $P < 0.001$; between-group: $^{\#\#}$, $P < 0.01$; $^{\#\#\#}$, $P < 0.001$.

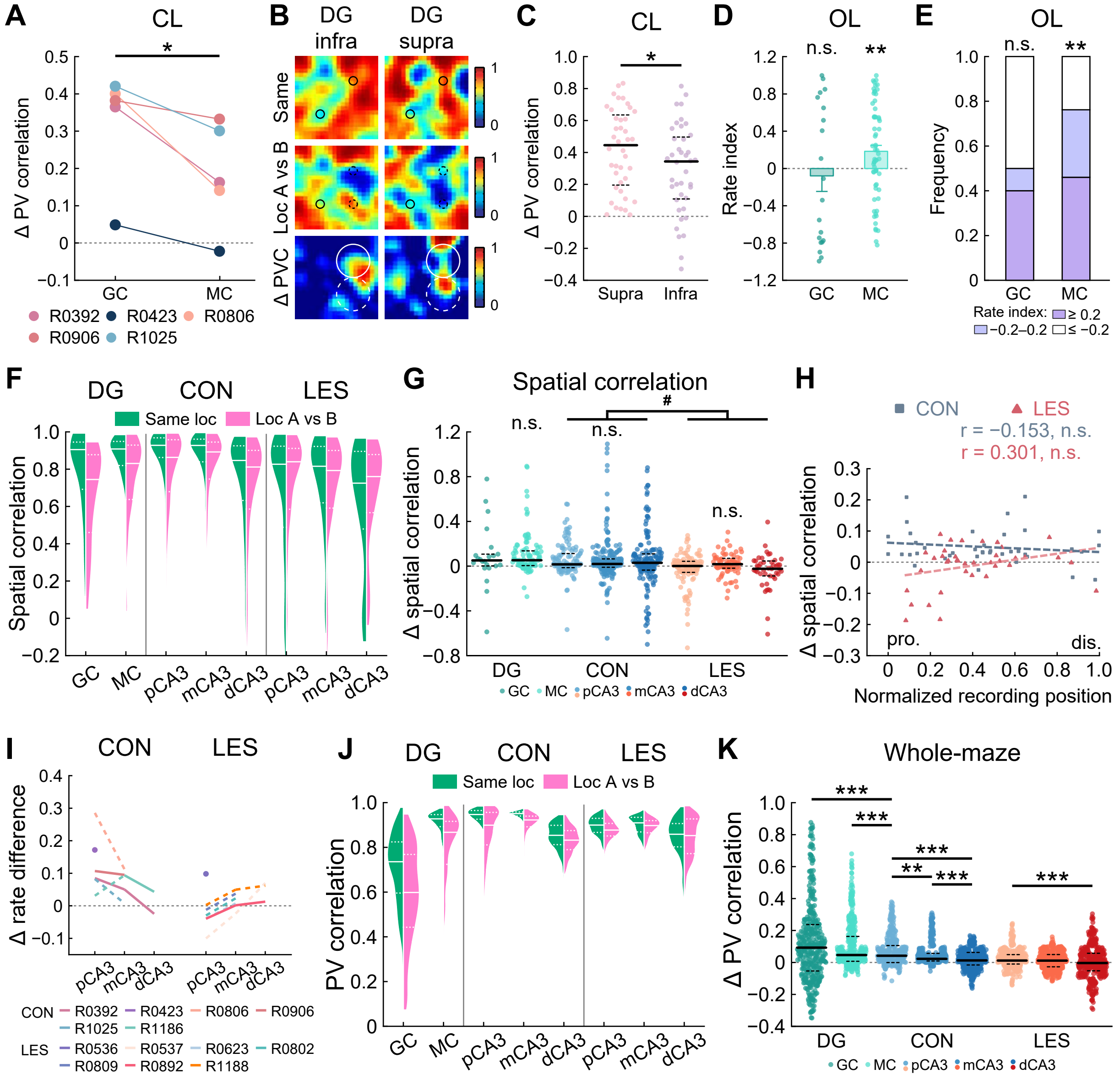

A
CL
Δ PV correlation
GC
MC
R0392 R0423 R0806
R0906 R1025
B
DG infra
DG supra
Same
Loc A vs B
Δ PVC
C
CL
Δ PV correlation
Supra
Infra
D
OL
Rate index
n.s.
**
GC
MC
E
OL
Frequency
n.s.
**
GC
MC
Rate index: ≥ 0.2
−0.2–0.2 ≤ −0.2
F
DG
CON
LES
Same loc
Loc A vs B
Spatial correlation
GC MC pCA3 mCA3 dCA3 pCA3 mCA3 dCA3
G
Spatial correlation
#
n.s.
n.s.
n.s.
Δ spatial correlation
DG CON LES
GC MC pCA3 mCA3 dCA3
H
CON
LES
r = −0.153, n.s.
r = 0.301, n.s.
Δ spatial correlation
pro.
dis.
Normalized recording position
I
CON
LES
Δ rate difference
pCA3 mCA3 dCA3
CON R0392 R0423 R0806 R0906
R1025 R1186
LES R0536 R0537 R0623 R0802
R0809 R0892 R1188
J
DG
CON
LES
Same loc
Loc A vs B
PV correlation
GC MC pCA3 mCA3 dCA3 pCA3 mCA3 dCA3
K
Whole-maze
Δ PV correlation
***
**
DG CON LES
GC MC pCA3 mCA3 dCA3

**Figure S10. Extended analyses of the object-relocation task.**

(A) CL-zone ΔPVC for simultaneously recorded GCs and MCs per rat (colored dots). GC, 0.382 (0.365–0.401); MC, 0.163 (0.141–0.301). GC vs. MC, Wilcoxon signed-rank test, 5 rats, $Z = 2.023$, $P = 0.043$, $\eta^2 = 0.819$.

(B) PVC heatmaps for supra- vs infrapyramidal GCs.

(C) CL-zone ΔPVC by blade. Supra, 123 cells, 0.445 (0.196–0.635); Infra, 46 cells, 0.343 (0.109–0.497); 40 bins, $Z = 2.231$, $P = 0.026$, $\eta^2 = 0.124$.

(D) Rate index (RI) at OL. Negative RI: reduced firing; positive RI, increased firing following object removal. GC, −0.079 ± 0.167, one sample *t*-test, 20 cells, $t = -0.471$, $P = 0.643$. MC, 0.184 ± 0.064, 63 cells, $t = 2.859$, $P = 0.006$, Cohen's $d = 0.360$. MCs showed a significant firing enhancement.

(E) Proportion of cells at OL, sorted by RI. RI > 0.2, rate increased; RI < −0.2, rate decreased; −0.2 < RI < 0.2, stable. Proportion of GCs: rate increased, 40.00%, stable, 10.00%, decreased, 50.00%; increased vs decreased, chi-square test, $\chi^2 = 0.404$, $P = 0.525$. MCs: increased, 46.03%, stable, 30.16%, decreased, 23.81%; $\chi^2 = 6.845$, $P = 0.009$, $\varphi = 0.233$.

(F and G) SC distributions (F), and relocation-induced ΔSC (repeated-trial SC – SC between pre- and post-relocation trials; G) across DG and CA3 subregions. DG: GC, 0.051 (0.002–0.107); MC, 0.053 (0.005–0.136). CON: pCA3, 0.016 (−0.013–0.112); mCA3, 0.020 (−0.010–0.065); dCA3, 0.030 (−0.036–0.110). LES: pCA3, 0.002 (−0.056–0.043); mCA3, 0.018 (−0.019–0.070); dCA3, −0.023 (−0.087–0.046). GC vs. MC vs. pCA3, LMM: 235 cells; group, $F(2, 265) = 2.290$, $P = 0.103$, negligible between-animal variance. CON vs. LES, LMM: 751 cells, group, $F(1, 10.741) = 8.709$, $P = 0.013$; band, $F(2, 462.626) = 1.287$, $P = 0.277$; group × band, $F(2, 462.626) = 1.304$, $P = 0.272$; random effect of animal, $P = 0.305$.

(H) Tetrode position versus CA3 ΔSC (CON, 39 tetrodes; LES, 38 tetrodes). The control group showed greater ΔSC between repeated trials and pre- versus post-relocation trials than the lesion group (Wilcoxon rank-sum test, $Z = 4.055$, $P < 0.001$, $\eta^2 = 0.214$). No significant transverse gradients were observed in either group, nor did gradient slopes differ between groups (permutation test, $P = 0.052$).

(I) Relocation-induced ΔRD (repeated-trial RD – RD between pre- and post-relocation trials) per rat. Colored lines and dots, animal means; dashed lines, < 20 place cells. Trends mirror cell-level comparisons.

(J) Violin plots showing PVC distributions for GCs, MCs, and CA3 neurons. Solid line, median; dashed lines, quartiles.

(K) Relocation-induced whole-maze ΔPVC (repeated-trial PVC – PVC between pre- and post-relocation trials). GC, 0.093 (−0.053–0.237); MC, 0.046 (0.007–0.163). CON: pCA3, 0.042 (−0.001–0.106); mCA3, 0.023 (0.009–0.057); dCA3, 0.013 (−0.016–0.063). LES: pCA3, 0.012 (−0.010–0.049); mCA3, 0.011 (−0.027–0.050); dCA3, −0.003 (−0.052–0.057). GC vs. MC vs. pCA3, Friedman test, 400 bins, $\chi^2$ (2) = 29.195, $P < 0.001$, Kendall's W = 0.036. CON vs. LES, rmANOVA, 400 bins, group: $F$ (1, 1 197) = 160.069, $P < 0.001$, $\eta^2 = 0.118$; band: $F$ (2, 1 197) = 34.458, $P < 0.001$, $\eta^2 = 0.054$; group × band: $F$ (2, 1 197) = 8.790, $P < 0.001$, $\eta^2 = 0.014$.

n.s., not significant; *, $P < 0.05$; **, $P < 0.01$; ***, $P < 0.001$; between-group: $^{\#}$, $P < 0.05$.

**Table S1. Detailed statistical tests and parameter values for Figures 1 to 5.**

| Fig. ID | Task and parameter | Method | Statistics | *P* value | Effect size | N number |
|---|---|---|---|---|---|---|
| Fig. 1E | Tetrode tip positions | Kolmogorov–Smirnov test | Z = 1.047 | $P = 0.223$ | - | 85 |
| Fig. 1G | Left, DG | *t*-test | $t = 46.573$ | $P < 0.001$ | Cohen's $d = 25.911$ | CON, 6; LES, 7 |
| | Left, CA3 | | $t = -0.491$ | $P = 0.633$ | - | |
| | Left, CA1 | | $t = 3.211$ | $P = 0.016$ | Cohen's $d = 1.656$ | |
| | Right, DG | | $t = 29.997$ | $P < 0.001$ | Cohen's $d = 16.689$ | |
| | Right, CA3 | | $t = 1.869$ | $P = 0.088$ | - | |
| | Right, CA1 | | $t = 9.791$ | $P < 0.001$ | Cohen's $d = 5.077$ | |
| Fig. 2B | Place-cell fraction, GC vs MC | chi-square test | $\chi^2 = 11.378$ | $P = 0.001$ | $\varphi = 0.432$ | Table S2 |
| | Place-cell fraction, CON | | $\chi^2 = 36.331$ | $P < 0.001$ | Cramer's $V = 0.230$ | |
| | Place-cell fraction, LES | | $\chi^2 = 3.377$ | $P = 0.185$ | - | |
| | Non-PC fraction, GC vs. MC | Fisher's exact test | - | $P = 0.429$ | - | |
| | Non-PC fraction, pCA3, CON vs. LES | | - | $P = 0.064$ | - | |
| | Non-PC fraction, mCA3, CON vs. LES | | - | $P = 0.347$ | - | |
| | Non-PC fraction, dCA3, CON vs. LES | | - | $P = 0.269$ | - | |
| Fig. 2C | Active in one room, GC vs MC | chi-square test | $\chi^2 = 26.189$ | $P < 0.001$ | $\varphi = 0.739$ | GC: 16; MC:3; CON p/m/d: 67/163/44; LES p/m/d:75/23/16 |
| | Active in one room, CON | | $\chi^2 = 47.791$ | $P < 0.001$ | Cramer's $V = 0.323$ | |
| | Active in one room, LES | | $\chi^2 = 2.745$ | $P = 0.253$ | - | |
| Fig. 2D | Between-room SC, GC vs. MC vs. pCA3 | Linear mixed-effects model | Intercept: $F(2, 109.054) = 0.010$ | $P = 0.990$ | - | Table S2 |
| | | | Animal | $P = 0.348$ | - | |
| | Between-room SC, CON vs. LES | | Group, $F(1, 6.738) = 0.825$ | $P = 0.395$ | - | |
| | | | Band, $F(2, 569.147) = 2.947$ | $P = 0.053$ | - | |
| | | | Interaction, $F(2, 569.147) = 3.662$ | $P = 0.026$ | - | |
| | | | Animal | $P = 0.152$ | - | |
| Main text | Fractions of globally remapped neurons, pCA3, CON vs. LES | chi-square test | $\chi^2 = 0.065$ | $P = 0.799$ | - | CON p/m/d: 11/39/30; LES p/m/d:15/7/5 |
| | Fractions of globally remapped neurons, mCA3, CON vs. LES | | $\chi^2 = 0.035$ | $P = 0.852$ | - | |
| | Fractions of globally remapped neurons, dCA3, CON vs. LES | | $\chi^2 = 0.353$ | $P = 0.553$ | - | |
| Fig. 2E | CA3 recording position and between-room SC | Wilcoxon rank-sum test | Z = −0.411 | $P = 0.681$ | - | CON, 31; LES, 23 |
| | CON slope vs. LES slope | Permutation test | - | $P = 0.018$ | - | |
| Fig. 2G | Between-room PVC, GC vs. MC vs. pCA3 | Friedman test | $\chi^2$ (2) = 33.740 | $P < 0.001$ | Kendall's $W = 0.042$ | 400 bins |
| | Between-room PVC, CON vs. LES | Repeated-measures two-way ANOVA | Group, $F(1, 1\,197) = 1.167$ | $P = 0.280$ | - | |
| | | | Band, $F(2, 1\,197) = 676.730$ | $P < 0.001$ | $\eta^2 = 0.531$ | |
| | | | Interaction, $F(2, 1\,197) = 81.728$ | $P < 0.001$ | $\eta^2 = 0.120$ | |
| Fig. 2H | Within-room SC, GC vs. MC vs. pCA3 | Linear mixed-effects model | Intercept: $F(2, 65.539) = 11.170$ | $P < 0.001$ | - | Table S2 |
| | | | Animal | $P = 0.554$ | - | |
| | Within-room SC, CON vs. LES | | Group, $F(1, 6.645) = 7.323$ | $P = 0.032$ | - | |
| | | | Band, $F(2, 543.564) = 13.801$ | $P < 0.001$ | - | |
| | | | Interaction, $F(2, 543.564) = 2.618$ | $P = 0.074$ | - | |
| | | | Animal | $P = 0.176$ | - | |
| Fig. 2I | CA3 recording position and within-room SC | Wilcoxon rank-sum test | Z = 3.332 | $P = 0.001$ | $\eta^2 = 0.206$ | CON, 31; LES, 23 |
| | CON slope vs. LES slope | Permutation test | - | $P = 0.173$ | - | |
| Fig. 2J | Within-room PVC, GC vs. MC vs. pCA3 | Friedman test | $\chi^2$ (2) = 302.195 | $P < 0.001$ | Kendall's $W = 0.378$ | 400 bins |
| | Within-room PVC, CON vs. LES | Repeated-measures two-way ANOVA | Group, $F(1, 1\,197) = 110.718$ | $P < 0.001$ | $\eta^2 = 0.085$ | |
| | | | Band, $F(2, 1\,197) = 1\,060.426$ | $P < 0.001$ | $\eta^2 = 0.639$ | |
| | | | Interaction, $F(2, 1\,197) = 373.477$ | $P < 0.001$ | $\eta^2 = 0.384$ | |

| Fig. ID | Task and parameter | Method | Statistics | *P* value | Effect size | N number |
|---|---|---|---|---|---|---|
| Fig. 2K | SL and DL across CA3 bands | Repeated-measures two-way ANOVA | Group, $F$(1, 1 197) = 22.377 | $P$ < 0.001 | $\eta^2$ = 0.018 | 400 bins |
| | | | Band, $F$(2, 1 197) = 369.883 | $P$ < 0.001 | $\eta^2$ = 0.382 | |
| | | | Interaction, $F$(2, 1 197) = 27.671 | $P$ < 0.001 | $\eta^2$ = 0.044 | |
| Main text | Navigational behavior, CON, 1 min | Paired $t$-test | $t$ = 0.675 | $P$ = 0.530 | - | 6 rats |
| | Navigational behavior, CON, 2 min | | $t$ = 0.934 | $P$ = 0.393 | - | |
| | Navigational behavior, CON, 10 min | | $t$ = 0.153 | $P$ = 0.884 | - | |
| | Navigational behavior, LES, 1 min | | $t$ = 2.341 | $P$ = 0.058 | - | 7 rats |
| | Navigational behavior, LES, 2 min | | $t$ = 1.232 | $P$ = 0.264 | - | |
| | Navigational behavior, LES, 10 min | | $t$ = 0.898 | $P$ = 0.404 | - | |
| Fig. 3B | Place-cell fraction, GC vs. MC | chi-square test | $\chi^2$ = 238.810 | $P$ < 0.001 | $\varphi$ = 0.744 | Table S2 |
| | Place-cell fraction, CON | | $\chi^2$ = 150.988 | $P$ < 0.001 | Cramer's $V$ = 0.316 | |
| | Place-cell fraction, LES | | $\chi^2$ = 1.024 | $P$ = 0.599 | - | |
| | Non-PC fraction, GC vs. MC | Fisher's exact test | - | $P$ = 1.000 | - | |
| | Non-PC fraction, pCA3, CON vs. LES | | - | $P$ = 0.014 | $\varphi$ = 0.073 | |
| | Non-PC fraction, mCA3, CON vs. LES | | - | $P$ = 0.029 | $\varphi$ = 0.080 | |
| | Non-PC fraction, dCA3, CON vs. LES | | - | $P$ = 0.336 | - | |
| Fig. 3D | Color-reversal ΔRD, GC vs. MC vs. pCA3 | Linear mixed-effects model | Intercept: $F$(2, 129.206) = 2.080 | $P$ = 0.129 | - | Table S2 |
| | | | Animal | $P$ = 0.695 | - | |
| | Color-reversal ΔRD, CON vs. LES | | Group, $F$(1, 1 313) = 14.886 | $P$ < 0.001 | - | |
| | | | Band, $F$(2, 1 313) = 12.096 | $P$ < 0.001 | - | |
| | | | Interaction, $F$(2, 1 313) = 7.558 | $P$ = 0.001 | - | |
| | | | Animal | - | - | |
| Fig. 3E | CA3 recording position and ΔRD | Wilcoxon rank-sum test | Z = 3.135 | $P$ = 0.002 | $\eta^2$ = 0.117 | CON, 39; LES, 45 |
| | CON slope vs. LES slope | Permutation test | - | $P$ = 0.049 | - | |
| Fig. 3F | Fractions of rate-remapping cells, GC vs. MC | chi-square test | $\chi^2$ = 3.035 | $P$ = 0.082 | - | Fig. 3F |
| | Fractions of rate-remapping cells, pCA3, CON vs. LES | | $\chi^2$ = 22.369 | $P$ < 0.001 | $\varphi$ = 0.212 | |
| | Fractions of rate-remapping cells, mCA3, CON vs. LES | | $\chi^2$ = 0.466 | $P$ = 0.495 | - | |
| | Fractions of rate-remapping cells, dCA3, CON vs. LES | | $\chi^2$ = 2.716 | $P$ = 0.099 | - | |
| Fig. 3I | Color-reversal ΔPVC for central bins, GC vs. MC vs. pCA3 | Friedman test | $\chi^2$ (2) = 25.541 | $P$ < 0.001 | Kendall's $W$ = 0.065 | 196 bins |
| | Color-reversal ΔPVC for central bins, CON vs. LES | Repeated-measures two-way ANOVA | Group, $F$(1, 585) = 213.515 | $P$ < 0.001 | $\eta^2$ = 0.267 | |
| | | | Band, $F$(2, 585) = 445.302 | $P$ < 0.001 | $\eta^2$ = 0.604 | |
| | | | Interaction, $F$(2, 585) = 115.433 | $P$ < 0.001 | $\eta^2$ = 0.283 | |
| Fig. 3J | SL and DL for central bins across CA3 bands | Repeated-measures two-way ANOVA | Group, $F$(1, 585) = 15.588 | $P$ < 0.001 | $\eta^2$ = 0.026 | 196 bins |
| | | | Band, $F$(2, 585) = 280.475 | $P$ < 0.001 | $\eta^2$ = 0.490 | |
| | | | Interaction, $F$(2, 585) = 111.949 | $P$ < 0.001 | $\eta^2$ = 0.277 | |
| Fig. 4D | Object-induced ΔRD, GC vs. MC vs. pCA3 | Linear mixed-effects model | Intercept: $F$(2, 57.748) = 1.712 | $P$ = 0.190 | - | Table S2 |
| | | | Animal | $P$ = 0.882 | - | |
| | Object-induced ΔRD, CON vs. LES | | Group, $F$(1, 9.778) = 7.020 | $P$ = 0.025 | - | |
| | | | Band, $F$(2, 486.849) = 3.582 | $P$ = 0.029 | - | |
| | | | Interaction, $F$(2, 486.849) = 3.497 | $P$ = 0.031 | - | |
| | | | Animal | $P$ = 0.641 | - | |
| Fig. 4E | CA3 recording position and ΔRD | $t$-test | $t$ = 2.019 | $P$ = 0.047 | Cohen's $d$ = 0.457 | CON, 39; LES, 39 |
| | CON slope vs. LES slope | Permutation test | - | $P$ = 0.044 | - | |

| Fig. ID | Task and parameter | Method | Statistics | *P* value | Effect size | N number |
|---|---|---|---|---|---|---|
| Fig. 4F | Relative DL | Friedman test | $\chi^2$ (2) = 71.120 | *P* < 0.001 | Kendall's *W* = 0.089 | 400 bins |
| Fig. 4H | Object zone ΔPVC, GC vs. MC vs. pCA3 | Friedman test | $\chi^2$ (2) = 21.775 | *P* < 0.001 | Kendall's *W* = 0.136 | 80 bins |
| | Object zone ΔPVC, CON vs. LES | Repeated-measures two-way ANOVA | Group: *F* (1, 237) = 447.848 | *P* < 0.001 | $\eta^2$ = 0.654 | |
| | | | Band: *F* (2, 237) = 62.995 | *P* < 0.001 | $\eta^2$ = 0.347 | |
| | | | Interaction, *F* (2, 237) = 130.384 | *P* < 0.001 | $\eta^2$ = 0.524 | |
| Fig. 4I | SL and DL across CA3 bands | Repeated-measures two-way ANOVA | Group, *F* (1, 237) = 580.188 | *P* < 0.001 | $\eta^2$ = 0.710 | 80 bins |
| | | | Band, *F* (2, 237) = 253.369 | *P* < 0.001 | $\eta^2$ = 0.681 | |
| | | | Interaction, *F* (2, 237) = 38.608 | *P* < 0.001 | $\eta^2$ = 0.246 | |
| Main text | Navigational behavior, CON, 1 min | Wilcoxon signed-rank test | Z = 0.734 | *P* = 0.463 | - | 6 rats |
| | Navigational behavior, CON, 2 min | Paired *t*-test | *t* = 0.424 | *P* = 0.689 | - | |
| | Navigational behavior, CON, 10 min | Paired *t*-test | *t* = 0.675 | *P* = 0.529 | - | |
| | Navigational behavior, LES, 1 min | Paired *t*-test | *t* = 1.235 | *P* = 0.263 | - | 7 rats |
| | Navigational behavior, LES, 2 min | Wilcoxon signed-rank test | Z = 1.183 | *P* = 0.237 | - | |
| | Navigational behavior, LES, 10 min | Paired *t*-test | *t* = 1.169 | *P* = 0.287 | - | |
| Fig. 5D | CL-zone (top) and OL-zone (bottom) ΔPVC (GC vs. MC vs. pCA3) | Two-way ANOVA | Band, *F* (2, 234) = 7.935 | *P* < 0.001 | $\eta^2$ = 0.064 | 240 bins |
| | | | Location, *F* (1, 234) = 1.827 | *P* = 0.178 | - | |
| | | | Interaction, *F* (2, 234) = 28.013 | *P* < 0.001 | $\eta^2$ = 0.193 | |
| | CL-zone (top) and OL-zone (bottom) ΔPVC between condition pairs | Three-way ANOVA (group, band, location) | Group, *F* (1, 468) = 99.639 | *P* < 0.001 | $\eta^2$ = 0.176 | 480 bins |
| | | | Band, *F* (2, 468) = 37.241 | *P* < 0.001 | $\eta^2$ = 0.137 | |
| | | | Location, *F* (1, 468) = 8.741 | *P* = 0.003 | $\eta^2$ = 0.018 | |
| | | | Interaction, *F* (2, 468) = 20.378 | *P* < 0.001 | $\eta^2$ = 0.080 | |
| Fig. 5F | Relocation-induced ΔRD, GC vs. MC vs. pCA3 | Linear mixed-effects model | Intercept: *F* (2, 99.404) = 2.893 | *P* = 0.060 | - | Table S2 |
| | | | Animal | *P* = 0.552 | - | |
| | Relocation-induced ΔRD, CON vs. LES | | Group, *F* (1, 7.449) = 4.181 | *P* = 0.078 | - | |
| | | | Band, *F* (2, 499.884) = 0.393 | *P* = 0.675 | - | |
| | | | Interaction, *F* (2, 499.884) = 5.986 | *P* = 0.003 | - | |
| | | | Animal | *P* = 0.258 | - | |
| Fig. 5G | CA3 recording position and ΔRD | *t*-test | *t* = 4.156 | *P* < 0.001 | Cohen's *d* = 0.935 | CON, 39; LES, 40 |
| | CON slope vs. LES slope | Permutation test | - | *P* = 0.003 | - | |
| Fig. 5H | CL vs. OL for discrimination loss | Two-way ANOVA | Band, *F* (2, 234) = 45.023 | *P* < 0.001 | $\eta^2$ = 0.278 | 240 bins |
| | | | Location, *F* (1, 234) = 1.679 | *P* = 0.196 | - | |
| | | | Interaction, *F* (2, 234) = 28.177 | *P* < 0.001 | $\eta^2$ = 0.194 | |
| | Stability loss vs. discrimination loss, CL | Repeated-measures two-way ANOVA | Group, *F* (1, 117) = 36.375 | *P* < 0.001 | $\eta^2$ = 0.237 | 40 bins |
| | | | Band, *F* (2, 117) = 99.988 | *P* < 0.001 | $\eta^2$ = 0.631 | |
| | | | Interaction, *F* (2, 117) = 51.518 | *P* < 0.001 | $\eta^2$ = 0.468 | |
| | Stability loss vs. discrimination loss, OL | Repeated-measures two-way ANOVA | Group, *F* (1, 117) = 21.637 | *P* < 0.001 | $\eta^2$ = 0.156 | 40 bins |
| | | | Band, *F* (2, 117) = 2.254 | *P* = 0.110 | - | |
| | | | Interaction, *F* (2, 117) = 3.371 | *P* = 0.038 | $\eta^2$ = 0.054 | |

**Table S2. Cell counts across behavioral tasks.**

Two-room task

| Group | CON | | | | | LES | | |
|---|---|---|---|---|---|---|---|---|
| Subtype/Region | GC | MC | pCA3 | mCA3 | dCA3 | pCA3 | mCA3 | dCA3 |
| Place cells | 19 | 29 | 87 | 245 | 126 | 108 | 41 | 27 |
| Non-PCs | 0 | 1 | 0 | 12 | 5 | 5 | 5 | 4 |
| Inactive cells | 21 | 0 | 80 | 111 | 19 | 80 | 44 | 27 |
| Total | 40 | 30 | 167 | 368 | 150 | 193 | 90 | 58 |

Color-reversal task

| Group | CON | | | | | LES | | |
|---|---|---|---|---|---|---|---|---|
| Subtype/Region | GC | MC | pCA3 | mCA3 | dCA3 | pCA3 | mCA3 | dCA3 |
| Place cells | 56 | 124 | 220 | 315 | 298 | 278 | 148 | 60 |
| Non-PCs | 4 | 1 | 9 | 8 | 9 | 24 | 12 | 5 |
| Inactive cells | 246 | 0 | 326 | 266 | 63 | 288 | 153 | 50 |
| Total | 306 | 125 | 555 | 589 | 370 | 590 | 313 | 115 |

Object-placement task

| Group | CON | | | | | LES | | |
|---|---|---|---|---|---|---|---|---|
| Subtype/Region | GC | MC | pCA3 | mCA3 | dCA3 | pCA3 | mCA3 | dCA3 |
| Place cells | 48 | 105 | 153 | 285 | 189 | 201 | 108 | 50 |
| Non-PCs | 7 | 0 | 2 | 13 | 5 | 10 | 10 | 2 |
| Inactive cells | 139 | 0 | 192 | 246 | 36 | 289 | 126 | 58 |
| Total | 194 | 105 | 347 | 544 | 230 | 500 | 244 | 110 |

Object-relocation task

| Group | CON | | | | | LES | | |
|---|---|---|---|---|---|---|---|---|
| Subtype/Region | GC | MC | pCA3 | mCA3 | dCA3 | pCA3 | mCA3 | dCA3 |
| Place cells | 39 | 98 | 131 | 272 | 171 | 139 | 80 | 49 |
| Non-PCs | 3 | 0 | 2 | 12 | 3 | 10 | 6 | 11 |
| Inactive cells | 127 | 0 | 161 | 239 | 37 | 223 | 108 | 63 |
| Total | 169 | 98 | 294 | 523 | 211 | 372 | 194 | 123 |

Two-room task (rate remapping)

| Group | CON | | | | | LES | | |
|---|---|---|---|---|---|---|---|---|
| Subtype/Region | GC | MC | pCA3 | mCA3 | dCA3 | pCA3 | mCA3 | dCA3 |
| Place cells | - | - | 35 | 7 | - | 53 | 35 | 9 |
| Non-PCs | - | - | 1 | 0 | - | 3 | 3 | 4 |
| Inactive cells | - | - | 41 | 9 | - | 43 | 36 | 11 |
| Total | - | - | 77 | 16 | - | 99 | 74 | 24 |

The table lists, for each subtype/region (granule cells, GC; mossy cells, MC; proximal CA3, pCA3; middle CA3, mCA3; and distal CA3, dCA3) and group (control, CON vs. lesion, LES), the number of cells classified as place cells, non-place cells (non-PCs), inactive cells, and the total recorded in each task. Note that non-place cells and inactive cells were excluded from spatial correlation and rate difference analyses.

**Table S3. Correlation between CA1 lesion extent and loss of CA3 discriminative ability.**

| | Δ SC | | | Δ RD | | | Δ PVC | | |
|---|---|---|---|---|---|---|---|---|---|
| | pCA3 | mCA3 | dCA3 | pCA3 | mCA3 | dCA3 | pCA3 | mCA3 | dCA3 |
| Two-room (within-room) | 0.148 | 0.092 | —— | 0.889 | 0.280 | —— | 0.127 | 0.263 | —— |
| Two-room (between-room) | 0.722 | 0.243 | —— | 0.853 | 0.249 | —— | 0.518 | 0.061 | —— |
| Color-reversal | 0.431 | 0.992 | 0.651 | 0.166 | 0.875 | 0.756 | 0.302 | 0.247 | 0.251 |
| Object-placement | 0.773 | 0.050 | 0.332 | 0.490 | 0.264 | <0.001 | 0.351 | 0.444 | 0.224 |
| Object-relocation | 0.203 | 0.078 | 0.428 | 0.465 | 0.067 | 0.648 | 0.494 | 0.955 | 0.446 |

The table presents *P*-values from correlation analyses examining whether the extent of CA1 damage is associated with CA3 deficits across five tasks and three spatial measures. The overall lack of significant correlations indicates that CA3 discriminative impairments are largely independent of CA1 lesion extent. Dashes indicate cases where data were not applicable or where the sample size was insufficient for correlation analysis (sample sizes: pCA3, n = 7; mCA3, n = 6; dCA3, n = 3; for the two-room task, n = 5, 4, and 2, respectively).

**Table S4. Rate remapping in control and lesioned rats.**

| Group | Animal | N cells | Stability | RD same color | RD color-reversal | Z value | *P* value |
|---|---|---|---|---|---|---|---|
| CON | R0392 | 223 | 0.742 | 0.190 | 0.214 | −3.592 | <0.001 |
| CON | R0423 | 109 | 0.874 | 0.133 | 0.195 | −3.569 | <0.001 |
| CON | R0806 | 74 | 0.802 | 0.191 | 0.261 | −3.456 | 0.001 |
| CON | R0906 | 166 | 0.840 | 0.178 | 0.206 | −5.605 | <0.001 |
| CON | R1025 | 96 | 0.776 | 0.135 | 0.170 | −2.677 | 0.007 |
| CON | R1186 | 345 | 0.820 | 0.161 | 0.172 | −3.951 | <0.001 |
| LES | R0536 | 42 | 0.602 | 0.332 | 0.333 | 0.569 | 0.569 |
| LES | R0537 | 69 | 0.681 | 0.297 | 0.282 | 0.236 | 0.813 |
| LES | R0623 | 87 | 0.667 | 0.190 | 0.213 | −1.274 | 0.203 |
| LES | R0802 | 80 | 0.686 | 0.145 | 0.149 | −2.155 | 0.031 |
| LES | R0809 | 34 | 0.831 | 0.176 | 0.225 | −1.796 | 0.073 |
| LES | R0892 | 78 | 0.777 | 0.155 | 0.195 | −1.739 | 0.082 |
| LES | R1188 | 96 | 0.792 | 0.151 | 0.139 | −1.352 | 0.176 |

This table summarizes the evaluation of rate remapping across individual animals. Rate remapping was defined by two criteria: (i) stable spatial firing, quantified as median spatial correlation > 0.5 across conditions, and (ii) a significant difference in firing rates between same-color and color-reversal conditions (Wilcoxon signed-rank test, $P < 0.05$). All control (CON) animals exhibited high spatial stability (median spatial correlation > 0.74) and significant firing rate differences under color-reversal conditions (all $P < 0.01$), consistent with robust rate remapping. In contrast, although lesioned (LES) animals maintained moderate spatial stability (median spatial correlation > 0.60), 6 of 7 animals failed to show significant firing rate differences, indicating impaired rate remapping following lesion.

**Table S5. Normalized changes in spatial metrics across CA3 subregions and experimental groups under non-spatial task conditions.**

| Color-reversal task | Spatial information content | Coherence | Stability |
|---|---|---|---|
| CON pCA3 | 0 (−0.002–0.037) | 0 (−0.015–0.019) | 0 (−0.012–0.015) |
| CON mCA3 | 0 (−0.005–0.015) | 0 (−0.014–0.005) | 0 (−0.007–0.014) |
| CON dCA3 | 0 (−0.010–0.010) | 0 (−0.016–0.012) | 0 (−0.018–0.034) |
| LES pCA3 | 0 (−0.005–0.014) | 0 (−0.010–0.033) | 0 (−0.005–0.032) |
| LES mCA3 | 0 (−0.003–0.014) | 0 (−0.011–0.034) | 0 (−0.013–0.012) |
| LES dCA3 | 0 (−0.008–0.014) | 0 (−0.002–0.028) | 0 (−0.017–0.005) |
| *P* value | Group: 0.231;<br>band: 0.247;<br>group × band: 0.941;<br>animal: 0.193 | Group: 0.689;<br>band: 0.639;<br>group × band: 0.649;<br>animal: —— | Group: 0.967;<br>band: 0.162;<br>group × band: 0.206;<br>animal: —— |

| Object-placement task | Spatial information content | Coherence | Stability |
|---|---|---|---|
| CON pCA3 | 0.012 (−0.026–0.119) | 0.009 (−0.033–0.126) | 0.020 (−0.020–0.144) |
| CON mCA3 | 0.012 (−0.028–0.059) | 0 (−0.032–0.041) | 0.002 (−0.042–0.044) |
| CON dCA3 | 0.018 (−0.031–0.102) | 0.014 (−0.042–0.098) | 0.021 (−0.037–0.258) |
| LES pCA3 | 0.001 (−0.023–0.061) | 0.005 (−0.027–0.096) | 0 (−0.066–0.086) |
| LES mCA3 | 0.002 (−0.035–0.064) | 0.001 (−0.049–0.082) | −0.006 (−0.064–0.062) |
| LES dCA3 | 0.009 (−0.033–0.137) | 0 (−0.038–0.190) | 0.011 (−0.025–0.396) |
| *P* value | Group: 0.770;<br>band: 0.280;<br>group × band: 0.138;<br>animal: —— | Group: 0.665;<br>band: 0.789;<br>group × band: 0.707;<br>animal: 0.282 | Group: 0.287;<br>band: 0.482;<br>group × band: 0.708;<br>animal: —— |

| Object-relocation task | Spatial information content | Coherence | Stability |
|---|---|---|---|
| CON pCA3 | 0.007 (−0.022–0.062) | 0.003 (−0.019–0.076) | 0.006 (−0.020–0.064) |
| CON mCA3 | 0.009 (−0.032–0.061) | 0.007 (−0.022–0.043) | 0.010 (−0.018–0.057) |
| CON dCA3 | 0.006 (−0.044–0.077) | 0 (−0.055–0.083) | 0 (−0.073–0.059) |
| LES pCA3 | 0.012 (−0.047–0.060) | 0 (−0.066–0.048) | 0.002 (−0.129–0.064) |
| LES mCA3 | 0.002 (−0.041–0.091) | 0 (−0.077–0.058) | 0.004 (−0.059–0.085) |
| LES dCA3 | 0 (−0.042–0.035) | −0.030 (−0.159–0.079) | −0.032 (−0.130–0.042) |
| *P* value | Group: 0.097;<br>band: 0.792;<br>group × band: 0.781;<br>animal: 0.814 | Group: 0.281;<br>band: 0.099;<br>group × band: 0.055;<br>animal: —— | Group: 0.292;<br>band: 0.416;<br>group × band: 0.852;<br>animal: 0.759 |

Note: values displayed as 0 fall within ±0.001 of zero (numerical integration tolerance); true zeros were not observed in this dataset. Dash indicates negligible between-animal variance.

**Table S6. Comparison of linear mixed-effects models with random intercept only versus random intercept plus random slopes.**

| Figure panel | Task and parameter | Slope | Group | Region | Interaction | Random intercept | Random slope |
|---|---|---|---|---|---|---|---|
| Fig. 2D | Two-room task between-room SC (GC vs. MC vs. pCA3) | Common | 0.990 | - | - | 0.348 | - |
| | | Random | 0.990 | - | - | 0.348 | - |
| Fig. 2D | Two-room task between-room SC (CON vs. LES) | Common | 0.395 | 0.053 | 0.026* | 0.152 | - |
| | | Random | 0.398 | 0.160 | 0.098* | 0.178 | 0.783 |
| Fig. 2H | Two-room task within-room SC (GC vs. MC vs. pCA3) | Common | <0.001* | - | - | 0.554 | - |
| | | Random | 0.127* | - | - | 0.576 | 0.251 |
| Fig. 2H | Two-room task within-room SC (CON vs. LES) | Common | 0.032 | <0.001 | 0.074 | 0.176 | - |
| | | Random | 0.032 | <0.001 | 0.074 | 0.176 | - |
| Fig. 3D | Color-reversal task ΔRD (GC vs. MC vs. pCA3) | Common | 0.129 | - | - | 0.695 | - |
| | | Random | 0.129 | - | - | 0.695 | - |
| Fig. 3D | Color-reversal task ΔRD (CON vs. LES) | Common | <0.001 | <0.001 | 0.001 | - | - |
| | | Random | 0.003 | 0.002 | 0.009 | - | 0.755 |
| Fig. 4E | Object-placement task ΔRD (GC vs. MC vs. pCA3) | Common | 0.190 | - | - | 0.882 | - |
| | | Random | 0.249 | - | - | - | 0.774 |
| Fig. 4E | Object-placement task ΔRD (CON vs. LES) | Common | 0.025 | 0.029 | 0.031 | 0.641 | - |
| | | Random | 0.025 | 0.029 | 0.031 | 0.641 | - |
| Fig. 5F | Object-relocation task ΔRD (GC vs. MC vs. pCA3) | Common | 0.060 | - | - | 0.552 | - |
| | | Random | 0.303 | - | - | 0.641 | 0.197 |
| Fig. 5F | Object-relocation task ΔRD (CON vs. LES) | Common | 0.078 | 0.675 | 0.003 | 0.258 | - |
| | | Random | 0.078 | 0.675 | 0.003 | 0.258 | - |
| Fig. S3F | Two-room task between-room RD (GC vs. MC vs. pCA3) | Common | <0.001 | - | - | - | - |
| | | Random | <0.001 | - | - | - | - |
| Fig. S3F | Two-room task between-room RD (CON vs. LES) | Common | 0.621 | <0.001 | 0.017* | 0.385 | - |
| | | Random | 0.616 | 0.004 | 0.085* | 0.516 | 0.633 |
| Fig. S3G | Two-room task within-room RD (GC vs. MC vs. pCA3) | Common | 0.562 | - | - | 0.670 | - |
| | | Random | 0.562 | - | - | 0.670 | - |
| Fig. S3G | Two-room task within-room RD (CON vs. LES) | Common | 0.307 | 0.001 | 0.898 | 0.353 | - |
| | | Random | 0.299 | 0.041 | 0.955 | 0.784 | 0.549 |
| Fig. S4B | Theta phase-locking | Common | 0.151 | <0.001 | 0.002* | 0.066 | - |
| | | Random | 0.155 | 0.010 | 0.090* | 0.078 | 0.697 |
| Fig. S4C | Theta phase precession | Common | 0.756 | 0.002 | 0.683 | 0.134 | - |
| | | Random | 0.663 | 0.032 | 0.819 | 0.308 | 0.624 |
| Fig. S4F | Mean firing rate | Common | 0.674 | <0.001 | 0.655 | 0.096 | - |
| | | Random | 0.670 | <0.001 | 0.688 | 0.103 | 0.941 |
| Fig. S4G | Peak firing rate | Common | 0.009 | <0.001 | 0.841 | 0.368 | - |
| | | Random | <0.001 | <0.001 | 0.652 | - | - |
| Fig. S4H | Place-field sizes | Common | 0.065 | <0.001 | 0.008* | 0.074 | - |
| | | Random | 0.066 | <0.001 | 0.123* | 0.137 | 0.513 |
| Fig. S4I | Number of place fields per cell | Common | 0.933 | 0.032 | 0.584 | 0.760 | - |
| | | Random | 0.992 | 0.076 | 0.643 | - | 0.662 |
| Fig. S4J | Spatial information content | Common | 0.001 | <0.001 | 0.001 | 0.074 | - |
| | | Random | 0.001 | <0.001 | 0.001 | 0.074 | - |

| Figure panel | Task and parameter | Slope | Group | Region | Interaction | Random intercept | Random slope |
|---|---|---|---|---|---|---|---|
| Fig. S4K | Spatial information rate | Common | 0.040 | <0.001 | 0.900 | 0.234 | - |
| | | Random | 0.067 | 0.008 | 0.819 | 0.522 | 0.424 |
| Fig. S4L | Coherence | Common | 0.002 | <0.001 | 0.049* | 0.079 | - |
| | | Random | 0.004 | 0.004 | 0.236* | 0.387 | 0.147 |
| Fig. S4M | Stability | Common | 0.021 | <0.001 | 0.005* | 0.043 | - |
| | | Random | 0.028 | 0.004 | 0.081* | 0.091 | 0.227 |
| Fig. S5B | Color-reversal task ΔSC (GC vs. MC vs. pCA3) | Common | 0.128 | - | - | 0.227 | - |
| | | Random | 0.361 | - | - | 0.389 | 0.526 |
| Fig. S5B | Color-reversal task ΔSC (CON vs. LES) | Common | 0.308 | 0.016 | 0.043* | 0.133 | - |
| | | Random | 0.307 | 0.108 | 0.153* | 0.173 | 0.920 |
| Fig. S7M | Color-reversal task ΔRD (downsampling) | Common | 0.031 | 0.083 | 0.022 | - | - |
| | | Random | 0.031 | 0.083 | 0.022 | - | - |
| Fig. S8G | Two-room task ΔSC (rate remapping) | Common | 0.002 | 0.048 | 0.050 | - | - |
| | | Random | 0.002 | 0.048 | 0.050 | - | - |
| Fig. S8J | Two-room task ΔRD (rate remapping) | Common | <0.001 | 0.098 | 0.670 | - | - |
| | | Random | <0.001 | 0.098 | 0.670 | - | - |
| Fig. S9F | Object-placement task ΔSC (GC vs. MC vs. pCA3) | Common | 0.270 | - | - | 0.430 | - |
| | | Random | 0.270 | - | - | 0.430 | - |
| Fig. S9F | Object-placement task ΔSC (CON vs. LES) | Common | 0.429 | 0.001 | 0.027* | 0.768 | - |
| | | Random | 0.309 | 0.009 | 0.084* | - | 0.667 |
| Fig. S10G | Object-relocation task ΔSC (GC vs. MC vs. pCA3) | Common | 0.103 | - | - | - | - |
| | | Random | 0.247 | - | - | - | 0.527 |
| Fig. S10G | Object-relocation task ΔSC (CON vs. LES) | Common | 0.013 | 0.277 | 0.272 | 0.305 | - |
| | | Random | 0.013 | 0.277 | 0.272 | 0.305 | - |
| Table S4 | Color-reversal task ΔSIc | Common | 0.231 | 0.247 | 0.941 | 0.193 | - |
| | | Random | 0.239 | 0.287 | 0.955 | 0.341 | 0.918 |
| Table S4 | Color-reversal task Δcoherence | Common | 0.689 | 0.639 | 0.649 | - | - |
| | | Random | 0.689 | 0.639 | 0.649 | - | - |
| Table S4 | Color-reversal task Δstability | Common | 0.967 | 0.162 | 0.206 | - | - |
| | | Random | 0.967 | 0.162 | 0.206 | - | - |
| Table S4 | Object-placement task ΔSIc | Common | 0.770 | 0.280 | 0.138 | - | - |
| | | Random | 0.743 | 0.415 | 0.299 | - | 0.432 |
| Table S4 | Object-placement task Δcoherence | Common | 0.665 | 0.789 | 0.707 | 0.282 | - |
| | | Random | 0.785 | 0.876 | 0.586 | - | 0.152 |
| Table S4 | Object-placement task Δstability | Common | 0.287 | 0.482 | 0.708 | - | - |
| | | Random | 0.287 | 0.482 | 0.708 | - | - |
| Table S4 | Object-relocation task ΔSIc | Common | 0.097 | 0.792 | 0.781 | 0.814 | - |
| | | Random | 0.097 | 0.792 | 0.781 | 0.814 | - |
| Table S4 | Object-relocation task Δcoherence | Common | 0.281 | 0.099 | 0.055 | - | - |
| | | Random | 0.281 | 0.099 | 0.055 | - | - |
| Table S4 | Object-relocation task Δstability | Common | 0.292 | 0.416 | 0.852 | 0.759 | - |
| | | Random | 0.265 | 0.425 | 0.825 | - | 0.557 |

Fixed-effects estimates and their significance patterns were highly consistent between the random-intercept-only and random-intercept-plus-slope models for the majority of comparisons. “-” indicates singularity (random slope variance estimated as zero). Results for all tested variables are shown; *: inconsistent results.

#### Supplementary Methods

##### Subjects

Data were obtained from 16 male Long-Evans rats (3–4 months, 400–500 g). Seven received bilateral dentate-gyrus (DG) lesions and tetrode implants targeting hippocampal CA3. Six sham-operated controls received identical implants. Three additional rats were given unilateral DG lesions for immunofluorescence. Animals were housed individually post-surgery under controlled temperature (20–23 °C) and humidity (40–60%) with a reversed 12 h light/dark cycle (lights off 9:00 a.m.). All behavioral testing took place during the dark phase. Rats had free access to water and were mildly food-restricted to 85–90% of *ad libitum* weight.

##### Approvals

All procedures were conducted at the Kunming Institute of Zoology, Chinese Academy of Sciences (Kunming, China), in accordance with national and institutional guidelines for the care and use of laboratory animals. The study was approved by the Institutional Animal Care and Use Committee (KIZ-IACUC-RE-2021-06-019), and every effort was made to minimize animal suffering and the number of animals used.

##### Electrode preparation

The multi-tetrode ‘‘hyperdrive’’, consisting of 18 independently movable tetrodes made of 17 μm polyimide-coated platinum-iridium (90:10%) wire (California Fine Wires, USA), was prepared as previously described (1). The electrode tips were electroplated to 150–250 kΩ at 1 kHz (Biomega, Bio-Signal Technologies, China) and gas-sterilized with ethylene oxide (Sanqiang Medical, China) within 48 hours before implantation.

##### Surgery

Anesthesia was induced and maintained with isoflurane gas (airflow: 0.8–1.0 L/min, 0.5–3% v/v in oxygen), with concentration adjusted in accordance with physiological monitoring parameters (RWD Life Science, China). Animals were placed in a stereotaxic apparatus (RWD Life Science, China) for colchicine infusion and electrode implantation. Pre-operative analgesia and infection prophylaxis consisted of subcutaneous meloxicam (2 mg/mL, 1 mg/kg body weight), enrofloxacin (50 mg/mL, 5 mg/kg), and atropine (0.5 mg/mL, 0.1 mg/kg). The scalp was infiltrated with 2 % lidocaine (200 μL, s.c.).

Colchicine (0.3–0.5 μg/μL; MEC, Cat# HY-16569, CAS 64-86-8) or phosphate-buffered saline (PBS) was infused at 0.5 μL/site into the dorsal DG of both hemispheres using a 5-μL Hamilton syringe at 10 μL/h. At each injection site, the needle was lowered 0.1 mm beyond the target, paused 1 min, retracted to injection depth, to create space for the infusion. After each injection, the needle was left in place at least 5 min to permit complete diffusion and prevent backflow along the needle track.

Infusion coordinates were:

1) Proximal and middle CA3 recordings: (1) anteroposterior (AP) 4.2 mm, mediolateral (ML) ±2.2 mm relative to bregma, dorsoventral (DV) 2.6 mm and 3.1 mm from dura; (2) AP 5.6 mm, ML ±2.6 mm relative to bregma, DV 2.8 mm from dura.

2) Distal CA3 recordings: (1) AP 3.9 mm, ML ±2.1 mm relative to bregma, DV 3.3 mm and 4.1 mm from dura; (2) AP 5.3 mm, ML ±2.3 mm relative to bregma, DV 3.9 mm from dura.

Immediately after infusions, the hyperdrive was implanted above the right hippocampus (AP 3.1–5.7 mm; ML 2.4–5.4 mm relative to bregma). Two cerebellar skull screws served as ground. The assembly was secured with stainless-steel screws and dental cement. Rats received soft diet and daily meloxicam and enrofloxacin for 5–7 days; they were monitored continuously until fully recovered. Unilateral-lesion animals received identical colchicine doses in the right DG and PBS in the left for immunofluorescence controls.

**Electrophysiological recordings**

The tetrodes were advanced into the CA1 pyramidal layer within three days after implantation, and subsequently in small daily increments until either the DG granule cell layer or the CA3/CA2 pyramidal layer was reached, using their distances from the CA1 cell layer as a reference when applicable. The cell layers were identified by the presence of high-amplitude complex-spike activity (2). Electrophysiological recording procedures followed previously described methods (3). Recordings began only after electrodes had remained stationary for at least 24 h to ensure signal stability. Repeated sampling from the same tetrode was accepted only if subsequent recording sites were separated by ≥ 40 μm, minimizing overlap between cell populations.

Neural signals were amplified, band-pass filtered (300 Hz–7.5 kHz) and digitized at 30 kHz with a Zeus multichannel data acquisition system (Bio-Signal Technologies, China). Spike waveforms exceeding a −50 μV threshold were time-stamped and recorded for 1 ms, referenced to a quiet cortical tetrode. LFPs were acquired simultaneously from one channel per tetrode (0.3–300 Hz band, 1 kHz sampling rate), using the two cerebellar skull screws as a reference. A 50 Hz notch filter was applied to eliminate interference from the mains frequency. Position was tracked at 50 Hz via head-mounted light-emitting diodes (Cyclops, Bio-Signal Technologies, China).

**Test environments**

Rats foraged for randomly scattered cookie crumbs in a fixed, dimly lit square arena (100 × 100 cm, < 10 lux). A single high-contrast cue card (50 × 30 cm) centered on one wall provided the only proximal visual landmark. For the color-reversal task, interchangeable color boards were separated from the animal by transparent plexiglass; flipping the boards altered wall color only, leaving odor and textural cues unchanged. Except during the two-room protocol, the arena was surrounded by ceiling-to-floor curtains (120 × 120 cm) that masked distal room cues. A resting box sat between the curtained arena and the experimenter. The two-room task used two separate rooms of comparable size but distinct spatial layouts, using identical arenas and curtains in both.

**Behavioral paradigms**

Behavioral paradigms were carried out as previously described (3). All protocols followed an A-B-B-A design (10 min per trial), in which A and B denoted distinct spatial or non-spatial conditions. Rats completed four trials per session with 5-min inter-trial rests outside the arena while the floor was cleaned with 75% ethanol. Animals were pre-trained repeatedly on the color-reversal task for 2–3 weeks until they consistently explored > 90% of the arena within 10 min. Once tetrodes reached the target layer, each rat was tested on five consecutive days in the following order: color-reversal, two-room, object-placement, and object-relocation tasks. Recording experiments were typically repeated 3–4 times per task, as tetrodes in the same animal rarely reached optimal sampling positions simultaneously.

Task details:

1) Color-reversal: Trials 1 & 4: standard black walls; Trials 2 & 3: walls switched to white; arena unchanged.

2) Two-room: Trials 1 & 4: Room A; Trials 2 & 3: Room B (identical arena geometry).

3) Object-placement: Trials 1 & 4: empty arena; Trials 2 & 3: two identical glass bottles placed 30 cm from the nearest walls in quadrants 3 and 4.

4) Object-relocation: Trials 1 & 4: familiar bottles in quadrants 3 and 4; Trials 2 & 3: quadrant-4 bottle moved to quadrant 1.

**Histology and recording sites**

After the final recording session, electrodes were left in place and rats were deeply anaesthetized and transcardially perfused with 0.9% saline followed by formalin. Brains were post-fixed for 24 h, cryoprotected in 30% sucrose, and sectioned coronally at 40 μm on a cryostat microtome (KD-2950, KEDEE, China). Sections were Nissl-stained with cresyl violet (Sigma Aldrich, USA, CAS: 10510-54-0, Cat# C5042-10G) and photographed under a bright-field microscope (UB103i, COIC; 2×/0.06 NA, 4×/0.13 NA objectives) with UOPView v2.0.

Hippocampal subregions were delineated according to established anatomical criteria (4). Because (a) tetrodes often advanced slightly during the post-experimental period before animals were sacrificed, and (b) post-fixation tissue shrinkage with tetrodes *in situ* typically caused track termini to appear deeper than their actual recording positions, tetrode positions were verified against physiological and anatomical landmarks. Tetrodes with tips located in or immediately below the granule cell layer were assigned to DG. CA3 positions were measured from the proximal pole, normalized to total CA3 length in ImageJ (v1.52a, RRID:SCR_003070), and classified as proximal (pCA3, 0–0.40), middle (mCA3, 0.40–0.70), and distal (dCA3, 0.70–1.0) portions. The dCA3 band may include CA2-contributed activity from tetrodes near the CA3/CA2 border.

**Lesion quantification**

Nissl-stained sections were digitized with a 10×/0.4 NA objective on an Olympus BX61 microscope (RRID:SCR_020343). To account for mossy-fiber spread along the longitudinal axis (5), every 40 μm section within ±500 μm of the recording site (12 sections per animal) was analyzed. The uninterrupted lengths of the DG granule-cell layer and the CA3/CA1 pyramidal layers were traced bilaterally in ImageJ and expressed as a percentage of length measured in age-matched control brain. Regions with only minor thinning (partially damaged) were excluded.

**Immunofluorescence**

Free-floating brain sections were blocked with 3% normal goat serum (NGS; Bytotime, China, Cat# C0265) in PBS for 1 hour at room temperature, then incubated overnight at 4 °C with primary antibodies diluted in PBS with 3% NGS and 0.5% Triton X-100. Mouse anti-calbindin D-28k (CB, 1:5,000, Swant, Cat# 300, RRID AB_10000347) and rabbit anti-Purkinje cell protein 4 (PCP4, 1:1,000, Sigma-Aldrich, Cat# HPA005792, RRID AB_1855086) were used to detect mossy cells, and mouse anti-parvalbumin (PV, 1:5,000, Swant, Cat# 235, RRID AB_10000343) was used for PV-expressing basket cells. After three PBS washes, sections were incubated for 2 h at room temperature with secondary antibodies (Cy3-conjugated goat anti-mouse IgG, 1:200, Proteintech, Cat# SA00009-1, RRID AB_2814746; CoraLite488-conjugated goat anti-rabbit IgG, 1:500, Proteintech, Cat# SA00013-2, RRID AB_2797132) diluted in PBS with 3% NGS and 0.5% Triton X-100, protected from light. Finally, sections were washed, mounted on gelatin-coated slides, and cover-slipped with anti-fade mounting medium.

Images were captured with a TissueFAXS PLUS confocal tissue cytometer (TissueGnostics, Austria) with a 10×/0.3 NA objective. Brightness and contrast were adjusted linearly and uniformly to the images under analysis. CB-negative, PCP4-positive neurons in the DG were counted as mossy cells; and PV-positive neurons adjacent to the dentate granule cell layer were counted as basket cells.

**Behavioral metrics**

Four metrics were employed to quantify the rat's navigational behavior in each experiment:

Speed: median instantaneous speed within a ±20 ms sliding window.

Tortuosity: median momentary tortuosity expressed by the ratio of path length to straight-line distance between trajectory ends within a ±1 s sliding window.

Coverage: proportion of 5 × 5 cm spatial bins with an occupation time > 0.5 s.

Evenness: mean global Moran's I index over occupation time in spatial bins (6):

$$I = \frac{N}{W}\frac{\sum_i \sum_j w_{ij}(x_i - \bar{x})(x_j - \bar{x})}{\sum_i (x_i - \bar{x})^2};$$

Where *N* is the number of spatial bins indexed by *i* and *j*, $w_{ij}$ is a matrix of spatial weights (second order queen's case), *W* is the sum of all $w_{ij}$, *x* is the time the rat spent in each spatial bin, $\bar{x}$ is the mean of *x*. Evenness = 0 indicates random distribution; Evenness = 1 indicates perfect clustering.

**Object preference and thigmotaxis**

Occupation time (OT): proportion of trial time spent within a 20-cm radius of an object.

Discrimination index (DI): $DI = \frac{T_1 - T_2}{T_1 + T_2};$

Where $T_1$ denotes the exploration time around the relocated or novel object, and $T_2$ denotes the exploration time around the unchanged object. Positive values indicate a stronger preference for the relocated or novel object.

The same metric was used to quantify thigmotaxis by substituting $T_1$ = time along the walls and $T_2$ = time in the arena center; positive values denote wall-following behavior.

**Spike sorting and cell classification**

Single units were isolated offline with MClust graphical cluster-cutting package (v4.4, A.D. Redish) in MATLAB (RRID:SCR_001622) as previously described (3). Manual clustering used two-dimensional projections of a multidimensional feature space that included peak amplitude, energy, and peak-to-valley values from the four tetrode channels. Waveform and autocorrelation functions were also used as additional reference criteria. Spike sorting was conducted across the entire recording dataset, and the resulting spike trains were subsequently divided into the four behavioral trials.

Putative excitatory neurons were selected using criteria that excluded interneurons and axonal fibers: spike width > 200 μs (peak-to-trough), mean firing rate < 5 Hz, and the presence of occasional burst discharges (7). An additional filter of spatial information content > 0.5 bits/spike was applied to exclude putative hilar interneurons (8). Putative CA3 interneurons were classified based on their spike width, firing rate and waveform. Units that failed any criterion or that were recorded repeatedly in the same task—defined by similar spatial firing locations and/or inter-spike interval distributions—were excluded. This resulted in the exclusion of approximately 25% (two-room task), 53% (color-reversal task), 39% (object-placement task), and 41% (object-relocation task) of DG cells, and correspondingly 43%, 58%, 36%, and 36% of CA3 neurons, respectively.

**Rate maps and place fields**

Epochs with low-speed movement (< 2.5 cm/s) or tracking artifacts (> 100 cm/s) were discarded. Cells firing ≥ 0.10 Hz (total spikes / trial duration) were retained. Spike counts were binned into 5 × 5 cm bins and divided by dwell time to produce raw rate maps, then smoothed with a Gaussian filter centered on each bin. A place field was a contiguous region of at least 225 $cm^2$ (≥ 9 bins) where the firing rate exceeded 20% of the cell's peak and the peak within the area was ≥ 1 Hz (3). Non-overlapping fields were counted for each cell.

**Dentate cell type classification**

DG principal neurons were classified as putative granule cells (GCs) or mossy cells (MCs) with a random-forest classifier identical to that validated previously (8-11). 300 decision trees were trained on boot-strapped samples; at each split three features were selected from four predictors: mean exploration firing rate, burstiness (fraction of inter-spike intervals < 6 ms), rest-to-navigation rate ratio, and peak in-box rate.

Training data comprised 70 electrophysiologically identified excitatory units recorded in the two-room task. Consistent with established functional signatures, GCs fire sparsely and usually in only one environment, whereas MCs are active in both (8, 11), therefore, cells with < 0.10 Hz mean rate or single-room activity were used as GC prototypes (n = 40), and cells active in both rooms as MC prototypes (n = 30). Out-of-bag (OOB) misclassification rate was 8.57 %.

Classification confidence was quantified as:

$$Confidence = \frac{V_{GC} - {}^{N}/_{2}}{{}^{N}/_{2}};$$

Where $V_{GC}$ is the number of GC votes across the 300 trees and $N$ = 300. Values approaching ±1 reflect maximal certainty; values near 0 indicate ambiguity.

**Spatial modulation**

The spatial information rate ($SI_r$, bits/s) and spatial information content ($SI_c$, bits/spike) (12):

$$SI_r = \sum_i p_{i\lambda_i} \log_2 \frac{\lambda_i}{\lambda}; \qquad SI_c = \sum_i p_i \frac{\lambda_i}{\lambda} \log_2 \frac{\lambda_i}{\lambda};$$

where $\lambda_i$ is the mean firing rate in that bin, $\lambda$ is the overall mean firing rate of the cell, and $p_i$ is the occupancy probability of the $i$ -th bin (occupancy time in the bin divided by total recording time). $SI_r$ was obtained by multiplying $SI_c$ by the mean firing rate.

Spatial coherence: Pearson correlation between each unsmoothed bin and the mean of its eight adjacent bins (first-order autocorrelation).

Spatial stability: Pearson correlation between smoothed rate maps from the first and second halves of the trial.

**Spatial correlation (SC)**

To assess whether a cell's firing pattern remained stable or reorganized following spatial or non-spatial changes, we computed the Pearson correlation between smoothed 5 × 5 cm rate maps from two trials. Pairs in which both maps fired < 0.10 Hz on average were discarded. In the two-room task, the map from the second room was rotated in 90° steps and the maximum correlation value was taken. Animals with mean SC values across all active CA3 pyramidal neurons higher than 0.5 were classified as exhibiting rate remapping (3).

**Rate difference (RD) and rate index (RI)**

RD was computed as absolute rate change between trials:

$$RD = \left|\frac{FR_1 - FR_2}{FR_1 + FR_2}\right|;$$

where $FR_1$ and $FR_2$ are the mean firing rates of the same cell in two different trials. RD = 1 indicates exclusive activity in one trial, whereas RD = 0 denotes equal activity between trials. Neurons with RD changes across same-color and reversed-color pairs > 0.2 (corresponding to mean + 2 SD in control dCA3, which showed minimal rate remapping) were considered rate-remapped to color reversal.

RI quantified rate modulation within a 20-cm region of interest (ROI) centered on the manipulated object:

$$RI = \frac{FR_2 - FR_1}{FR_1 + FR_2};$$

where $FR_1$ and $FR_2$ are the mean rates of the same cell within the ROI before and after manipulation. |RI| > 0.2 signified enhancement (positive) or suppression (negative).

**Population vector correlation (PVC)**

For each condition, rate maps of principal neurons were assembled into a 20 × 20 × z array (x, y, spatial bins; z, cell identity), including units firing < 0.10 Hz and lacking a defined place field (13). In the two-room task, maps were rotated in 90° steps and the orientation yielding maximal single-cell SC was selected; For each x-y bin, the vector of firing rates across the z-dimension constituted the composite population vector (PV) at that location. Pearson correlations between matched-bin PVs were computed across trial pairs.

**Stability loss and discrimination loss**

Stability loss (SL), reflecting the reduction in firing stability between repeated trials under identical conditions following DG lesion, was calculated as:

$$SL = repeated\text{-}trial\ PVC_{CON} - repeated\text{-}trial\ PVC_{LES};$$

Discrimination loss (DL), reflecting diminished sensitivity to room-switch in the two-room task following DG lesion, was calculated as:

$$DL\_A = cross\text{-}condition\ PVC_{LES} - cross\text{-}condition\ PVC_{CON};$$

For tasks conducted within the same room—where place field locations remained unchanged across conditions—discrimination was defined as:

$$Discrimination = repeated\text{-}trial\ PVC - cross\text{-}condition\ PVC;$$

Therefore, total DL was computed as:

$$DL\_B = Discrimination_{CON} - Discrimination_{LES} = SL + DL\_A;$$

Relative changes in PVC were calculated as:

$$Reduction\ in\ stability = \frac{SL}{median\ repeated\text{-}trial\ PVC};$$

$$Fracton\ of\ SL = \frac{SL}{median\ (DL\_A + SL)};$$

$$Reduction\ in\ discrimination = \frac{DL\_B}{median\ discrimination}.$$

**Permutation test**

This procedure was used to examine differences in CA3 proximo-distal gradients in control and lesioned groups. Tetrode data and recording positions from both groups were pooled and randomly resampled without replacement into two groups. For each iteration, the difference in regression slopes between the two groups was calculated. This process was repeated 10,000 times to generate a null distribution of slope differences. The observed slope difference was then compared against this distribution; the *P* value was defined as the proportion of permuted values exceeding the observed difference.

**LFP power analysis**

Time-frequency decomposition was performed using the continuous wavelet transform. Absolute power spectra (0–100 Hz) were estimated from consecutive 5-s windows using the multi-aper method implemented in Chronux analysis software (v2.12, http://chronux.org/) and Buzsáki lab toolbox (https://github.com/buzsakilab/buzcode), with time-bandwidth product = 3 and 5 tapers. Experiments with poor LFP quality were excluded. Aperiodic components were removed using established method (14).

**Theta phase locking**

Theta-phase locking was assessed by quantifying the coupling between CA1 unit activity and hippocampal theta oscillations. Instantaneous theta phases were derived from the filtered LFP (7–10 Hz) using the Hilbert transform (MATLAB Signal Processing Toolbox). Each spike was assigned the nearest theta phase value at its corresponding time point. These phase values were then binned into 10° intervals (n = 36 bins) spanning 0–360°.

The strength of phase locking was characterized by the mean vector length (MVL), calculated using Circular Statistics Toolbox (v1.21; https://github.com/circstat/circstat-matlab):

$$MVL = \left|\frac{1}{n}\sum_i a_i(\cos\theta_i + j\sin\theta_i)\right|;$$

where *j* represents the imaginary unit, *n* is the number of phase bins (36), and $\theta_i$ and $a_i$ denote the theta phase and normalized response (spike count) of the *i*-th bin, respectively.

**Theta phase precession**

The 2D open arena is suboptimal for phase precession analysis because rats can enter place fields from multiple directions. To estimate the degree of phase precession, we computed the difference between intrinsic theta frequency and local theta frequency, adapting methods established in bats (15). When a neuron exhibits phase precession, its spikes shift to progressively earlier phases of the LFP theta oscillation, manifesting as a higher frequency in the spike train autocorrelogram. Local theta frequency (*Fl*) was defined as the peak frequency within theta band in the LFP power spectrum. Intrinsic theta frequency (*Fi*) for each neuron was estimated from the first peak in the spike train autocorrelogram computed with 10 ms bins. Theta phase precession was then calculated as:

$$Phase\ precession = F_i - F_l.$$

**Statistics and reproducibility**

Results are reported as mean ± standard error of the mean (SEM), or medians and interquartile ranges (IQR), depending on normality. Linear mixed-effects models (LMMs) were used to examine the fixed effects of group (lesion vs. control), CA3 band (pCA3, mCA3, dCA3), and their interaction on ΔSC, ΔRD, and spatial coding metrics, with random intercepts for neurons nested within rats. LMMs with random slopes were evaluated but not used (Table S6), as they may be unreliable due to unbalanced cell sampling across animals and CA3 bands (16). To account for repeated measurements of spatial bins across CA3 bands and groups, PVCs were analyzed with repeated-measures two-way analysis of variance (rmANOVA) with both bands and groups treated as within-subject factors. Variability of the PVC measure originates from spatial bins. Significant interactions were followed by Kruskal–Wallis (independent) or Friedman (related) tests. Pairwise comparisons used paired or unpaired *t*-tests, Wilcoxon rank-sum or signed-rank tests as appropriate. Frequency data were compared using Pearson's chi-square or Fisher's exact tests; linear relationships were assessed with Pearson correlation coefficients. Holm-Bonferroni *post hoc* tests were used to identify significant differences between pairs. Multiple comparisons between frequency bands were corrected using false discovery rate (FDR). In all cases, the cell (or the spatial bin, for PVC, or the tetrode, for LFP) was the unit of statistical analysis, not individual session-pair comparisons. All tests were two-tailed, and statistical significance was set at $P < 0.05$. Analyses were performed in SPSS v25 (IBM, RRID:SCR_002865), and plots were created in Origin 2024 (OriginLab Inc., Northampton, MA, USA).

**Data and code availability**

All data supporting the findings of this study, if not available in the Supplementary Data file, are available from the Lead Contact upon request. All algorithms and tools used in the analyses have been listed in the resources table and are available online.

**Supplementary References**


1. L. Lu, B. Popeney, J. D. Dickman, D. E. Angelaki, Construction of an Improved Multi-Tetrode Hyperdrive for Large-Scale Neural Recording in Behaving Rats. *J Vis Exp* 10.3791/57388 (2018).
2. L. Lu *et al.*, Impaired hippocampal rate coding after lesions of the lateral entorhinal cortex. *Nat Neurosci* **16**, 1085–1093 (2013).
3. Y. L. Duan *et al.*, Object-translocation induces event coding in the rat hippocampus. *Commun Biol* **8**, 797 (2025).
4. N. L. Cappaert, N. M. van Strien, M. P. Witter, "Hippocampal Formation" in The Rat Nervous System, G. Paxinos, Ed. (Academic Press, 2015), https://doi.org/10.1016/B978-0-12-374245-2.00020-6 chap. 20, pp. 511–573.
5. L. Acsády, A. Kamondi, A. Sík, T. Freund, G. Buzsáki, GABAergic cells are the major postsynaptic targets of mossy fibers in the rat hippocampus. *J Neurosci* **18**, 3386–3403 (1998).
6. P. A. P. Moran, The Interpretation of Statistical Maps. *Journal of the Royal Statistical Society: Series B (Methodological)* **10**, 243–251 (1948).
7. L. Lu, K. M. Igarashi, M. P. Witter, E. I. Moser, M. B. Moser, Topography of place maps along the CA3-to-CA2 axis of the hippocampus. *Neuron* **87**, 1078–1092 (2015).
8. D. GoodSmith *et al.*, Spatial representations of granule cells and mossy cells of the dentate gyrus. *Neuron* **93**, 677–690 e675 (2017).
9. D. GoodSmith, H. Lee, J. P. Neunuebel, H. Song, J. J. Knierim, Dentate gyrus mossy cells share a role in pattern separation with dentate granule cells and proximal CA3 pyramidal cells. *J Neurosci* **39**, 9570–9584 (2019).
10. D. GoodSmith *et al.*, Flexible encoding of objects and space in single cells of the dentate gyrus. *Curr Biol* **32**, 1088–1101.e1085 (2022).
11. S. H. Kim *et al.*, Global remapping in granule cells and mossy cells of the mouse dentate gyrus. *Cell Rep* **42**, 112334 (2023).
12. W. E. Skaggs, B. L. McNaughton, M. A. Wilson, C. A. Barnes, Theta phase precession in hippocampal neuronal populations and the compression of temporal sequences. *Hippocampus* **6**, 149–172 (1996).

13. S. Leutgeb *et al.*, Independent codes for spatial and episodic memory in hippocampal neuronal ensembles. *Science* **309**, 619–623 (2005).
14. T. Donoghue *et al.*, Parameterizing neural power spectra into periodic and aperiodic components. *Nat Neurosci* **23**, 1655–1665 (2020).
15. T. Eliav *et al.*, Nonoscillatory Phase Coding and Synchronization in the Bat Hippocampal Formation. *Cell* **175**, 1119–1130 e1115 (2018).
16. X. A. Harrison *et al.*, A brief introduction to mixed effects modelling and multi-model inference in ecology. *PeerJ* **6**, e4794 (2018).
